# CelltypeR: A flow cytometry pipeline to annotate, characterize and isolate single cells from brain organoids

**DOI:** 10.1101/2022.11.11.516066

**Authors:** Rhalena A. Thomas, Julien Sirois, Shuming Li, Alexandre Gestin, Ghislaine Deyab, Valerio E. Piscopo, Paula Lépine, Meghna Mathur, Carol X.Q. Chen, Vincent Soubannier, Taylor M. Goldsmith, Lama Fawaz, Thomas M. Durcan, Edward A. Fon

## Abstract

Motivated by the growing number of single cell RNA sequencing datasets (scRNAseq) revealing the cellular heterogeneity in complex tissues, particularly in brain and induced pluripotent stem cell (iPSC)-derived brain models, we developed a high-throughput, standardized approach for reproducibly characterizing cell types in complex neuronal tissues based on protein expression levels. Our approach combines a flow cytometry (FC) antibody panel targeting brain cells with a computational pipeline called CelltypeR, with functions for aligning and transforming datasets, optimizing unsupervised clustering, annotating and quantifying cell types, and statistical comparisons. We applied this workflow to human iPSC-derived midbrain organoids and identified the expected brain cell types, including neurons, astrocytes, radial glia, and oligodendrocytes. Defining gates based on the expression levels of our protein markers, we performed Fluorescence-Activated Cell Sorting of astrocytes, radial glia, and neurons, cell types were then confirmed by scRNAseq. Among the sorted neurons, we identified three subgroups of dopamine (DA) neurons; one reminiscent of substantia nigra DA neurons, the cell type most vulnerable in Parkinson’s disease. Finally, we use our workflow to track cell types across a time course of organoid differentiation. Overall, our adaptable analysis framework provides a generalizable method for reproducibly identifying cell types across FC datasets.

## Introduction

Investigating the molecular, cellular, and tissue properties of the human brain requires the use of cellular models, as live human brain tissue cannot be easily accessed for research. Patient-derived disease 3D tissues, such as human midbrain organoids (hMOs), derived from human induced pluripotent stem cells (iPSCs), provide a promising physiologically relevant model for human brain development and diseases, including neurodegenerative diseases such as Parkinson’s disease (PD).^1–4^ Yet, as new models emerge, the complexity and reproducibility of these systems needs to be captured to utilize this models in addressing biological questions. To determine how faithfully organoids recapitulate the human brain and how organoids derived from individuals with disease differ from those derived from healthy controls, new approaches towards characterization are required. Effective and quantitative methods are needed to determine the cell types within these complex tissues and to apply these benchmarks reproducibly across experiments. At present, individual cells within brain or organoid tissue can be identified using single cell RNA sequencing (scRNAseq) or labeling of protein or RNA in tissue sections. These tools are useful but limited. scRNAseq is a powerful tool that has been used to identify known and novel cell types, cell states, and cell fate trajectories^5–7^. However, using scRNAseq to compare proportions or populations of cells between genotypes over multiple time points is not practical for hMOs and may result in sampling bias, as less than 1% of the whole tissue is sequenced. While scRNAseq provides detailed expression values to determine sub-types of cells, only relatively few samples can be run at a given time and all the cells must be alive and prepared in parallel, which can lead to technically challenging experiments. These experiments are also costly for the number of replicates needed to ensure sufficient power for comparing multiple time points, disease states, or pharmacological treatments^8–10^. Another option to quantify cell types is immunostaining or *in situ* hybridization of tissue sections. This has the advantage of capturing cell morphology and spatial resolution. However, sample preparation, image acquisition, and analysis are labour intensive and limited in quantitative accuracy. Moreover, for 3D tissues, either only a small section can be analyzed or the entire tissue must be reconstructed and only a few cell types can be detected at once.^11,12^

Here, we use flow cytometry (FC) to measure the protein expression levels of a panel of cell surface markers enriched in specific brain cell types. FC is a fast, quantitative, and robust method, used widely in immunology and cancer research^13–15^ but to date only sparsely in neuroscience. Typically in neurobiology, only two or three antibodies are used to distinguish between pairs of cell types^16,17^ or to enrich one cell type.^18,19^ Traditional FC analysis methods using commercially available analysis software packages, such as FlowJo (Becton-Dickinson Biosciences) are time consuming and subject to user error. Methods to standardize data preprocessing and analyze combinations of more than 3 antibodies in one experiment are starting to emerge.^20,21^ However, no methods are available to streamline cell type annotation in FC from complex tissues such as brain or 3D brain organoids using a large antibody panel. To create such an analysis framework, we produced an experimental dataset using cultured hMOs differentiated from human iPSCs.^1,4,22^ Our workflow also provides the methods to select subtypes of cells and gate these cells for further analysis, such as RNAseq, proteomics, or enriching cultures. We select example cell populations, sort these cell types, and further characterize these with scRNAseq. Here we present a complete framework for annotating cell types within complex tissue and comparing proportions of cell types across conditions and experiments.

## Results

### An antibody panel to identify multiple cell types in human midbrain organoids

In **Figure 1A**, we provide a schematic of the CelltypeR analysis workflow (see methods) used to quantify and compare cell types from tissues containing a heterogeneous population of cells with a particular focus on neuronal tissue through brain organoids. To test our CelltypeR pipeline, we used hMOs^4^ differentiated from iPSC lines derived from three unrelated healthy individuals (**Methods Table 1**). The hMOs were grown for 9 months in culture, a time point at which neurons are expected to be mature and astrocytes and oligodendrocytes have been shown to be present.^1,23^ Immunofluorescence staining of cryosections show that these organoids contain neurons, astrocytes and oligodendrocytes (**Figure 1B and S1**). In FC, combinations of the relative intensities of 2-3 antibodies are often used to distinguish between cell types. However, in hMOs we expect approximately nine cellular types with a continuum of stages of differentiation.^1,4,24^ We first defined a panel of 13 antibodies, which included well-characterized antibodies previously used to define neural stem cells, neurons, astrocytes, and oligodendrocytes or to define other cell types in cultured immortalized human cell lines, blood, or brain tissues (**Table 1**). We dissociated the mature hMOs and labeled the cell suspension with these antibodies then measured the fluorescence intensity values corresponding to the protein targets using FC. The single live cells were sequentially gated using FlowJo. The FC results show that each protein has a range of expression across different cells (**Figure 1C and S2**). We conclude that the antibody panel has the potential to define cell types by identifying combinations of protein expression profiles unique to different cell groups.

**Figure 1:**
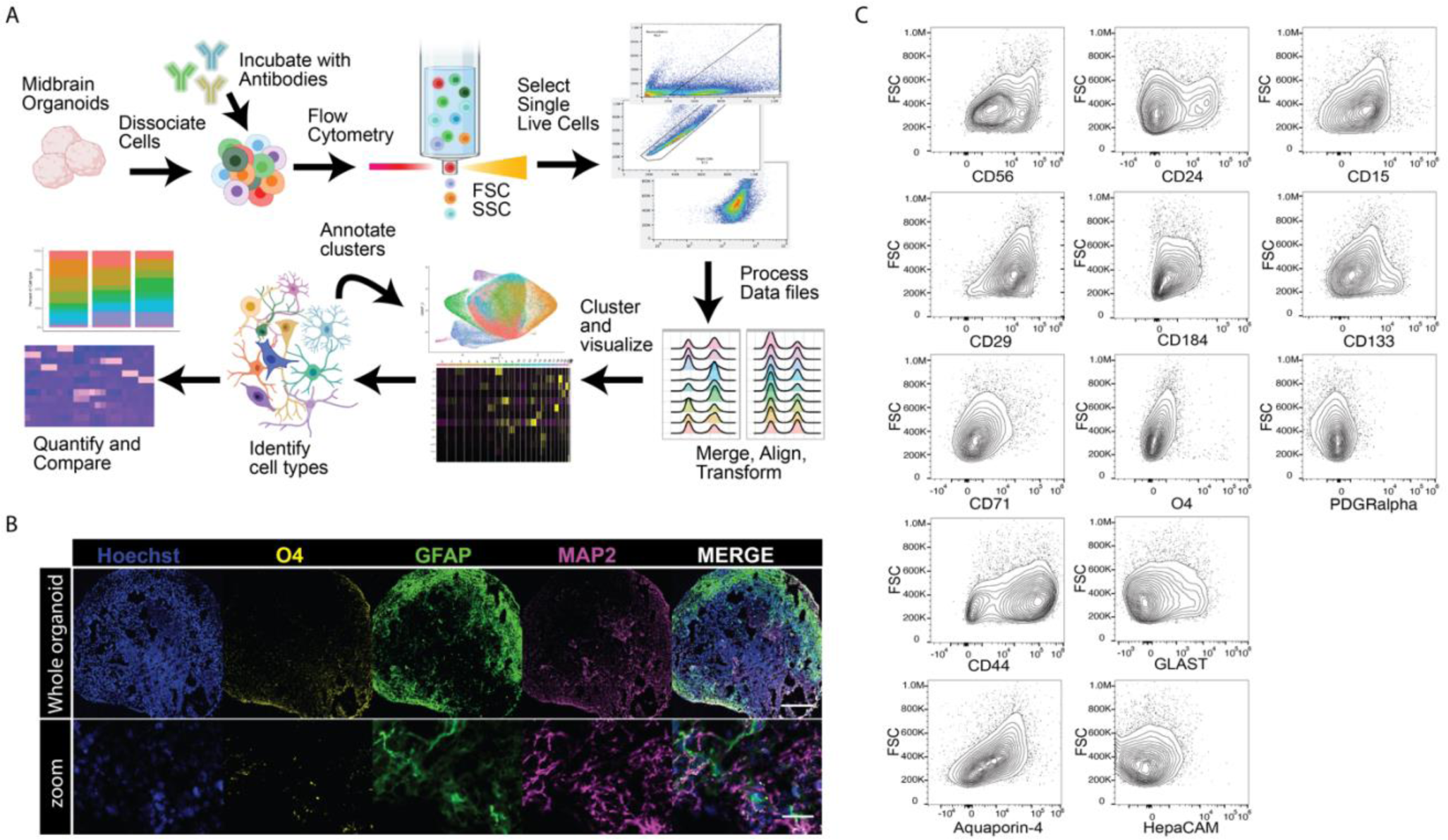
A workflow to identify and quantify cell types in midbrain organoids (hMOs) using a panel of flow cytometry antibodies. **A)** Schematic of the CelltypeR workflow: tissue (hMO) is dissociated and labelled with an antibody panel, expression levels are measured on individual cells using flow cytometry (FC), and live single cells are gated from the debris and doublets in FlowJo. The data is then preprocessed in R, merging files and harmonizing the data if desired. Unsupervised clustering is used to find groups of cell types, methods are provided to aid in cluster annotation, annotated cells are quantified, and statistical analyses are applied. **B)** Example image of a cryosection from an AJG001-C4C hMO, 285 days in final differentiation culture, showing total nuclei (Hoechst), oligodendrocytes (O4), astrocytes (GFAP), and neurons (MAP2). Top: cross section of a whole hMO stitched together from tiled images, scale bar = 250mm. Bottom: zoomed in image cropped from the whole hMO image, scale bar = 250mm. **C)** Contour plots showing the cell size on the y-axis (FSC) and intensity of staining for each antibody in the panel on the x-axis (log scale biexponential transformation).

**Table 1:**
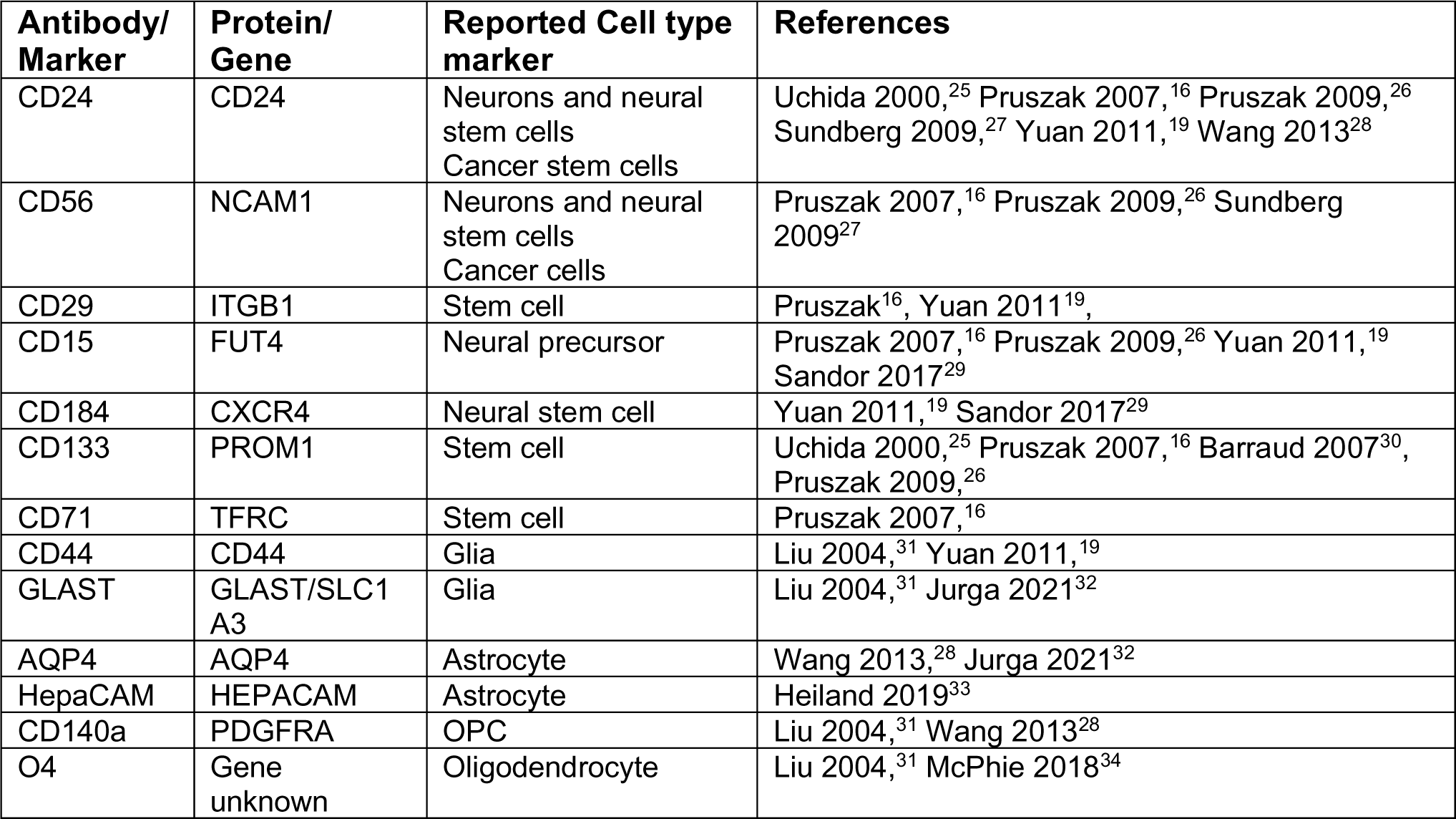
Antibody panel with cell types previously reported to be identified by each marker.

### Analysis of expression profiles in 2D cultures

To validate the expression of the proteins targeted by the selected antibodies on known cell types, we separately differentiated iPSCs into dopaminergic neuronal precursor cells (DA NPCs),^22^ dopaminergic neurons (DA neurons),^35^ astrocytes,^36^ and oligodendrocytes^34^ (**Figure 2A**). The cultures were dissociated and the 13 antibodies in the FC panel were applied. We examined the staining for each antibody across the cultured 2D cells (**Figure 2B**). Within each cell type there was a variation in protein expression levels that could be used to define subgroups of cells. To identify subgroups of cells and visualize the markers, we applied unsupervised clustering developed as part of the CelltypeR workflow. Some tools exist for automated processing and formatting of FC and numerous tools exist for cluster analysis of single cell transcriptomic data that can be applied to other FC data. Thus, we took advantage of these existing tools and created new functions in an R package to process FC data (see methods). We combined the FC acquired protein expression levels from the five separate iPSC derived cultures, normalized the data, and performed dimensional reductions. The UMAP visualization shows separate groups for each of the five cell types with some overlap (**Figure 2C**). The iPSCs are separate from all other cell cultures. Whereas the NPC culture splits into separate groups and overlaps with different cell types, the same is observed for the oligodendrocyte culture (**Figure S3A**). Clustering analysis identifies subgroups of cell types and some clusters with cells from multiple 2D cell cultures (**Figure 2D and S3B**). The DA NPC culture is an intermediate stage between iPSC and the three other cell cultures; therefore, it is not surprising that the cells from the NPC culture cluster together with other cell cultures. In the oligodendrocyte culture there is one cluster with the highest O4 expression that represents the oligodendrocytes within the culture (**Figure 2E and S4**). We conclude from these findings using iPSC-derived 2D cultures that our antibody panel can distinguish different cell types and subgroups of cells that we expect to find in 3D hMOs and other complex neuronal tissues.

**Figure 2:**
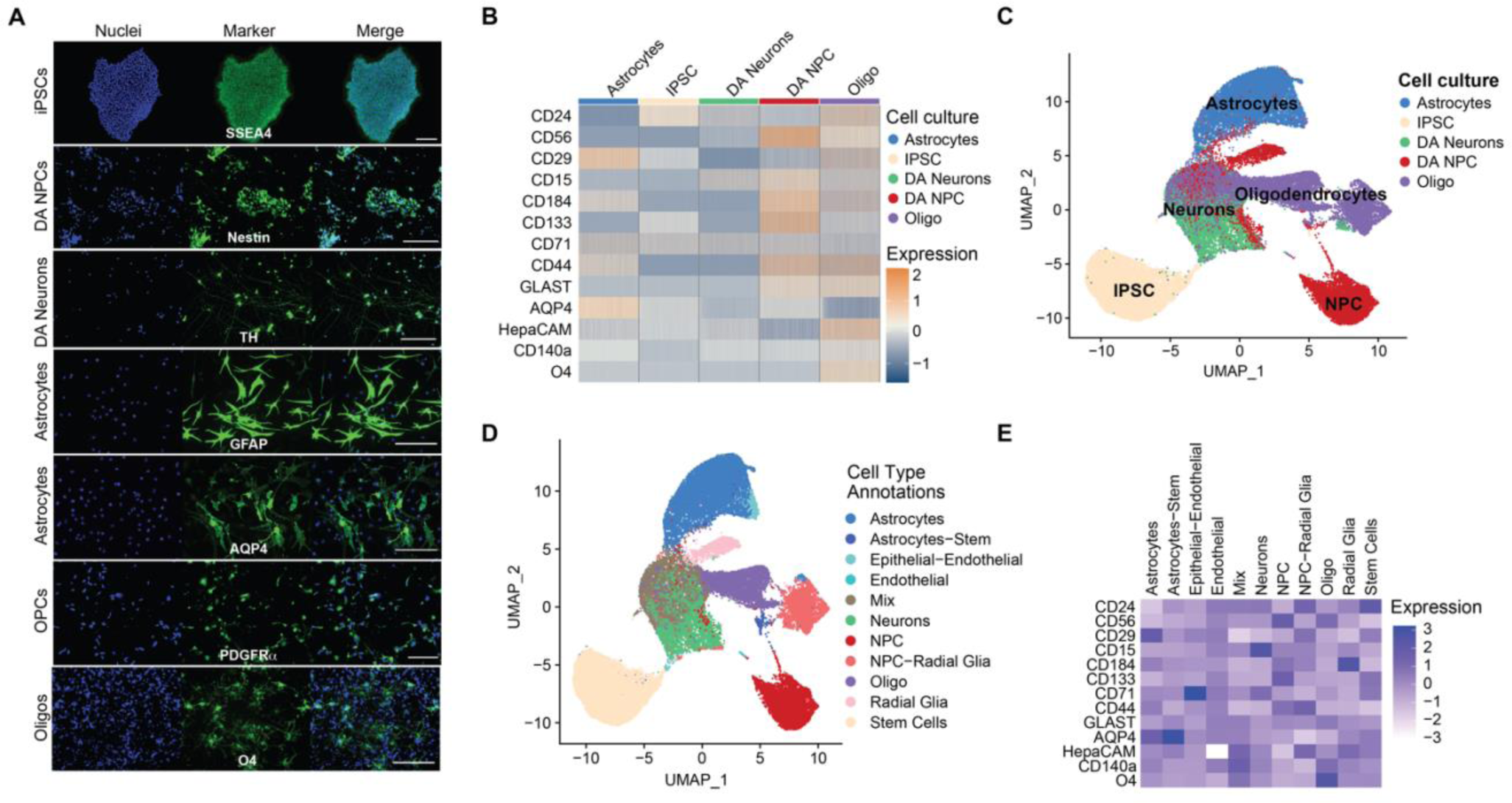
The antibody panel can be used to identify cell types expected to be present in hMOs. **A**) Example images of different brain cell types (indicated on the left) derived from the healthy control AIW002-02-02 human iPSC line and individually differentiated. Cell cultures were stained with a cell type specific marker (green) and Hoechst (blue) for nuclei. Scale bars 200mM. **B**) Heatmap of the normalized and z-scored protein levels measured by FC (area under the curve) for a subset of cells from each cell culture (indicated above). The marker proteins are indicated on the left. **C**) UMAP of merged cell cultures (indicated by colour) showing the separation and overlap of cell types **D**) The same UMAP with annotated clusters identified by Louvain network. Cell types are labelled by colour and in the legend. **E**) Heatmap of the mean expression of each protein within the cell subgroups identified by clustering. Values are z-scored.

### Identification of different brain cell types in human midbrain organoids

To identify cell types within hMOs using the antibody panel, we ran our R preprocessing pipeline to align and normalize the data. To compare samples from different iPSC lines, different batches of hMOs, and measurements run on different experiment days, we developed methods to combine and harmonize samples. This is the first step in the computational pipeline. We combined nine hMO samples and selected a subset of the total cells (9000 cells or the max number of cells available). The samples were first merged, then transformed and aligned to reduce batch effects, and finally retro-transformed for better cluster visualization (**Figure S5**). If removing batch effects is not desired (as in the separate cell cultures above), the preprocessing is stopped after merging. The hMOs contain a combination of neurons, neural precursor cells (NPCs), astrocytes, oligodendrocyte precursors (OPCs), oligodendrocytes, radial glia (RG), stem cells, pericytes, endothelial cells, and epithelial cells, all differentiated from the starting iPSCs.^4,23,37^ The standard method of manually defining cell groups using FlowJo or multiple scatter plots in R is time consuming and not reproducible across experiments. To overcome this barrier, we developed tools to identify cell types described below: A) A correlation cell type assignment model (CAM) using a custom reference matrix and B) clustering parameter exploration functions with tools to visualize and summarize of protein expression levels.

We created a reference matrix with the predicted relative expression of each cell surface marker in different cell types expected to be present in hMOs based on previous hMO data and human brain. Using scRNAseq data from human brain and organoids, total mRNA on brain cell types, and FC (see methods), we calculated the relative expression levels for each protein marker in our antibody panel (**Figure 3A**). The CAM function calculates the correlation of protein expression levels of the 13 markers in each hMO-derived cell to the expression levels of the same markers in the reference matrix we created, calculating the Pearson correlation coefficient (R). The R value is calculated for each cell type in the reference matrix. The cell type with the highest R value, above an adjustable threshold, out of the nine possible cell types is assigned for a given hMO derived cell (**Figure 3B**). Cells with R values below the selected cut-off are left unassigned. The FC panel contains 13 markers used as comparison points, thus an R value of 0.553 is required for a statistically significant correlation (p < 0.05). Applying this significance threshold, neurons are the most assigned cell type (**Figure 3C**). The number of assigned cells depends on the R threshold (**Figure S6**). Some hMO-derived cells correlated close to equally (within 0.05) with two cell types. When this was the case, these cells were assigned a merged cell type, and may represent an intermediated cell type, for example neurons and NPCs, which are the same cell type on a continuum of differentiation (**Figure S7 and S8**). The correlation assignment is a useful tool to provide biologists with a predicted cell type and guide annotation, however, it does not deliver the accuracy needed to quantify cell types across experiments. We therefore created tools to use CAM in combination with other methods. Clustering algorithms group together cells with similar expression profiles, thus cell that are not clearly identified as a given cell type in isolation can be identified based on their neighbours. We created functions to identify the topmost predicted cell types per cluster.

**Figure 3:**
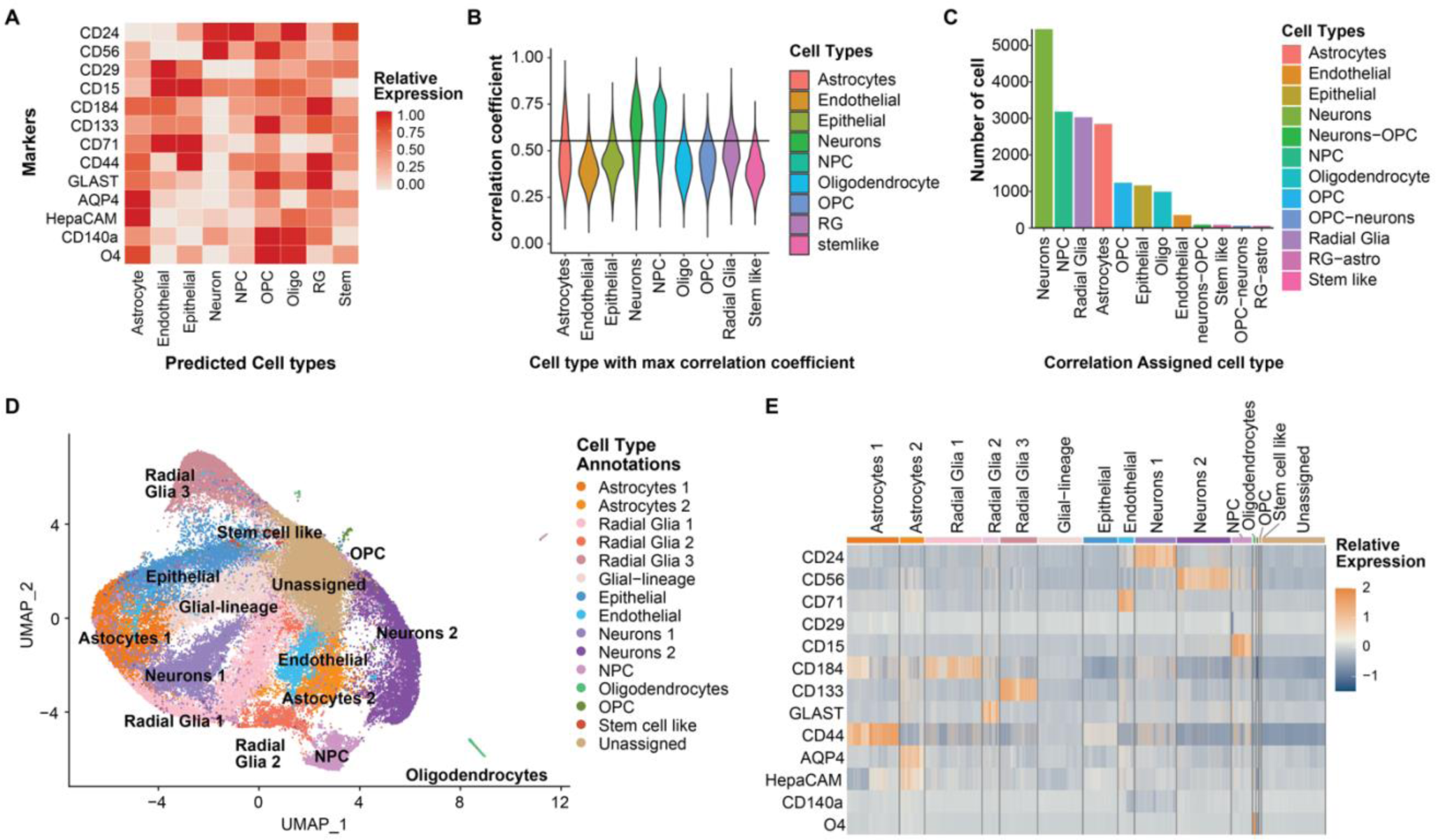
Identification of cell types in hMO using the flow cytometry antibody panel. **A)** Heatmap of predicted relative expression of each antibody in the FC panel for each potential cell type in hMOs. Values are calculated from 2D FC intensities, scRNAseq from hMOs and human brain, and RNAseq from human brain. **B)** Violin plot showing the distribution Pearson’s correlation coefficients (R) for hMO cells (y-axis) with the indicated potential brain cell type (x-axis). The R values are plotted for the cell type with the max R value. The black line indicates the threshold of R=0.45 which was set as the cut-off for assigning a cell type prediction. **C)** Bar chart showing the number of hMO cells categorized as each cell type by the max correlation. Each cell type is indicated on the x-axis. hMO cells were assigned as a double cell type if the first and second max R values were within 0.05. Only cell assignments with over 100 cells are included in the bar chart. **D)** UMAP showing unsupervised clustering by Louvain network detection. Cell types were annotated using a combination of correlation assignment and expert analysis of expression within clusters. **E)** Heatmap of relative expression of each antibody grouped by the cell types identified by unsupervised clustering of hMO cells. (n=73,578 cells from 9 hMO samples).

Using the functions in our CelltypeR library we performed unsupervised clustering using Louvain network detection and visualized the protein expression levels in each cluster (**Figure S9**). Clusters were annotated with cell types using a combination of marker expression by cluster and the output from the correlation predicted cell types (**Figure 3D**). We identified astrocytes, radial glia (RG), epithelial cells, endothelial cells, NPCs, neurons, a small proportion of oligodendrocytes and oligodendrocyte precursors (OPCs), and stem cell-like cells in the hMOs (**Figure 3E**). Clustering the hMO cells identified distinct subpopulations of RG, astrocytes, and neurons. All these cell types have a wide diversity in the brain and as well as in hMOs.^1,38^ For example, neurons 1 and 2 could represent different stages of maturation. We conclude that our workflow can be used to annotate cell types in hMOs and capture some diversity within cell types.

### Comparison of cell types between iPSC derived cell types and hMOs

After annotating a subset of 9000 cells from each of the nine hMO samples, we next analyzed the total available cells. We again followed the CelltypeR workflow and can now use the labelled subset of cells to annotate the full dataset. We first clustered the full dataset and then annotated the cells from the nine hMO samples (**Figure 4A**). Using CelltypeR functions, we trained a random forest classifier model (RFM) to predict cell types (**Figure S10**). In addition to analyzing protein expression profiles by cluster, we used three prediction methods (CAM, RFM, Seurat label transfer) to annotate cell types (**Figure S11 and S12**). We observe the same cell types in the full dataset as in the subset of data; however, we now identify a cell group predicted to be OPCs and Radial Glia 1, which we termed OPC-like (**Figure 4A**). We examined the protein expression levels within our cell type annotations and distinctive expression profiles (**Figure 4B**).

**Figure 4:**
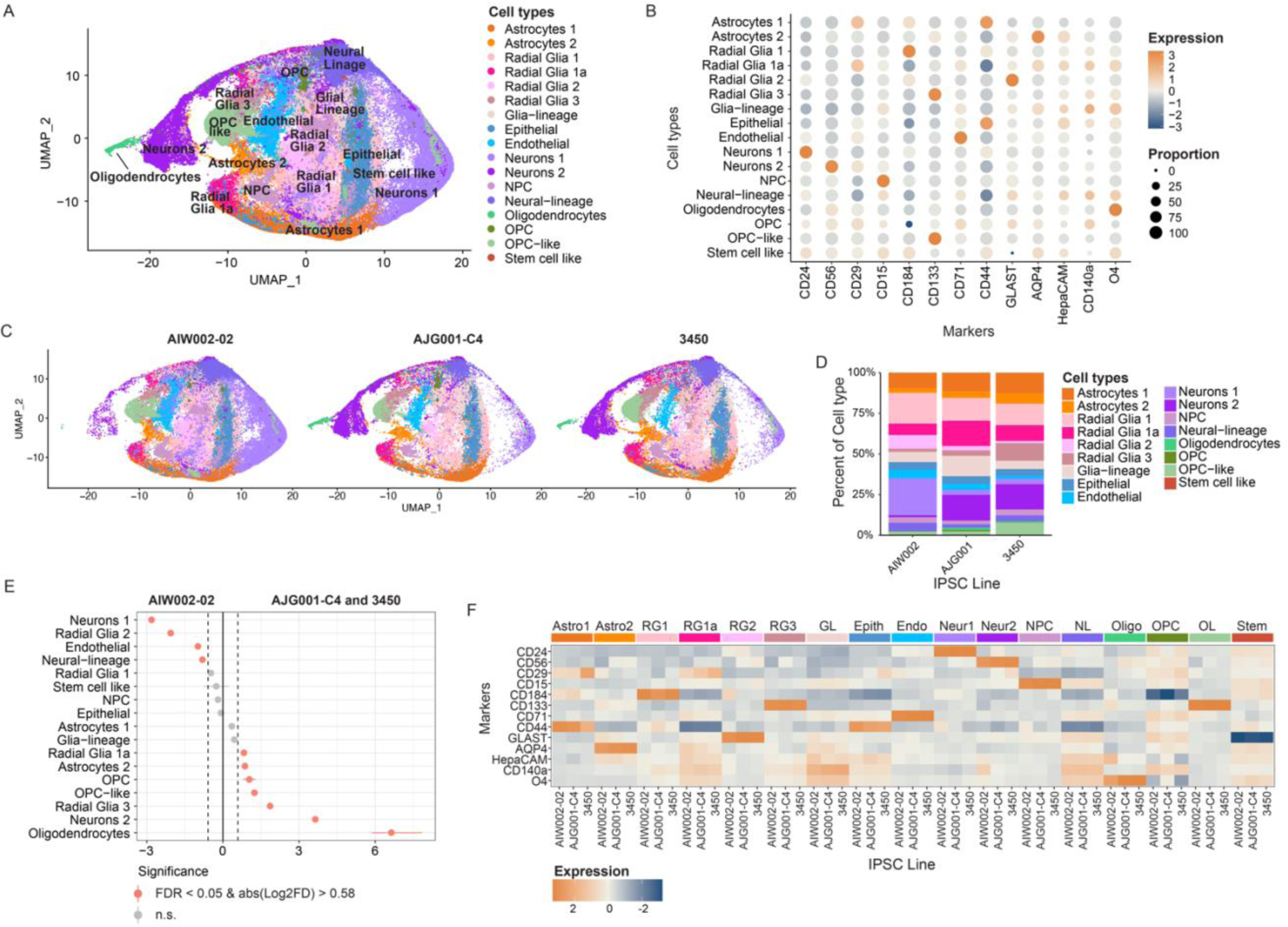
Differences in cell types and protein expression between three healthy control donor iPSCs. **A)** UMAP of the full dataset from nine hMO samples (n > 197,000 cells) annotated using CelltypeR. **B)** Dot plot of the expression level (colour intensity) and the proportion of cells (dot size) for each protein marker detected with the panel in each cell type group. Scaled z-score values are shown. **C)** UMAP split by iPSC line (3 samples pooled per iPSC line) showing the proportion of cells in each iPSC line. Cell annotations and colours are the same as the UMAP in A. **D)** Bar chart of the proportion of hMO cells in each cell type (indicated by colour) for each iPSC line (x axis). Colours corresponding to cell types are shown in the legend on the right. **E)** Dot plot with confidence interval for the proportionality test comparing the AIW002-02 iPSC line to the AJG001-C4 and 3450 iPSC lines, for each cell type (y-axis). Pink dots indicate a significant difference in cell type proportion (FDR < 0.05 and absolute value of Log2FD > 0.58). Negative log2FD values indicate cell proportions increased in AIW002-02 and positive values indicate cell proportions decreased in AIW002-02 compared to the other two iPSC lines. **F)** Heatmap of mean protein expression values grouped by cell type and split into the three iPSC lines. Line names are indicated on the bottom x-axis and cell types are indicated on the top x-axis. Scaled z- score values are shown.

Visualizing the distribution of cell types in hMOs derived from each cell line, we can see there are some differences in the proportion of cell types (**Figure 4C,D**). We observe more Neurons 1 and fewer Neurons2 in the AIW002 hMOs compared to the other two lines. The AIW002 hMOs also have less oligodendrocytes than AJG001-C4 and 3450 hMOs. We next preformed permutation tests to determine if the differences in proportions of cell types between the cell lines are significant. We find in addition to changes in neurons and oligodendrocytes cell proportions, subtypes of glia cells are also significantly different (**Figure 4E**). Comparing pairwise combinations between the three iPSC lines we also see differences across all lines. Notably AJG001-C4 hMOs have the most oligodendrocytes and OPCs, and fewer NPCs than the other two lines (**Figure S13**). To visualize if expression patterns differ within one cell type between iPSC lines, we plotted a heatmap of mean protein expression and observe most proteins have consistent expression across iPSC lines in most cell types (**Figure 4F**). To further explore expression differences between groups, we created functions in our R package to run ANOVAs, post-hoc tests, and identify significant differences. We performed two-way ANOVAs followed by Tukey’s post-hoc tests to compare expression levels for each marker protein across iPSC lines in each cell type. There are significant differences in overall marker expression levels between the lines in neurons, NPCs, oligodendrocytes, and OPC-like cells (**Table S1**). A few significant differences in expression of one protein are seen between pairs of iPSC lines (**Table S2**). Using our framework, we can reliably quantify cell types and compare proportions of cells and levels of antibody expression across different conditions. We find significant differences in the proportion of cell types and in marker expression levels within cell types between different healthy control iPSC lines.

### Application of the CelltypeR workflow to new datasets

We next generated new batches of hMOs using the control cell line AIW002-02 to validate the CelltypeR workflow on a new dataset. The antibody panel was applied, and intensity levels were measured by FC from two different batches on one experiment day, and on one of the batches on a second experiment day. The cell types in these new batches were processed as previously described above and in the methods. The cell types were annotated using CelltypeR functions. For RFM and Seurat label transfers, the cell type labels from Figure 3D were used as the reference data. The cell types in the new hMO samples were found to be consistent with the original AIW002-02 samples (**Figure 5A**). To determine if the proportion of cells was similar across the two new and two original AIW002-02 batches, we plotted the percentage of each cell type grouped by hMO batch and observe similar but varying proportions of cell types across batches (**Figure 5B**). Batches A and B are from the original data set and were grown with a different protocol than the two new batches C and D (see methods). We next performed permutation tests and find more differences in cell type proportions between batches A,B and C,D than between the batches grown with the same protocol (**Figure S14**). The relative proportion of several radial glia populations are increased in batches A and B compared to batches C and D. Whereas *Neurons 2*, *OPCs*, *oligodendrocytes* and *stem cell like* populations are all relatively decreased in batches A and B compared to batches C and D (**Figure 5B and S14**). We concluded that CelltypeR can identify cell types across data sets separately processed and compare between batches.

**Figure 5:**
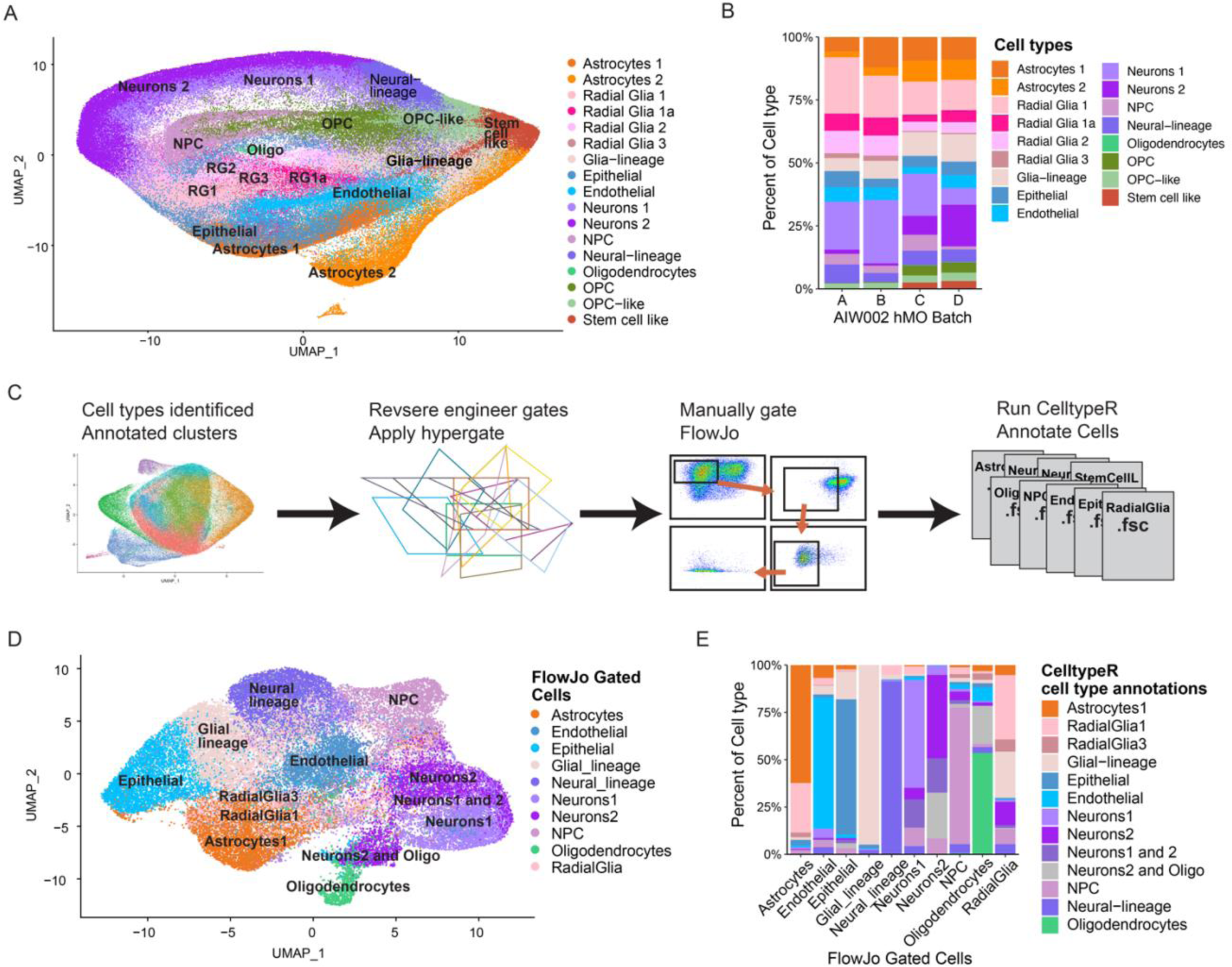
CelltypeR can be used to identify cell types in a new population, gate populations of interest and annotate the gated cells. **A)** Two new batches of AIW002-02 hMOs were processed with the CelltypeR workflow and cell types were annotated. UMAP shows cells from seven samples dissociated and acquired on two different days. **B)** Bar chart showing the proportions of cell types across four different batches of AIW002-02 hMOs. Batches A and B are the samples from the 9 hMO comparisons, batches C and D are the new samples shown in panel A. **C)** Schematic showing the method used to gate cell type populations defined with CelltypeR. Cell types were annotated and selected in the full 9 hMO dataset. Then the package *hypergate* was applied to reverse engineer the threshold expression levels to define each cell population. **D)** UMAP coloured by the FlowJo gated populations (legend), gated using the thresholds and markers selected by *hypergate.* Astrocytes, radial glia, oligodendrocytes, epithelia cells, endothelial cells, NPCs, neurons 1, and neurons 2 cell populations were exported as fsc files and input into the CelltypeR workflow. Gated cells were down sampled to 5000 cells, except for oligodendrocytes where all were included. The labels on the UMAP are the cell types annotated using the CelltypeR workflow. **E)** Bar chart with the proportion of cell types identified with CelltypeR (indicated by colour in the legend) within each FlowJo gated population (x-axis).

### Enriching and purifying populations of interest identified by CelltypeR clustering analysis

After annotating a dataset, we could plot the proportion and mean expression of each antibody marker in each group to try and define the relative marker expression of a given cell group and then isolate that population by FACS. However, manually reverse engineering a gating strategy is difficult with more than a few cell type markers. Thus, we defined cell types using CelltypeR, applied the package *hypergate*^39^ to identify which combinations of antibody markers clearly define a given cell population, and then manually gated these cells in FlowJo (**Figure 5C**). To reduce the number of potential populations to gate, subgroups of the radial glia cells and astrocytes were merged. *Hypergate* was applied to define the threshold for each antibody relevant for gating. The resulting gating accuracy for all cell types is above 95% **(Table S3)**. We next used the defined gates in FlowJo to gate the cell type populations (**Figure 5D**). After gating in FlowJo we found that OPC, OPC-like and stem cell like cell populations were much larger than expected and we didn’t include these populations in the next steps. The remaining gated populations were entered into the CelltypeR workflow and clusters of cells were annotated (**Figure 5D**). Some clusters clearly contained two cell types, and these were labelled as such to reflect the mixtures. The proportion of CelltypeR cell types within each FlowJo gated population was calculated (**Figure 5E**). The most frequent annotated cell type within each gated population is the intended cell type. Merging the populations, astrocytes resulted in gating the *astrocytes 1* population and not enough of astrocytes 2 to be detected. The merged radial glial populations resulted in a less effective gate that only selected *radial glia 1* and *radial glia 3* populations. We concluded that our workflow can be used to isolate selected populations, but these populations must be carefully selected and tested in FlowJo.

### Analysis of purified neuronal and glia populations followed by single cell sequencing analysis

Our workflow can be used to enrich populations of interest by FACS, separating selected populations for further analysis. We selected four cell types that were gated well in FlowJo and were of interest for further study: neurons 1, neurons 2, astrocytes, and radial glia. We next designed a gating strategy to simultaneously sort the four populations (**Figure 6A**). We sorted the hMO-derived cells using the defined gates, split the samples, and then acquired FC measurements and scRNAseq on the sorted populations. The protein expression levels in the sorted populations match the expected levels from the gates (**Figures 6B**). We also obtained a single cell transcriptomic library for each of the FACS populations (see methods). We first compared the RNA expression levels of the genes corresponding with the protein expression levels measured by FC and found they highly correlate (**Figure 6C and Table S4**). The scRNAseq libraries from the four populations were merged, clustered, and plotted on a UMAP to visualize the overlap between the different sorted cell types (**Figure 6D**). The *Neurons1* population is mostly separate from the other populations with some overlap with *Neurons2*. Clusters were first annotated for main groups of cell types: DA neurons, neurons, NPCs, radial glia, and astrocytes and the proportion of these cell types in each sorted population was calculated (**Figure 6E**). These main cell types were isolated and annotated for subtypes of cells using differential gene expression between clusters (**Figure 6F, S15-18 and Table S5)**. To annotate the three subgroups of DA neurons we used cluster markers and compared expression with published markers (**Figure S16 and Tables S6 and S7)**. Next, we examined the proportion of cellular subtypes in the sorted populations (**Figure S19 and Table S8**). We found that non-DA neurons in *Neurons1* are excitatory neurons, neurons with potential to be reactivated as neural stem cells, NPCs, and ventral zone (VZ) radial glia undergoing neurogenesis (**Figure S15**). The non-DA neurons in the *Neurons2* population are GABAergic, serotonergic (5HT), and endocrine neurons (**Figure S15)**. As quantification of DA neurons is of particular interest in hMOs for PD, we calculated the proportion of all the DA neurons and the three subtypes of DA neurons in the sorted populations. We find that Neurons1, Neurons2, and Radial Glia all contain DA neurons (**Figure S20 and Table S8**). The *Neurons1* FACS population has slightly more DA neurons overall, and specifically more of the substantia nigra (SN) subtype, whereas the Neurons2 FACS population has more cells from the cluster identified as general ventral midbrain (VM) DA neurons. The subgroup containing ventral tegmental area (VTA) DA neurons didn’t show a significant differential distribution between the *Neurons1* and *Neurons2* FACS groups (**Figure 6G and S20)**. Thus, the two sorted neuron populations contain distinctive subtypes of DA and non-DA neurons. Moreover, the astrocyte population also segregated into three subgroups, immature, resting, and reactive (**Figure S17 and S19)**. The radial glia population contains five different subtypes (**Figure S18 and S19**). Taken together, our findings indicate that each FACS population is enriched in the expected cell type and there are identifiable subtypes within these groups, confirming the effectiveness of the CelltypeR framework.

**Figure 6:**
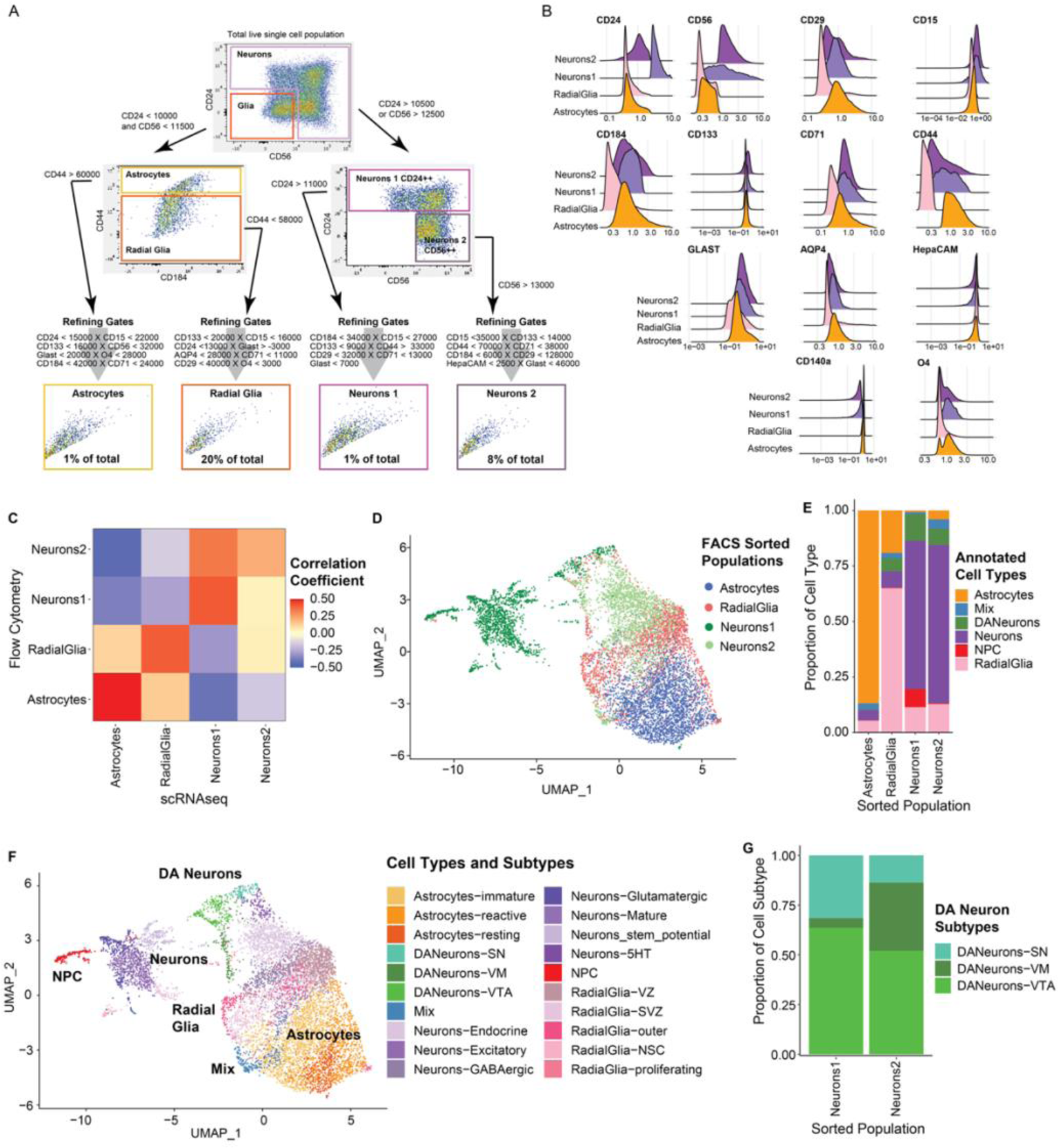
scRNAseq analysis of four FC sorted populations defined using CelltypeR confirms cell types and provides transcriptional profiles for these cell populations. **A)** FlowJo gating strategy applied to new hMO derived cells to isolate four cell populations by FACS: neurons1, neurons2, astrocytes, and radial glia. The approximate proportion of cells gated in each final sorted population is indicated in the gating box. **B)** Ridge plot of protein expression levels measured by FC antibody intensity for each FACS gated cell population. **C)** Correlation of RNA transcript expression of genes corresponding to the 13 protein markers used for FACS. Note there is a high correlation between RNA expression and protein expression for radial glia, neurons1, and astrocytes. Only the astrocytes RNA to protein correlation was statistically significant. The neurons2 protein expression correlates more strongly with the neurons1 RNA expression. **D)** UMAP of the four sorted populations merged and clustered with Louvain network detection. Neurons1 has only 1723 cells, neurons2 was down sampled to 2000, astrocytes were down sampled to 3000, and radial glia were down sampled to 2000 to improve visualization. The original FACS sorted population is indicated by colour and in the legend. **E)** Stacked bar chart of the proportion of each main cell type identified by the cluster transcriptomes in each FACS sorted population. **F)** UMAP of the four merged populations with cell types and cell subtypes annotated from the scRNAseq data. The UMAP is coloured by cell subtypes and the main cell types are labelled on top of the UMAP. Subtypes were identified from differential RNA between clusters to identify markers and comparison with reference datasets. **G)** Stacked bar chart showing the proportion of each DA neuron subtype within each sorted neuron population.

### CelltypeR workflow on a time course of organoids to identify changes in cell type proportion over time

To display the ability to adjust and refine the antibody panel we decided to change our antibody panel and apply the pipeline to a time course of hMOs in one genetic background. To directly track DA neurons we added TH, the only protein with a cytoplasmic epitope to the panel. Adding a cytoplasmic marker does slightly increase the experiment time, but it is still feasible for multiple samples (see methods). We made other alterations to the panel to display the flexibility of our CelltypeR pipeline. We added “SSEA4”, a pluripotency marker^40^ and “CD49f”^41^, reported to be a marker of activated astrocytes(ref) and removed CD71, AQP4 and GLAST, HepaCam and O4 (**Table S9**). A new reference matrix was created for the adapted marker panel, now including DA neurons and DA NPCs (**Figure 7A**). We acquired FC measure of protein expression across four time points in one batch of AIW002-02 hMOs and followed the CelltypeR workflow. Using CAM prediction with the new reference matrix, RFM and Seurat label transfer predictions from the overlapping antibodies and expression analysis to annotate clusters (**Figure 7B**). To determine if the proportion of cell types are changing over time in these hMOs, we plotted the proportion of cell types at each time point and found that cell types are changing over time and show less variation between replicates (**Figure 7C and S21**). We observe that the proportions of both DA neurons, neurons and DA-NPCs increase from 38 to 63 days, then decrease from 98 and 155 days. Stem cell like and glial populations show an increase over time. To test if the changes in the proportions of cell types are significant, permutation test were performed. There are significantly more DA-neurons at 63 than 38, but not between 63 and 98 days, then significantly fewer DA neurons at 155 days (**Figure 7D**). Using the proportion of each cell type we performed a two-way ANOVA with Tukey’s HSD post-hoc tests to determine which cell types were changing over time. DA-NPCs are significantly different between all-time point comparisons, however DA neurons are only different between 63 and 155 days (**Table S10**). The changes in cell type proportions match with the expected time course of differentiation, as stem cell and early glial cells will continue to divide where-as neurons and mature astrocytes do not divide. We conclude that the CelltypeR workflow can be applied with an altered antibody panel and is useful for detecting differences in cell type populations over time.

**Figure 7:**
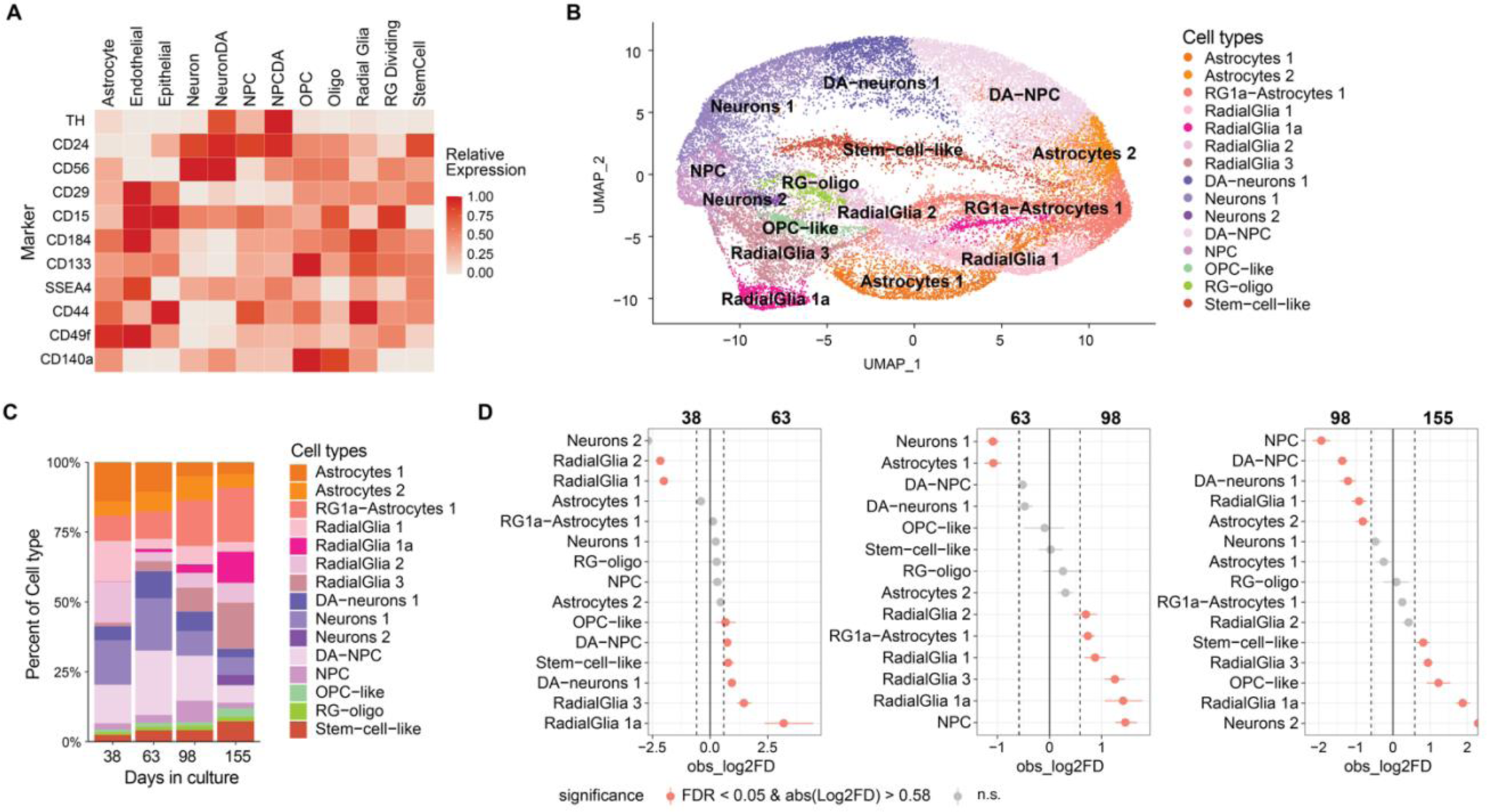
Cell type proportions in hMOs change over time in culture. **A)** Heatmap of predicted relative expression of each protein targeted in the new FC panel for each potential cell type in hMOs. Values are calculated from 2D FC intensities, scRNAseq from hMOs and human brain, and RNAseq from human brain. **B)** UMAP of cells from all four time points, annotated using the CelltypeR workflow. **C)** Barchart of the proportions of cell types for each time point. Each time point has 4 technical replicates and combined values are used. **D)** Proportionality tests comparing time points in pairs, from left to right: 30 days vs 60 days, 60 days vs 100 days and 100 days vs 150 days. Differences that have a change in proportion > 0.58 logFold change and an adjusted p value < 0.05 are shown in pink.

## Discussion

Taken together, we present the first comprehensive antibody-based workflow to identify, quantify, compare, and sort cell types in complex 3D tissue, specifically hMOs. We define a 13-antibody panel that can be used to distinguish between eight different brain cell types and identify subtypes of astrocytes, radial glia, and neurons. The antibody panel is modular, such that it can be altered or expanded for any organoid or tissue type and will function with the computational workflow. We show an example of this by generating an altered antibody panel using the cytosolic TH antibody to detect DA NPCs and neurons. In our CelltypeR library, we provide a method to preprocess and merge FC samples, acquired from multiple samples at different dates. We created tools to optimize and visualize clustering and to assist in consistent cell type annotation. We also created functions to quantify cell types and compare different conditions.

The same workflow with sorting can be used to isolate a more homogenous subpopulation of a given cell type for use in other assays that includes but is not limited to proteomics, lipidomics, testing small molecules, or even replating of cells in culture to grow as a purified population. Here we selected four populations, FACS sorted the cells, and then performed scRNAseq analysis. We confirmed that each of the populations, *Neurons1*, *Neurons2*, *Radial Glia* and *Astrocytes*, are all highly enriched in the expected cell types. Further analysis of the scRNAseq data identified subtypes within each cell type group. We identified DA neurons within both neuronal populations but find different DA neuron subtypes enriched in each of the two FACS sorted neuron populations. As an example, we identified TPGB as a DA subtype marker (ventral), in agreement with a recent publication proposing TPGB as a marker of ventral DA neurons in mice.^42^. The DA neurons of the SN are selectively lost in PD, whereas the VTA neurons are important for studying addiction and the reward system. We see an enrichment of SN-like neurons in one of our sorted neuron populations. Isolating specific subtypes of DA neurons is fundamental for research targeting specific disease pathways, like those leading to SN DA neuron cell death in PD. We also identified VTA DA neurons which were not specifically enriched in either sorted neuron population. The third group of DA neurons were identified as ventral midbrain DA neurons but were not specific for SN or VTA.

Using an adjusted antibody panel, we applied the CelltypeR workflow to track cell type proportions over time in AIW002-02 hMOs. We found that the proportion of DA-NPCs, neurons 1, and DA neurons all follow a similar pattern, increasing from 38 days to 63 days and then decrease at later time points. Neurons 2 only appear at the 155-day time point, indicating this group could be more mature. The neural populations within the organoids exhibited intriguing temporal dynamics. It’s important to note that this decrease in relative neuronal proportion does not necessarily signify a reduction in the total number of neurons but rather suggests changes in their proportions within the overall cell population. Additionally, we observed relative increases in radial glia 1a, radial glia 3, stem cell like cells, and OPC-like cell populations. These cells populations are likely proliferating and represent partially differentiated cells, which possess the potential to differentiate into astrocytes, neurons, or other brain cells over time.

In our analysis of the differences between three healthy control iPSC lines, we find a clear difference in the proportion of cells for the two subtypes of neurons between AIW002-02 compared to the other two lines, AJG001 and 3450. AIW002-02 has more *Neurons1* with higher CD24 expression and fewer of the *Neurons2* population, with lower CD24 expression than AJG001 and 3450. scRNAseq reveals the *Neurons1* population has more NPCs and DA neurons. We also find that AIW002-02 has more radial glia, fewer astrocytes, and fewer oligodendrocytes than the other two lines, indicating that this cell line may be less mature. AIW002-02 might mature at a slower rate or given the late age of the organoids, maintain cell populations with potential to become precursors perpetually. These findings also indicate that to study the role of myelination, the AJG001 or 3450 lines could be a better choice than AIW002-02.

In conclusion, we established an adaptable method for reproducibly identifying and quantifying cell types in complex 3D tissues, such as hMOs, using a flow cytometry panel. Our scalable single-cell biology workflow enables rapid and efficient cell type quantification across multiple replicates and experimental conditions. Additionally, we have highlighted the potential of our workflow in the context of tracking cell type dynamics over time, comparative analysis of iPSC lines, and its adaptability for diverse tissue types. Furthermore, we emphasize the importance of creating reference matrices for changes in the antibody panel, thereby enhancing the utility and applicability of the CelltypeR workflow in various research domains.

## Methods

### 1. Human midbrain organoids (hMOs)

#### 1.1 iPSC lines

Three iPSC cell lines were used: AJG001-C4-C4, AIW002-02-02, and 3450. All were previously reprogrammed from peripheral blood mononuclear cells as previously described.^22^ All work with human iPSCs was approved by McGill University Faculty of Medicine and Health Sciences Institutional Review Board (IRB Internal Study Number: A03-M19-22A).^22^

#### 1.2 hMO cultures

hMOs were derived from iPSC lines using two different protocols.^4,43^ For batches A and B (**Table M1**) iPSCs were seeded in separate ultra-low attachment plates in neural induction medium for embryonic bodies (EBs) to form. On day four, medium was changed to midbrain pattering medium to promote a dopaminergic neural cell fate. On day seven, hMOs were embed in Matrigel. On day eight, hMOs were transferred to 6-well plates with 4-6 hMOs per cell line in organoid growth media and placed in shaking cultures.^4^ hMOs were maintained in shaking cultures with media change every 2-3 days. For batches C, D, and E, iPSCs were seeded in eNuvio disks for EB formation and Matrigel embedding, then transferred to bioreactors for culture maintenance.^43^ Media changes were performed weekly, and all the same growth mediums were used in both protocols.

#### 1.3 hMO FC samples

The iPSC lines and time points for FC acquisition are shown in **Methods Table 1**. The date of seeding iPSCs, 8 days before final differentiation is indicated. Days in culture is the time between transfer to final differentiation and FC. The date of dissociation, labelling and acquisition (FC acquisition date) is indicated. The line AIW002-02 was used to generate hMOs for follow up experiments with FACS gates defined in the CelltypeR workflow. In sorting experiments FC measurements were taken before sorting from a sample of the total proportion of cells and post sorting on a small portion of the resulting sorted sample. The cell counts refer to the data acquired pre-sorting not the post sorted populations. The total values from all replicates are shown in #live cells. The number of days in culture is calculated from the first day of final differentiation media to the experiment date (FC acquisition date).

**Methods Table 1:**
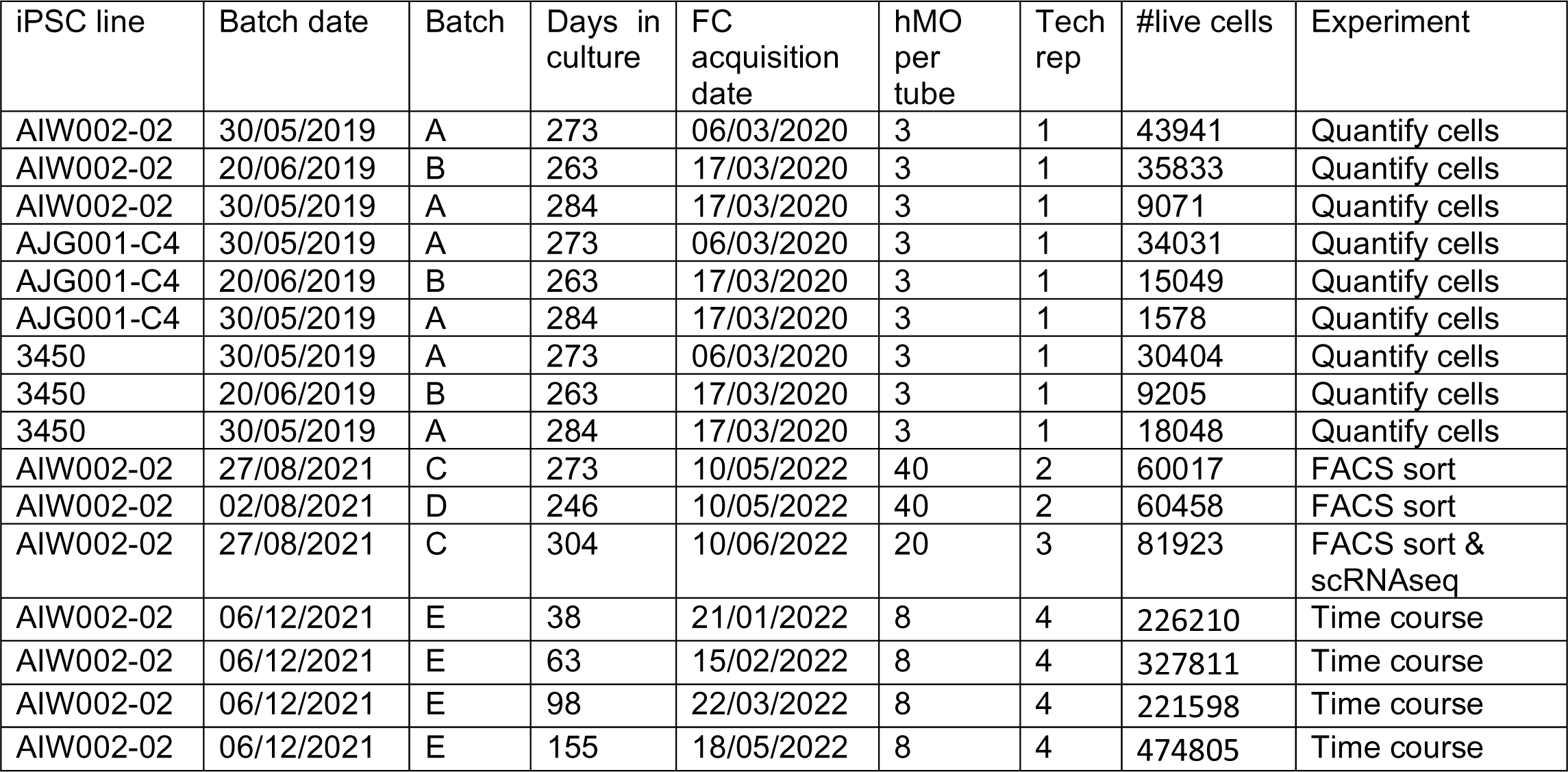
Description of hMO data sets.

### 2. 2D Cell culturing conditions

#### 2.1 iPSC

The control cell line AIW002-02 was used for all 2D cell cultures. Prior to differentiation, the iPSC cultures were maintained and expanded on Matrigel coated plates and grown in either mTeSR1 or E8 media as previously described.^22^

#### 2.2 Dopaminergic neural precursor cells (DA-NPC) and neurons (DA neurons)

DA-NPC cultures were generated by dissociating iPSCs into single cell suspensions and then culturing these cells in low attachment plates to generate EBs. EBs were re-plated onto polyornithine and laminin-coated plates and differentiated into neural rosettes, which were then differentiated into DA-NPCs. DA neurons were differentiated from DA-NPC cultures on laminin coated culture flasks in neural basal media with supplements and inhibitors as described.^35^

#### 2.3 Oligodendrocyte precursor cells (OPCs) and oligodendrocytes

To derive OPCs and oligodendrocytes we used a three-phase protocol as previously described.^34^ In phase one, iPSCs were induced towards neural progenitors while being patterned with Retinoic Acid in order to resemble spinal cord progenitors. The Sonic Hedgehog pathway was activated for ventral patterning to recapitulate the conditions of the oligodendrocyte fate. The progenitors were subsequently expanded as EBs with the addition of the bFGF. In phase two, OPCs were expanded in suspension and subsequently plated onto polyornithine/laminin-coated vessels for adhesion. Growth factors and mitogens were added in the medium for differentiation and maintenance of the OPCs, respectively. Images of PDGFRα positive cells were acquired at this phase. In phase three, mitogens are withdrawn to allow the progenitors to exit the cell cycle and to complete differentiation into oligodendrocytes. Imaging and FC were performed in this phase when oligodendrocytes would generate O4 positive cells.

#### 2.4 Astrocytes

Astrocytes were derived from NPCs cultures as previously described.^36^ NPCs were seeded at low cell density and grown in NPC expansion medium. The next day, medium was replaced with ‘Astrocyte Differentiation Medium 1’. Cells were split 1:4 every week and were maintained under these culture conditions for 30 days. At DIV50, cultures were switched to ‘Astrocyte Differentiation Medium 2’ and maintained with half medium changes every 3-4 days.

#### 2.5 2D cultures used in FC acquisition

**Methods Table 2:**
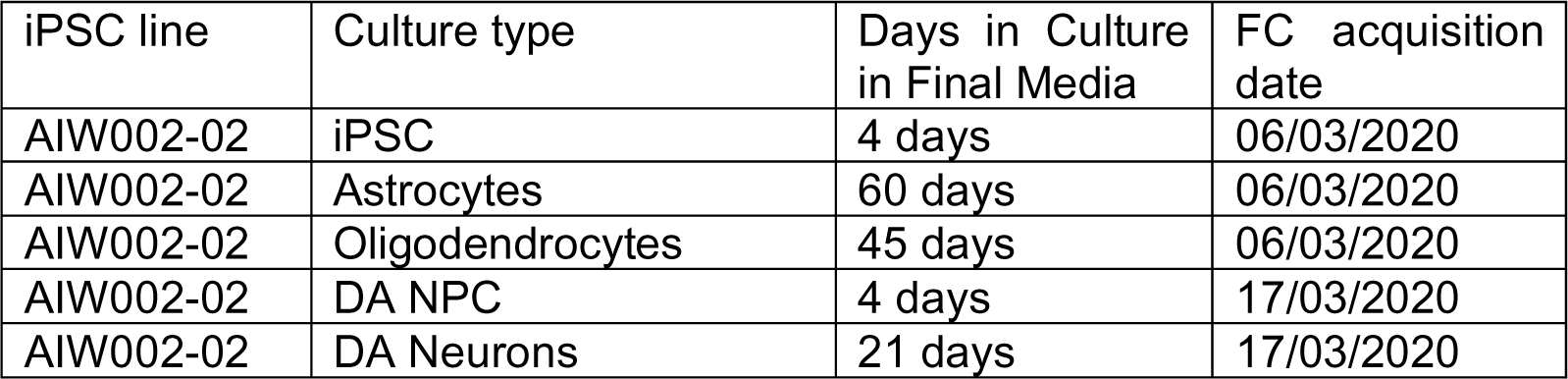
Description of 2D cultures.

### 3. Immunofluorescence

#### 3.1 iPSC, NPCs and Dopaminergic Neurons

Cells were fixed in 4% PFA/PBS at RT for 20 minutes, permeabilized with 0.2% Triton X-100/PBS for 10 min at room temperature (RT), and then blocked in 5% donkey serum, 1% BSA and 0.05% Triton X-100/ PBS for 2h. Cells were incubated with primary antibodies: MAP2 (1:1000, EnCor Biotech CPCA-MAP2); Nestin (1:500, Abcam ab92391); SSEA-4 (1:200, Santa Cruz Biotechnology sc-21704); in blocking buffer overnight at 4 °C. Secondary antibodies were applied for 2h at RT, followed by Hoechst 33342 (1/5,000, Sigma) nucleic acid counterstain for 5 minutes. Immunocytochemistry images were acquired using Evos FL-Auto2 imaging system (ThermoFisher Scientific).

#### 3.2 Astrocytes

Cells were fixed for 15 minutes at room temperature with 4% formaldehyde in PBS, followed by 3 washes of 5 minutes in PBS. Cells were permeabilized for 10 min at RT in blocking solution: 5% normal donkey serum (JacksonImmunoResearch Laboratories, West Grove, PA), 0.1% Triton-X-100, and 0.5 mg/ml bovine serum albumin (Sigma-Aldrich) in PBS. Cells were incubated for 1h at RT before overnight incubation at 4°C with primary antibodies: Glial Fibrillary Acidic Protein (GFAP) (1/500 Dako Cat. Number Z0334); AQP4 (1/500, SIGMA, cat# HPA014784). Secondary antibodies were incubated 2h at, followed by Hoechst 33258 (1/5,000, Sigma) for 5 min, mounted with Fluoromount-G, and examined by fluorescence microscopy.

#### 3.3 Oligodendrocytes and OPCs

Cells were fixed in 2% PFA for 10 min and blocked in 5% BSA, 0.05% Triton for 1h. Mouse anti-O4 (R&D, MAB1326) was added in live cells before fixation for 1h at a final concentration of 1μg/mL. Rabbit anti-PDGFRa (Cell Signaling, 3174) was added post-fixation at a dilution of 1:200 and incubated overnight at 4°C. Secondary antibodies were added at a dilution of 1:500 and incubated for 2h at RT. Nuclei were identified with incubation with Hoechst 33342 (1/5,000, Sigma) for 5 min.

#### 3.4 Midbrain organoids

hMOs were washed in PBS, and then fixed for 2h in 4% PFA diluted in PBS at RT, then placed in a sucrose gradient overnight at 4°C. hMOs were then embedded in Optimal Cutting Temperature Compound (OTC) (Fisher Healthcare 23-730-571) and frozen. Cryosections of 20μM were cut using Cryostat Cryostar NX70 (Thermo Scientific). The slides with the sections were rehydrated in PBS for 15 minutes and surrounded by a hydrophobic barrier using a hydrophobic pap pen. Sections were then blocked for 1 hour in blocking solution (5% Normal Goat Serum (Jackson Immuno Research Laboratories, West Grove, PA), 0.05% BSA (Sigma-Aldrich), 0.2% Triton X-100 in PBS), and incubated with primary antibodies diluted in blocking buffer: anti-O4 (1:200, R&D, MAB1326); Glial Fibrillary Acidic Protein (GFAP) (1/500 Dako Cat. Number Z0334); MAP2 (1:1000, EnCor Biotech CPCA-MAP2), Tyrosine-Hydroxylase (TH) (1/500 Pel-Freez P40101), Neurofilament (NF) (1/500 Sigma N5264), GIRK2 (1/250 NovusBio NB100-74575), FOXA2 (1/500 abcam ab117542), AQP4 (1/500 Sigma HPA014784), S100b (1/500 Sigma S2532) at RT for 1h. Fluorescent-labeled secondary antibodies (Invitrogen) were added at a dilution of 1:500 and incubated for 1 hour. Nuclei were identified with Hoechst 33258 (1:5000, Sigma) diluted in PBS and incubated with the cryosections for 10 minutes at RT. Cover slides were mounted using Aqua-Poly/Mount mounting medium (Polysciences) and imaged using confocal microscopy (Leica TCS SP8 confocal).

### 4. Sample preparation for flow cytometry

#### 4.1 Tissue dissociation and processing – Main data set hMOs

hMOs were dissociated with a combination of enzymatic digestion and mechanical dissociation. First, three individual hMOs from each of the data set of nine samples were removed from shaking cultures and combined into one 15mL tube. Pooled hMOs were washed three times with Dulbecco’s PBS (D- PBS) (Wisent) to completely remove remaining culture media. Then, after completely removing D- PBS, 2mL of TrypLE express (without phenol red) (ThermoFisher) was added to each sample. The hMOs were incubated at 37°C for ten minutes then removed to be subjected to mechanical dissociation by pipette trituration (slowly pipetting up and down ten times). The incubation and the pipette trituration are repeated twice more. Afterwards, 8mL of D-PBS was added to the samples to stop the enzymatic reaction. The samples were filtered through a 30µm filter (Miltenyi Biotec) to remove any clumps remaining after digestion and dissociation. Samples were washed twice more with D-PBS.

#### 4.2 Tissue dissociation and processing – Sorting data set hMOs

hMOs were dissociated with a combination of enzymatic digestion and mechanical dissociation. First, twenty individual hMOs were removed from a bioreactor and combined into one 50mL tube. Pooled hMOs were washed three times with Dulbecco’s PBS (D-PBS) (Wisent) to completely remove remaining culture media. Pooled hMOs were transferred to a gentleMACS M-Tube (Miltenyi Biotec). Then, after completely removing D-PBS, 2mL of TrypLE express (without phenol red) (ThermoFisher) was added to each sample. The hMOs inside the M-Tube are then next placed on an automated GentleMACS Octo Heated dissociator. The settings for the dissociation were as follows: 37°C is ON. Spin -20rpm for 24 minutes. Spin 197rpm for 1 minute. After incubation, 8mL D-PBS was added to the samples to stop the enzymatic reaction. The samples were filtered through a 30µm filter (Miltenyi Biotec) to remove any clumps remaining after digestion and dissociation. The samples were then washed twice more with D-PBS.

#### 4.3 Tissue dissociation and processing – 2D cell cultures

T-flasks containing cells were washed in PBS then incubated at 37°C in 2mL of TrypLE express (without phenol red) (ThermoFisher) for 5-20 minutes depending on cell type. Cells were washed off the growth surface with a pipette, then manual dissociated by trituration until no clumps were seen and transferred to a 15ml tube. Cells were washed twice in D-PBS.

#### 4.4 Antibody staining – All samples

After counting and isolating one million cells, single cell suspensions were incubated for 30 minutes at room temperature in the dark with Live/Dead Fixable dye to assess viability. Single cell suspensions were washed twice with D-PBS to remove any excess dye. After, single cell suspensions were incubated for 15 minutes at room temperature in the dark with Human TruStain FcX (Biolegend) at a concentration of 5µL per million cells to block unspecific Fc Receptor binding. Single cell suspensions were washed once with FACS Buffer (5% FBS, 0.1% NaN_3_ in D-PBS) and then incubated for 30 minutes at room temperature in the dark with a fluorescence-conjugated antibody cocktail in FACS Buffer (Methods Table 3). The information regarding working dilutions used in this antibody cocktail is in Methods Table 1. The optimal working dilutions were determined by titrations with similar hMOs and experimental conditions. After incubation, single cell suspensions were washed twice with FACS Buffer and resuspended in FACS Buffer. Samples were placed at 4°C until ready to be analyzed by flow cytometry.

In parallel, compensation control staining was performed with the same conditions as the single cell suspensions. The compensation controls used are UltraComp eBeads™ Plus Compensation Beads (ThermoFisher) and ArC™ Amine Reactive Compensation Bead Kit (ThermoFisher) Samples were placed at 4°C until ready to be acquired by flow cytometry.

**Methods Table 3:**
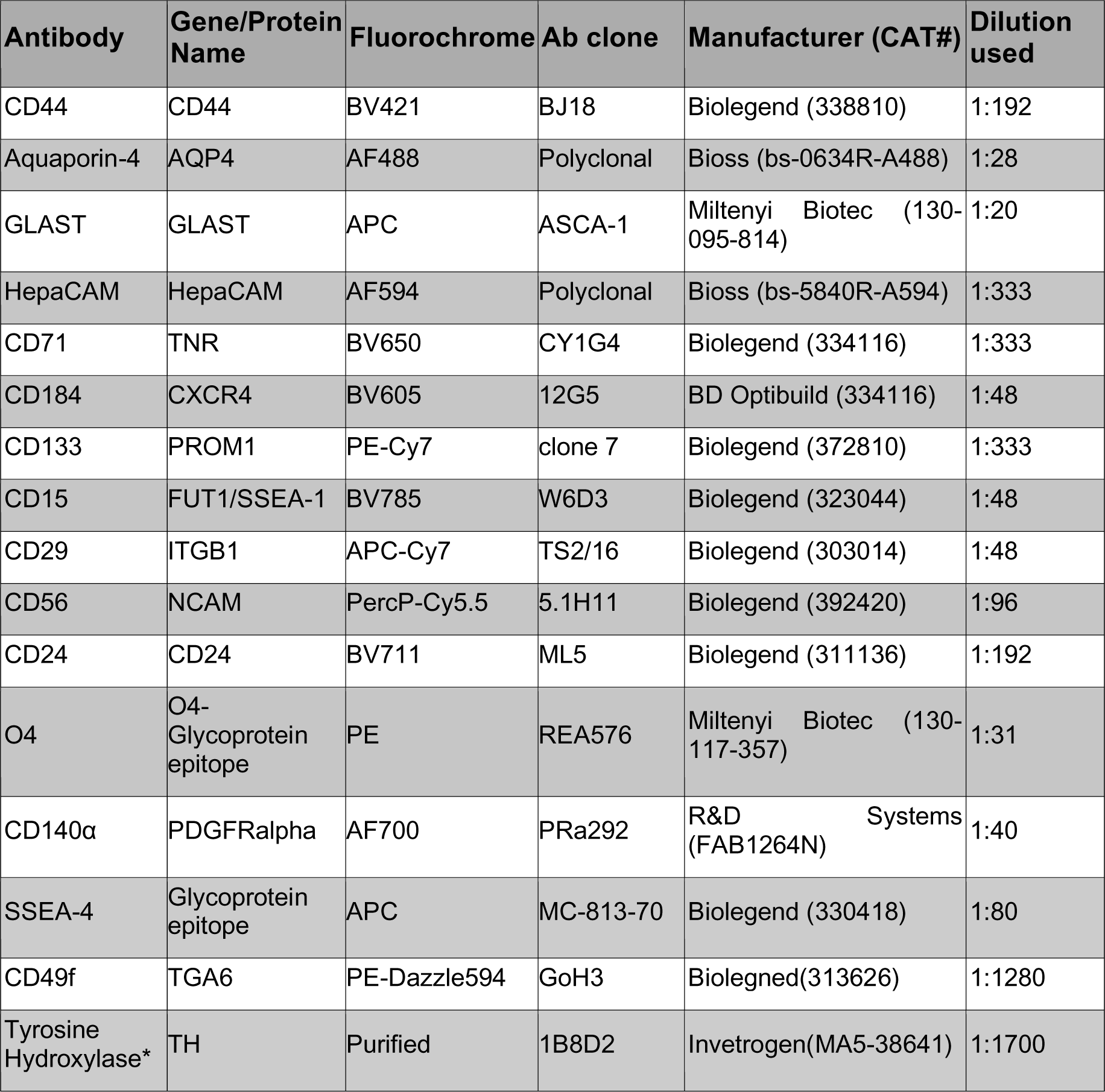
Panel of antibodies and protein targets to measure using Flow Cytometry to identify cell types in hMOs with Fluorochrome and antibody information indicated. *For TH antibody fluorochrome coupling was used following manufacturer’s protocol (Lightning-Link PE, Abcam, CAT# ab102918)

#### 4.5 Fixation and permeabilization for TH internal antibody labelling

Following cell surface antibody staining of the single cell suspension, all cells were washed twice and incubated with 2% PFA diluted in PBS for 15 minutes at room temperature in the dark. After fixation the cells were washed 3 times with FACS buffer and centrifuged at 350g for 5 minutes. Following washes, the cells were permeabilized (0.7% Tween-20 in FACS buffer) for 15 minutes at room temperature in the dark. Cells were washed once (centrifuge 350g 5 minutes) and incubated in the dark for 30 minutes with the fluorescent-conjugated TH antibody. Cells were washed twice in FACS buffer (centrifuged 350g for 5 minutes) and resuspended in FACS buffer for further analysis by flow cytometry.

#### 4.6 Flow Cytometry acquisition – All data sets

Single cell suspensions were acquired on an Attune NxT (ThermoFisher). The information for the configuration of this Flow Cytometer is in Methods Table 2. Daily CS&T performance tracking was done prior to cell acquisition by recommendation of manufacturer. PMT voltages were determined by Daily CS&T performance tracking. Compensation controls were also acquired, creating an acquired compensation matrix. Between 48 000 to 338 000 cells were acquired per sample.

#### 4.6 Flow Cytometry cell sorting defined by CelltypeR workflow – Sorting data set

Single cell suspensions were sorted on a FACSAria Fusion (Becton-Dickinson Biosciences). The information for the configuration of this Flow Cytometer is in Methods Table 2. Daily CS&T performance tracking was done prior to cell acquisition by recommendation of manufacturer. PMT voltages were determined by Daily CS&T performance tracking. Compensation controls were also acquired, creating an acquired compensation matrix.

**Methods Table 4:**
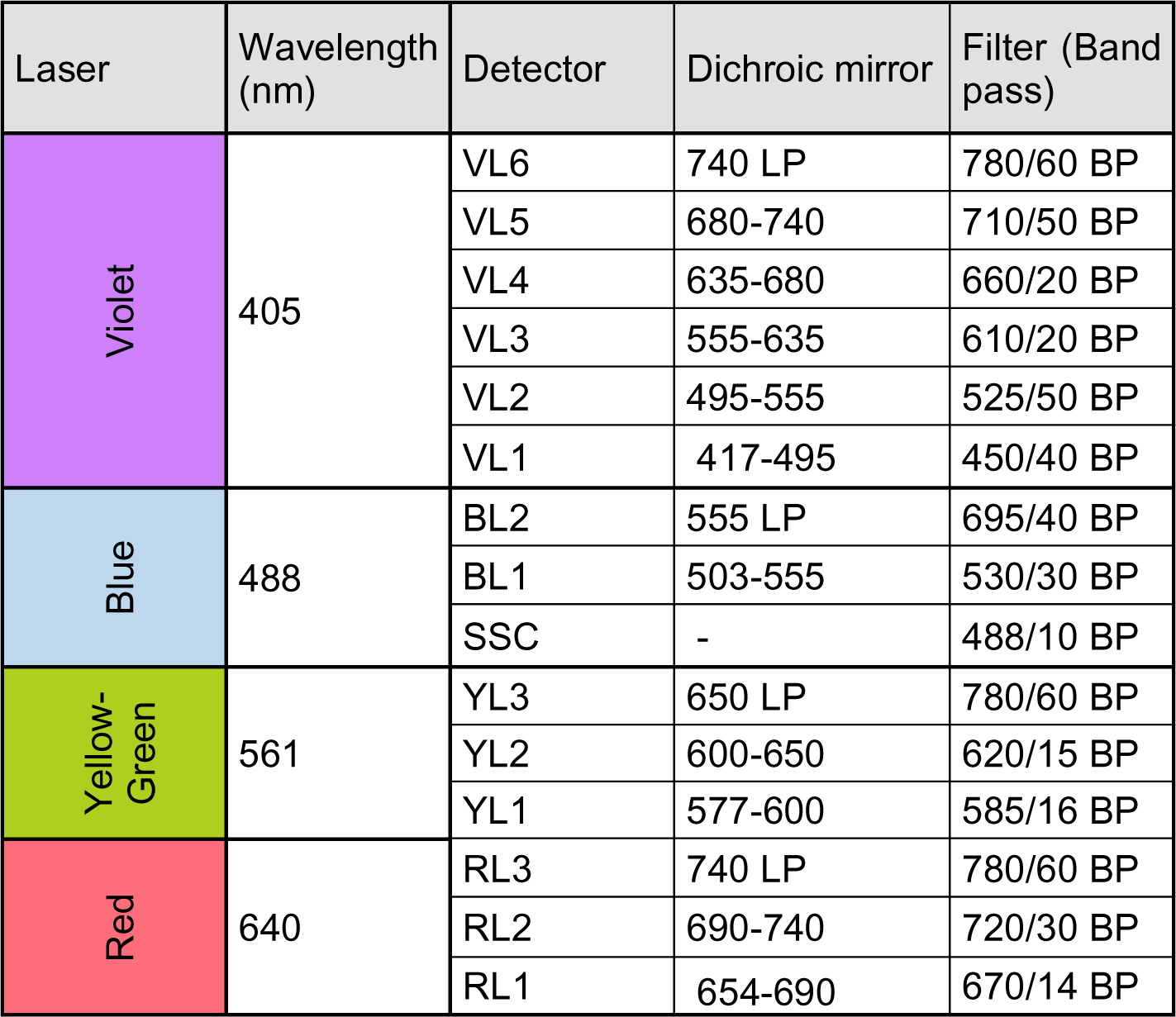
ThermoFisher’s Attune NxT optical path configuration.

**Methods Table 5:**
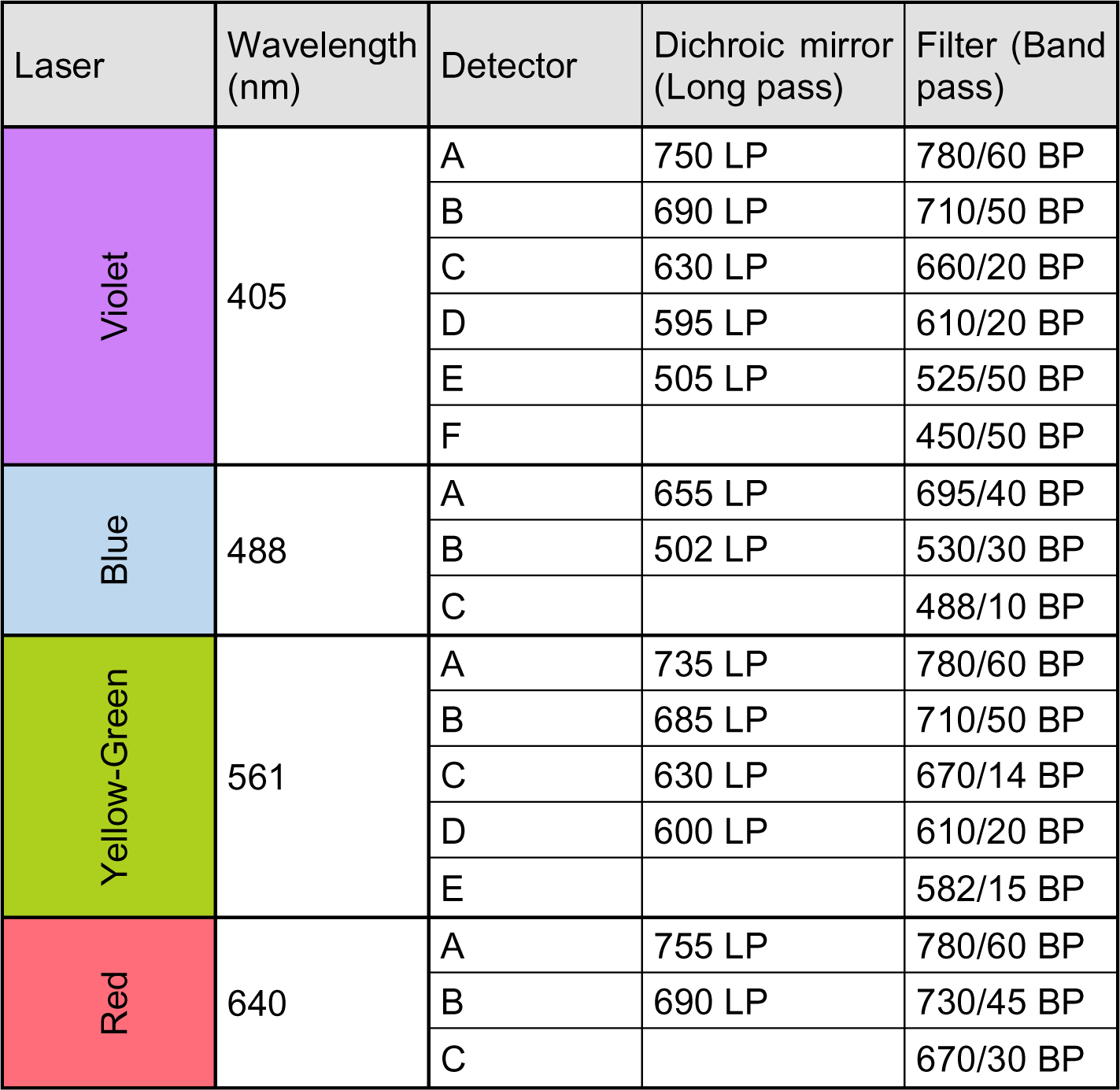
BD’s FACSAria Fusion optical path configuration.

### 5. Single cell sequencing of FACS sorted populations

Three separate tubes of AIW002-02 hMOs were dissociated as described above. At the antibody labelling stage oligonucleotide tagged antibodies (Hashtags, Biolegend) were added with the other cell type specific antibodies. The cells were sorted into FACS buffer. The same sorted populations from each of the three samples (replicates) were combined after sorting. These four populations were sorted into four gates and were sorted until the sample with fewest cells (Neurons1) contained 100,000 events. The sorted samples were centrifuged for 5 minutes at 400g and resuspended in 250 ml of D-PBS + 0.1% BSA. The cell concentrations were calculated with FACSAria Fusion (Becton-Dickinson Biosciences). The single cell suspensions were diluted to 1000 cells/ml targeting ∼15,000 cells captured for sequencing. One sample was prepared for each FACS sorted population.

Following the creation of the cell suspension, the Chromium NextGEM Chip G (PN-1000120) was then loaded as per manufacturer recommendation and run on the Chromium Controller (PN-1000204) for GEM creation. All proceeding thermocycler steps in the 10X protocol were carried out on a Bio-Rad C1000 Touch thermal cycler (1851196). Following GEM-RT incubations, samples were stored at 4°C overnight. Post GEM-RT cleanup and cDNA amplification were carried out per manufacturer protocol. Samples were stored at -20°C until they were processed for library generation. 3’ gene expression and cell surface protein libraries were constructed per manufacturer protocol and stored at -20°C until sequencing submission. 25 mL of each sample library was sent for sequencing at the McGill Genome Centre.

**Methods Table 6:**
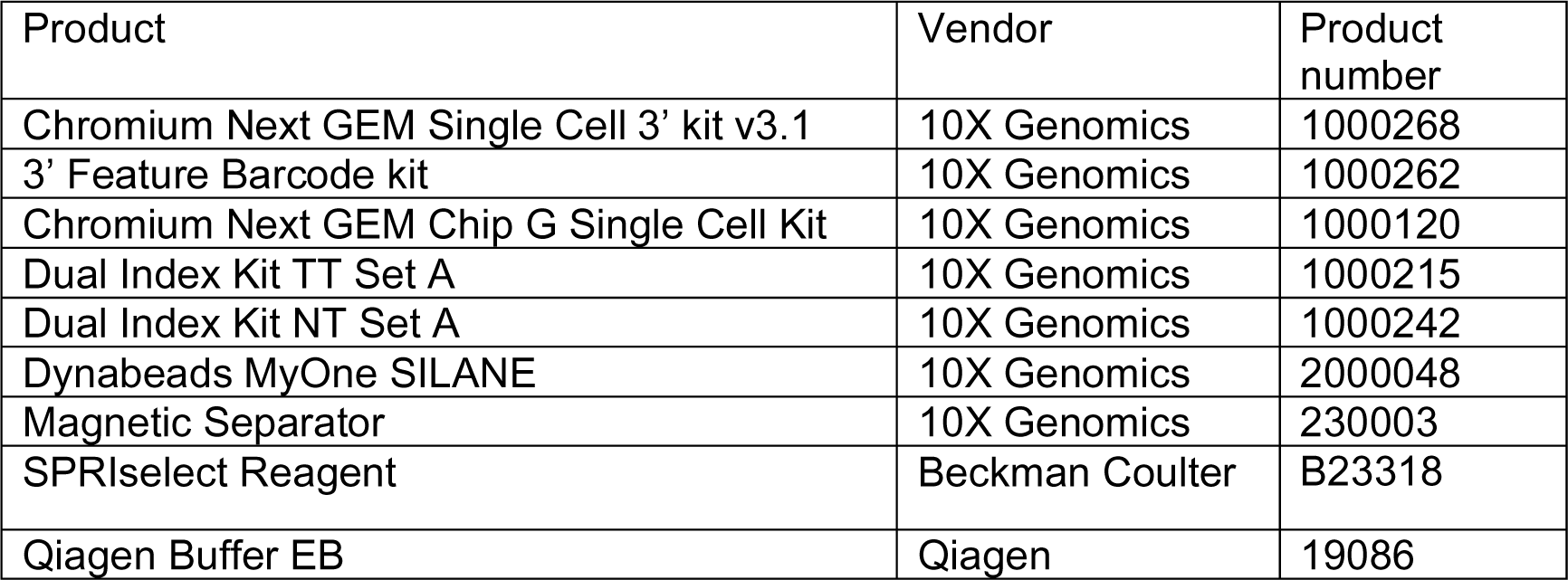
Single cell RNA sequencing reagents.

### 6. Data processing

#### 6.01 Flow Cytometry data cleanup for analysis – All data sets

The data generated was cleaned up using FlowJo (version 10.6) (Becton-Dickinson Biosciences). Briefly, a starting gate was used to select appropriate cell size (X: FSC-A, Y: SSC-A). A second gate was used to discriminate doublets from the analysis (X: FSC-W, Y: FSC-H). Finally, the last gate was used to remove dead cells from the analysis (X: LiveDead Fixable Aqua, Y: FCS-A). See Methods Figure 1 for a gating example. After data cleanup, a new .fcs file was generated with FlowJo and exported for further analysis done with R.

#### 6.02 Data analysis and CelltypeR R library

All computations were performed in R. We created a R library of functions to perform the analysis, CelltypeR. Our functions required functions from multiple other R libraries referenced in descriptions to follow.

The R library can be found, along with workbooks for the complete workflow and generation of each figure, at https://github.com/RhalenaThomas/CelltypeR.

Computational Workflow:

1. Data preprocessing:

a. Read FlowJo files into R.
b. Create a data frame with intensity measurements for each marker for all samples within the experiment to be analyzed.
c. Harmonize data if desired.
d. Create a Seurat single cell object for further analysis.
2. Creation of cell type clusters

a. Clustering optimization to compare clustering methods and parameters and visualize results.
b. Summarize statistics to compare clustering methods and parameters.
c. Select one method and smaller parameter space to compare cluster stability.
d. Evaluate statistics and visualization to determine the best clustering method for a given visualization.
3. Cluster annotation

a. For first data set: marker visualization and correlation assignment model.
b. For subsequent data sets: marker visualization and correlation assignment model, Random Forest Model, Seurat Label Transfer.
4. Quantify cell types and measure expression levels of markers within cell types.
5. Define marker levels for cell types.
6. Statistical analysis between different groups of interest.

#### 6.03 Data preprocessing

The .fcs files without dead cells, debris, and doublets created in FlowJo are read into R and processed. The .fsc files contain area, width, and height of the fluorescence signal for each marker as well as the forward and side scatter of the light. Then R using the flowCore package is used (Hahne et al., 2009). The area values for each channel are selected to represent the expression intensity for each antibody. All the .fsc files within one folder are read into into one R data object. A dataframe is created with the channels and saved for further use. Individual cell cultures and hMO organoid samples for testing the pipeline and gating were used in this raw format to create a Seurat single cell data object.

For the hMO samples, the data was aligned to remove batch effects and technical variability. Each file represents an experimental sample, and the samples were aligned as follows: First, to enhance the distinction between positive and negative antibody staining the raw data is transformed using the biexponential transform function from flowCore with default parameters (*a=0.5, b=1, c=0.5, d=1, f=0, w=0).* The transformed data was visually inspected to confirm there were no errors (see R workbooks on Github). To combine the nine different hMO samples and account for batch effects, the signals were aligned using an unbiased approach, the gaussNorm function in flowStats (Hahne et al., 2013). Local maxima are detected above the bandwidth we set to be above 0.05, to avoid picking up noise, each peak is given a confidence score reflecting the height and sharpness of the peak, the threshold for two peaks to be considered too close together was set too 0.05. Landmarks are then detected and aligned, such that each landmark is shifted to a benchmark, which corresponds to the position of the closest peaks across all samples. After alignment the data is reverse transformed to improve visualization by UMAP in downstream analysis.

#### 6.04 Creation of cell type clusters

For the analysis in Figure 3, to test cluster methods and cell type annotation methods, we selected a subset of hMO cells. From 8 of the hMO 9000 cells were randomly selected and one sample all the cells (1578) cells were selected before transformation and alignment. We compared FlowSom^44^, Phenograph^45^ and the Seurat^46^ Louvain network detection function as well as parameter space (k neighbours, resolution, k clusters) available for the different algorithms. We calculated intrinsic statistics and produced UMAPs and heatmaps for visualization. We found FlowSom was not suitable for creating clusters based on cell types, although the intrinsic statistics are best for FlowSom. Phenograph uses the Louvain network detection method and computes the Jaccard coefficient which considers the number of common neighbours between cells. Phenograph functions well, however we saw little difference to the Louvain using the Seurat library and proceeded to use the Seurat package for Louvain network detection to obtain clusters for ease of use with the overall workflow. We then proceeded to test the cluster stability at different resolutions, calculating the RAND Index and standard deviation of the number of clusters across 100 iterations of clustering with different random start points. The results informed the choice of cluster numbers to annotate.

#### 6.05 Cluster annotation

Cell type annotation was performed on the subset of 9000 cells using visualization and a correlation assignment model (CAM) we created. For visualization we created functions to make UMAPs for expression levels of each antigen targeted in the antibody panel as well as heatmaps grouped by cluster numbers. The expended expression patterns were of the antibodies as used in combination with the CAM predictions. For the 2D cultures in Figure 2, cell types were assigned by the visualization of expression values, the known original cell type, and the overlap in space on the UMAP. In our culture system iPSC can become any cell type, NPCs are precursor cells for all three other cell types included (astrocytes, DA neurons, and oligodendrocytes). The NPC cultures are multipotent but will contain cells that are beginning lineage selection and those retaining a multipotent state.

For the full hMO dataset of nine samples and the follow up hMO datasets used for gating and sorting experiments, a Random Forest Model trained on the subset hMO data and Seurat transfer labels predictions were used in addition to the CAM and visualization methods used on the subset data. The combined results of methods are more reliable than each method alone. Each of the four methods of annotation are input into the cluster annotate function to automate the cluster annotation process.

#### 6.6 Creation of the predicted expression matrix for antigen proteins in the antibody panel and the new time course antibody panel

Expression values for expected cell type were combined from several sources (Methods Table 7). Microglia are found in brain tissue but are not expected to be present in hMOs due to their mesodermal lineage and thus were not included in the reference matrix. In the first (13 antibody panel) there is not a specific DA marker and so we only define NPCs and neurons. The DA subtype is specified in the second (time course) panel reference matrix. For the 2D cell cultures we collected FC expression values for the 13-antibody panel. For the TH time course panel, we could only use the overlapping markers for the FC expression. For RNA seq data we used the gene expression equivalents to each marker in the two panels. Not all gene equivalents for the protein markers were available from all cell types or databases. For the antibody O4, the epitope is a glycoprotein, and the specific corresponding gene is unknown, however the gene NKX6.2 is a marker of mature oligodendrocytes, with expression highly correlated to O4 protein detection.^47^ For SSEA-4, another glycoprotein epitope we used gene encoding the SSEA4 synthase enzyme

We used total RNA and scRNAseq from adult human brain as well as scRNAseq data from organoids. For each expression data set the values were z-scored and normalized between 0-1. The mean expression form human brain and organoids were calculated and normalized. Then the mean values from 2D culture FC, brain RNA expression and organoid RNA expression were calculated and then normalized for the final matrix. Details for each step and how to make a qualitative reference matrix can be found in the R notebook *“CreateReferenceMatrix.Rmd”*.

Inputs were taken from bulk RNAseq from human brain tissue^48^, scRNAseq data from human fetal midbrain^49^ and adult midbrain^50^, from developing human cortex and forebrain, and cerebral organoids^51^. Finally, the FC data acquired in this study from 2D cell cultures, iPSC, neural precursor cells, neurons, astrocytes, and oligodendrocytes. For each data set the values were z-scored then minmax normalized marker by marker to fit between 0 and 1. The mean expression values were calculated separately for scRNAseq organoid data and scRNAseq brain data. Then the mean expression values were then calculated between scRNAseq-hMO, scRNAseq-Brain, and RNAseq. Then the mean of that result was calculated with the FC data. The FC data was weighted more highly than the public data sets because it is experimental data collected on protein levels with the exact antibodies used for hMO experiments; however, we didn’t generate data on all possible cell types. The predicted expression values were again z-scored then minmax normalized marker by marker to fit between 0 and 1 to be comparable to the transformed FC data to be used in the correlation assignment model.

**Methods Table 7:**
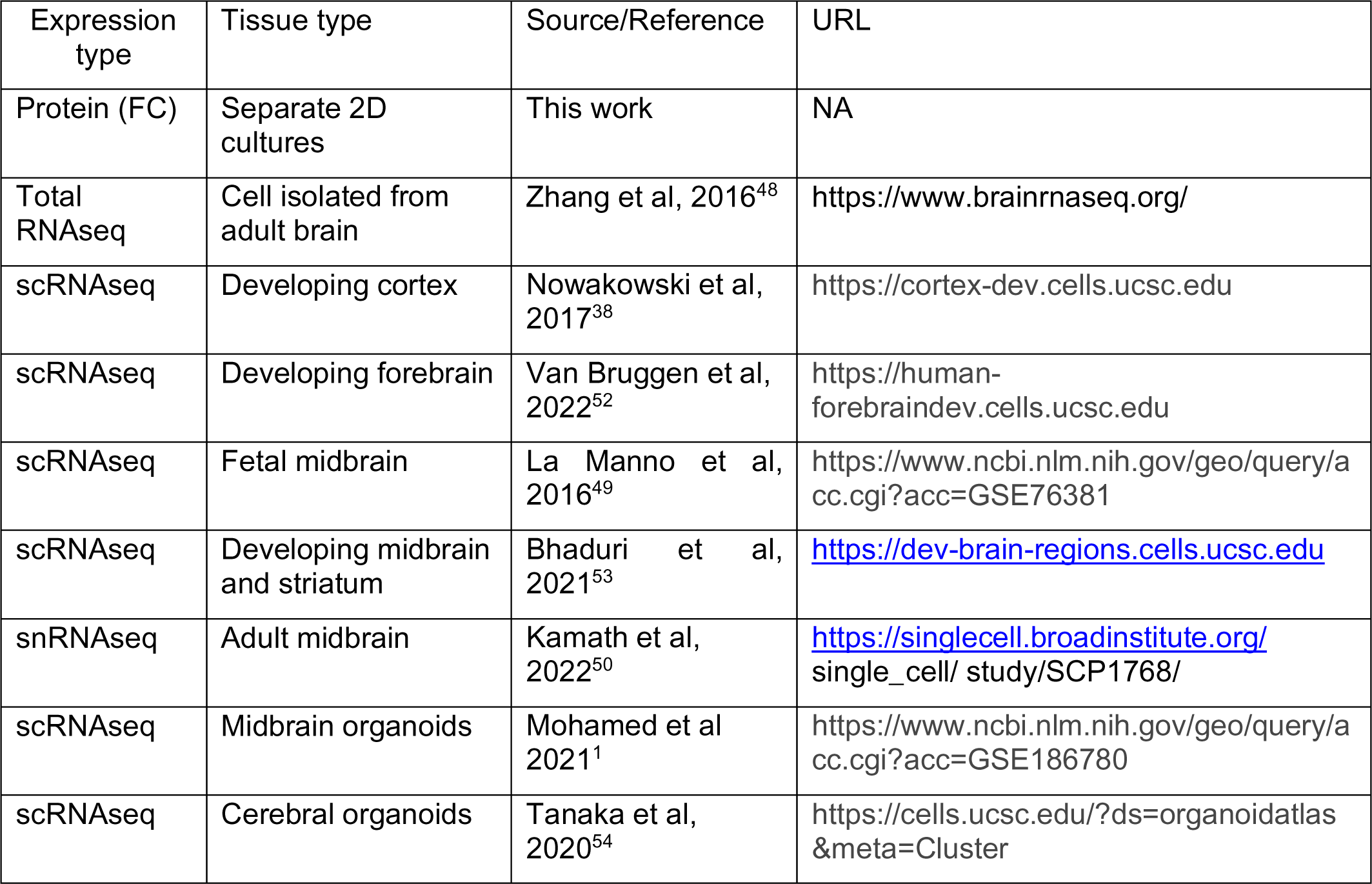
Sources of expression values used for reference matrices.

#### 6.07 Assigning cell type labels to clusters using correlation to the predicted expression matrix

Pearson correlation coefficient R values, were assigned to each cell, correlating the FC intensity expression levels of antibody panel to the predicted expression values in the reference matrix for each cell type expected in the hMO. The R values were calculated for each potential cell type. Then for each hMO cell, the max R value and the second max R value were selected. These values were then used to predict the cell type for each hMO cell. A threshold was set of R > 0.45 for a cell type to be predicted, otherwise the cell is assigned as ‘unknown’. If the Rmax1 – Rmax2 < 0.05, then a mixed cell type is assigned. For example, Neuron-NPC. For the cluster annotations, the top three most frequently predicted cell types for each cluster were calculated. If most of cells were predicted within a cluster as one cell type, this cell type was assigned to the cluster. If the frequencies of predicted cell types were distributed across many cell types, the cluster was assigned as mixed or unknown.

#### 6.08 Random Forest Model

A data frame was created from cell type from the 9000 cells per sample subset of hMO data and the matching expression. The data was split 50/50 into test and training data. The training data was input into the function RFM_train which uses the *randomForest* package and the *caret* package for optimization. A range of the number of variables (antibodies) was randomly sampled in each split (mtry) from 1 to 10, and the best mtry was 6. Ranges of other parameters were tested, and the optimal values were used in to train the final model: max nodes = 30, starting node size = 25 and number of trees = 1000. The trained model was then used to predict the cell type of each cell in the full data set and the new flow sorted data. The topmost predicted cell type for each cluster was used as the cluster annotation prediction.

#### 6.09 Label transfer using Seurat

We made a function that follows the Seurat workflow for label transfer combined into one function. The annotated Seurat object from the 9000 cells per sample subset of hMO data was used as the reference data and the full dataset and FACS datasets were used as the query objects. Anchors were found between the two objects using 25 principal components to predict the cell types, the max prediction was selected for each cell in the query data. No threshold for predictions was set. The most frequently predicted cell types within each cluster were used as the cluster predictions.

#### 6.10 Quantification of cell types and statistical analysis

Permutation tests were used to determine if the changes in cell type proportions were significant between two groups using the R library *scProportionTest* (https://github.com/rpolicastro/scProportionTest). In the permutation test, sample labels are repeatedly shuffled and the log2-fold is calculated change in fraction for each shuffle to create a null distribution of log2-fold changes. This distribution represents what we’d expect to see if there were no real differences between the two samples. Next, p-values are calculated for each cluster (cell type) by comparing how many times the permuted log2-fold changes are as extreme as the observed log2-fold change. These p-values are adjusted for multiple comparisons using the false discovery rate (FDR) method, providing a measure of the statistical significance of the observed differences in proportions. Permutations tests were selected over the commonly used Chi square or Fisher’s exact test because tests assume the observation are independent. However, the proportion of the cell types within a sample is directly dependent on the other cell types because the confounding these tests.

Two-way ANOVAs and Tukey’s pos hoc tests for main effects and interactions were all run using functions in our R library. A preprocessing function is used to pull the expression data out of the Seurat object and add the desired variables. The statistic functions used the base R functions *aov* and *TukeyHSD*. The effect of each variable was analyzed separately. A loop was used to analyze each cell type separately.

**Methods Table 8:**
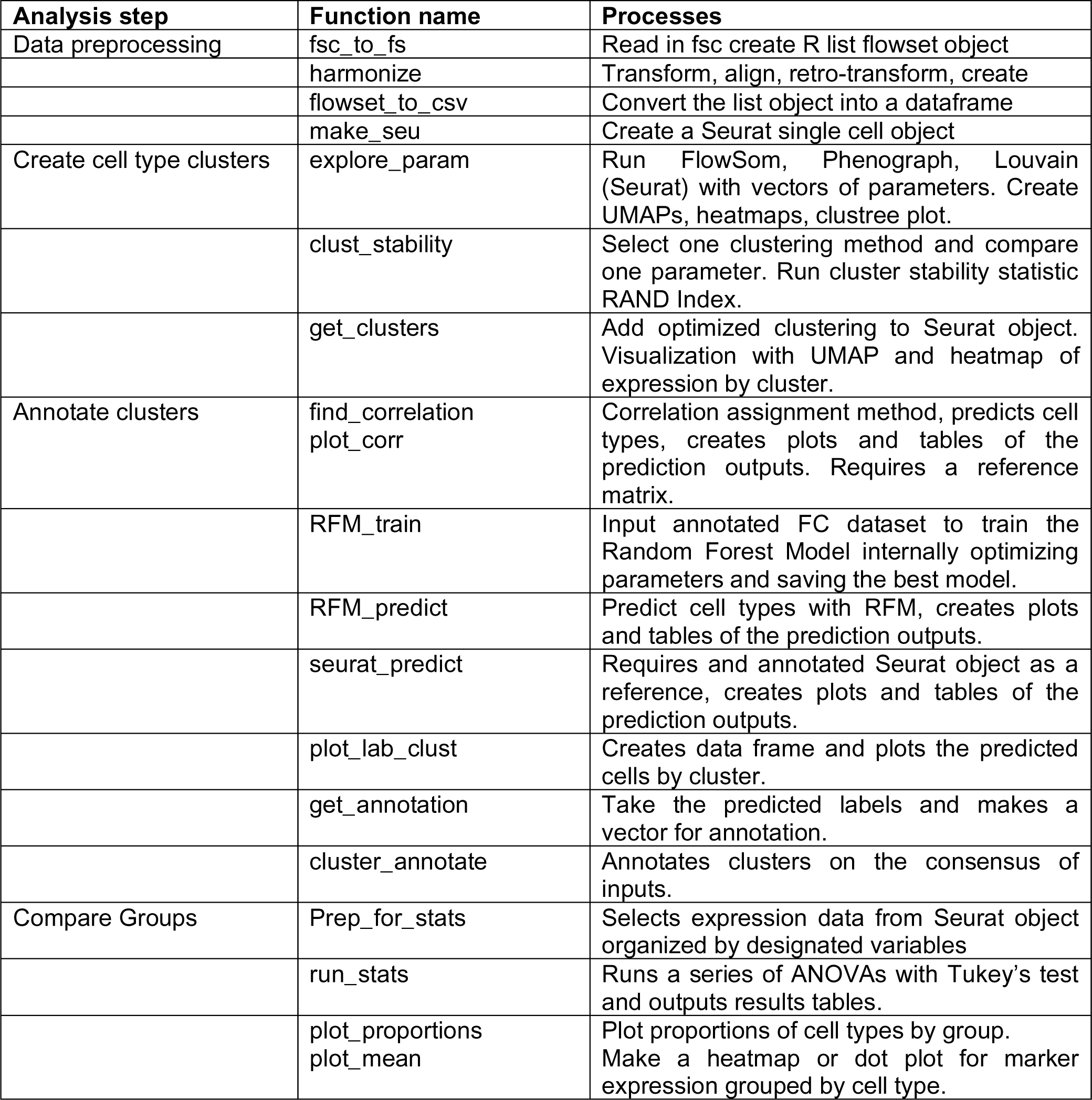
Function names in the CelltypeR library for analysis processes.

#### 6.11 Testing gates reverse engineered using *hypergate*

Cell types were selected in the full annotated hMO dataset and input into the *hypergate* function.^39^ A table of predictions was output. For each cell type the threshold levels for each antibody required to define the cell type were output. These thresholds are in order from most to least important. For testing the gates, manual gating was applied in FlowJo with the top gate for each cell type in each sample being set as live single cells. The gates were applied in an AIW002-02 sample and then applied across the other samples. For gating, the two antibodies were visualized by scatter plot and a box was drawn selecting the thresholded cells from the antibody pair. The gated cells were then selected and gated with the next pair of antibodies until all thresholds were applied. The final gated cell types from all samples were exported as fsc files and read into R following the CelltypeR workflow. To apply gates to FACS four selected samples examined each cell type gate and selected gates which mostly exclusive for different cell types. The neurons can be separated from glia and then split into two populations and the glia can be split into two populations.

#### 6.12 Single cell sequencing analysis

The FASTQ files processed using 10X CellRanger 5.0.1 software are installed on the Digital Research Alliance of Canada: Beluga computing cluster. For each of the four sorted populations, the CellRanger output files raw expression matrix, barcode, and feature files were used to create a Seurat data object with minimum filtering of RNA features > 100. After this point data was run locally and all details can be found in the R notebook, ‘scRNAseq_processing’. RNA features, RNA counts, and percent mitochondria were checked for quality control for each sample: Neurons1, Neurons2, Glia1(astrocytes), and Glia2 (radial glia). Further filters were applied.

**Methods Table 9:**
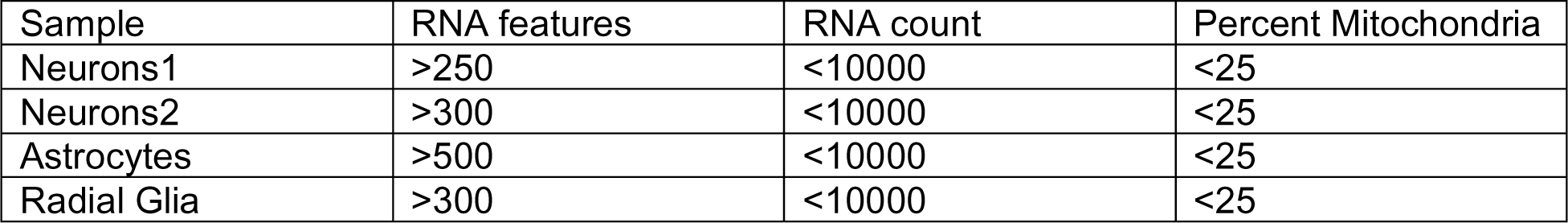
Filters applied to each scRNA population.

For the glia samples, there were a large number of cells after filtering. The Seurat function HTODemux was used to assign Hashtag (replicate labels). For neuron samples and radial glia all cells were selected, for glia1/astrocyte sample the original count was very high. To increase selection of true cells, cells with assigned hashtags were used for further processing. For all samples, doublets were removed using Doublet Finder^55^. The expected percent of doublets estimation was based on the number of cells present after filtering and the 10X version 3 user guide. For each sample data was normalized, variable features selected, PCA and UMAP dimensional reductions were performed, and clusters detected with Louvain network detection (25 dimensions and 43 neighbours selected, and a range of resolutions was run).

Clusters were annotated using a consensus between expression of known cell type markers from gene lists, analysis of cluster markers, and cell type predictions of reference data (see below) using Seurat find anchors and label transfer. Subtypes of major cell type groups were observed and at this point these clusters were all merged into major cell types. The individually processed samples were then merged, samples were down sampled to balance the data and decrease processing time.

**Methods Table 10:**
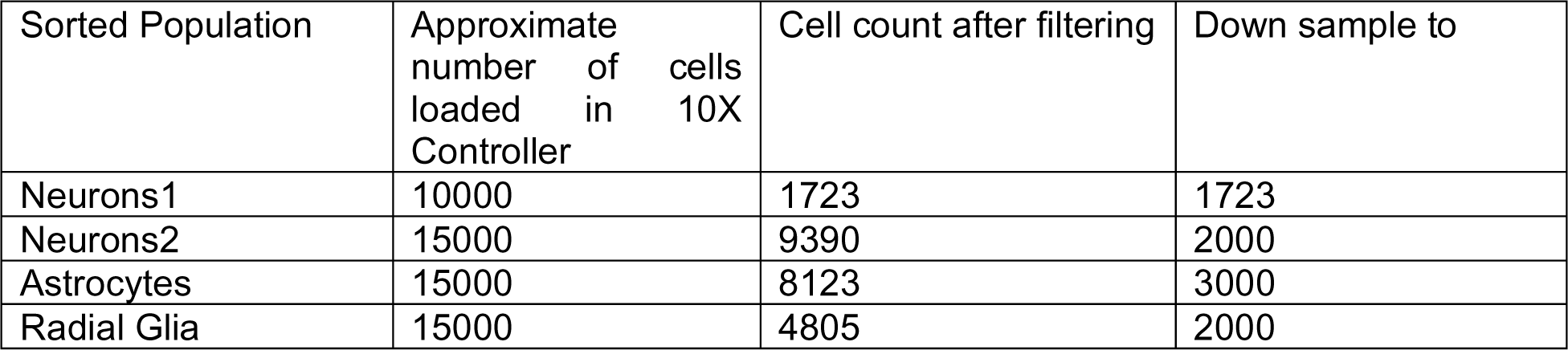
Cell counts after filtering and counts used in merged data object.

After the four samples were merged the standard processing and clustering was run again using the same settings. Clusters were annotated again, retaining subtypes of each cell type, and identifying the DA neurons. Each subtype was analyzed to find subtype markers and analyze using GO biological processes. Reference datasets using Seurat anchors and label transfer predictions were used to define subtypes of cells. All the scRNAseq data sets in Methods Table 7 were used. A threshold for cell type assignment was set to 0.5 for brain reference data and 0.8 for hMO scRNAseq data.

Developing cortex, forebrain and whole brain datasets were all reconstructed into Seurat objects from the UCSC cell browser following the website instructions.^56^ Each reference was down sampled in Seurat to reduce the total cell number to less than 50000.

For single nuclear RNAseq data from human adult postmortem brains (Kamath et al) three separate reference sets were created. The expression matrix, barcodes, and feature files were used to create a Seurat object. The metadata for cell type and cell subtype annotations data was added from the UMAP_tsv files provided by Kamath et al. The brain region data was added from the provided meta data file. The adult midbrain was subset by brain region selecting only the midbrain cells. The DA subtypes and astrocyte subtypes were separately subset by using the main cell type annotation.

1. All cell types (astrocytes, oligodendrocytes, microglia, endothelial cells, DA neurons and other neurons). This was used in the initial cell type annotations.
2. DA neuron subtypes, used to try to identify DA subtypes. All the hMO subtypes matched only one subtype from adult brain.
3. Astrocyte subtypes, used to identify astrocyte subtypes. All astrocyte subtypes in hMO matched one subtype.

After annotating the main groups of cell types (DA neurons, neurons, astrocytes, radial glia, NPCs, mixed) subtype annotations were applied. To annotated subtypes, the main cell type was subset. The *Seurat* find all markers function was used allowing both up and down regulated gene markers of the clusters within each main cell type. The top 5-10 marker genes sorted by highest Log2 Fold change with significant adjusted p-values were further investigated by literature search to determine the cell subtypes.

### 7. Data availability

Flow cytometry: FlowJo selected live gated cells in the form of .fsc files are available in the github repository: https://github.com/RhalenaThomas/CelltypeR.

Cell type annotation: The expression matrix for the proteins/genes targeted by the antibody panel to run CAM and a trained RFM are in the github repository in the data folder.

scRNAseq: The FASTQ files, CellRanger filtered outputs, processed annotated Seurat R objects for each of the four sorted population and the combined processed annotated Seurat data object are all deposited in the NCBI Gene Expression Omnibus (GEO) under the accession number GSE226890. https://www.ncbi.nlm.nih.gov/geo/query/acc.cgi?acc=GSE226890.

### 8. Code availability

All code is available on github: https://github.com/RhalenaThomas/CelltypeR

The repository includes:

1. R library *CelltypeR* containing all functions in Methods Table 5.
2. Code for the *CelltypeR* library functions.
3. Workbook with the analysis workflow and application of *CelltypeR* functions.
4. Workbooks with the code used to generate the figures.

## Acknowledgments

E.A.F. is supported by a CIHR Foundation grant (FDN – 154301), a Fonds d’Accéleration des Collaborations en Santé (FACS) grant from CQDM/MEI and by a Canada Research Chair (Tier 1) in Parkinson’s disease.

T.M.D. received funding through the McGill Healthy Brains for Healthy Lives (HBHL) initiative, the CQDM Quantum Leaps program with support from Brain Canada, the Alain and Sandra Bouchard Foundation, the Sebastien and Ghislaine Van Berkom Foundation, Médicament Québec and the Mowafaghian Foundation. T.M.D is supported by a project grant from CIHR (PJT-169095).

R.A.T. received funding through the McGill Healthy Brains for Healthy Lives (HBHL) Postdoctoral Fellowship and Molson NeuroEngineering Fellowship. The authors thank David Kalaydjian and Nguyen-Vi Mohamed for helping to test early midbrain organoid differentiation protocols. We thank Dr. Jo Anne Stratton for use of the 10X chromium controller and for constructive feedback on the manuscript.

## Contributions

The project was conceived by RAT and JS. The production and maintenance of hMOs was done by MM and PL. FACS, panel optimization, antibody titration and FC acquisition were done by JS. Sample preparation for flow acquisition, sorting and scRNAseq were performed by JS, RAT, and PL. The iPSC, NPC and DA neuron cultures and corresponding IF were by done by CXQC. Astrocyte cultures and IF by VS. Oligodendrocyte and OPC cultures and IF by VEP. The hMO cryosections were prepared by PL and the IF was done by VEP and GED. The scRNAseq library preparation was done by TMG and LF. FlowJo analysis was performed by JS and RAT. Computational workflow and the R library were designed, tested, and managed by RAT. R functions were written by AG, SL and RAT. Computational analysis was performed by RAT. The CelltypeR library was tested by GED and RAT. Data was interpretated by RAT, TMD and EAF. Project was supervised by RAT, TMD and EAF. The manuscript was written by RAT with contributions by JS, GED,TMG, EAF, TMD.

## Supplemental Figures

**Figure S1:**
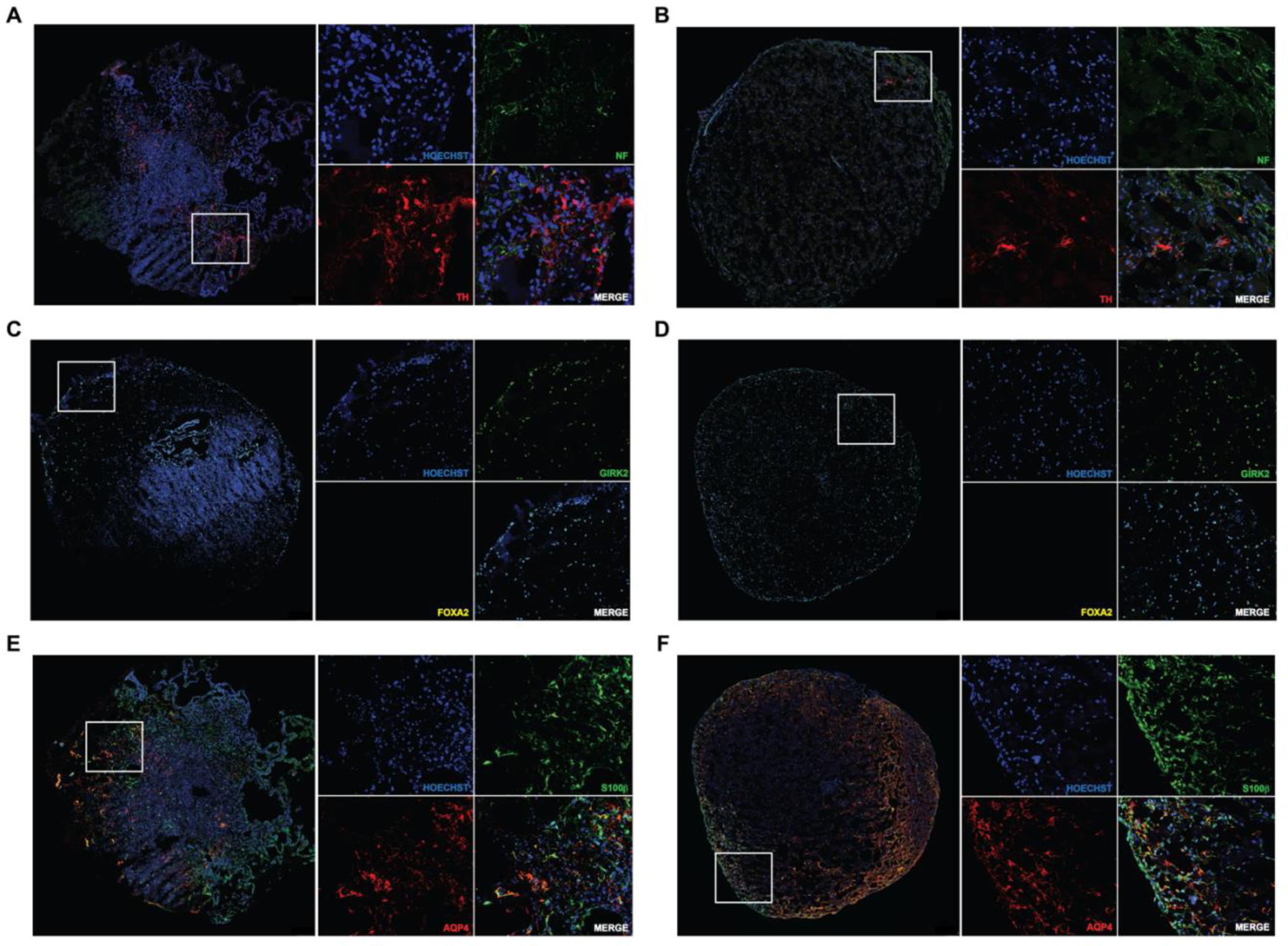
Example images with characterization of cell types in hMO 9 months in culture. Cryosection with immunostaining of iPSC lines AIW002-02 (left) and AJG001C (right). Whole organoid sections with merged signals are shown in the left panel, the area in the white rectangles is enlarged on the left with each staining shown separately. Nuclei are labelled with Hoechst and shown in blue. **A, B)** Tyrosine hydroxylase (TH) marker of DA neurons in red and neurofilament (NF) a general neuronal marker in green, indicate DA neurons are present in hMOs from both iPSC lines. **C,D)** FOXA2 a neural progenitor marker of DA neuron lineage is absent at this late stage (would be yellow). GIRK2 (green) is expressed in DA neurons of the SN and is present in the hMOs. **E,F)** Astrocyte markers S100β (green) and AQP4 (red) indicate astrocytes are present in the hMOs.

**Figure S2:**
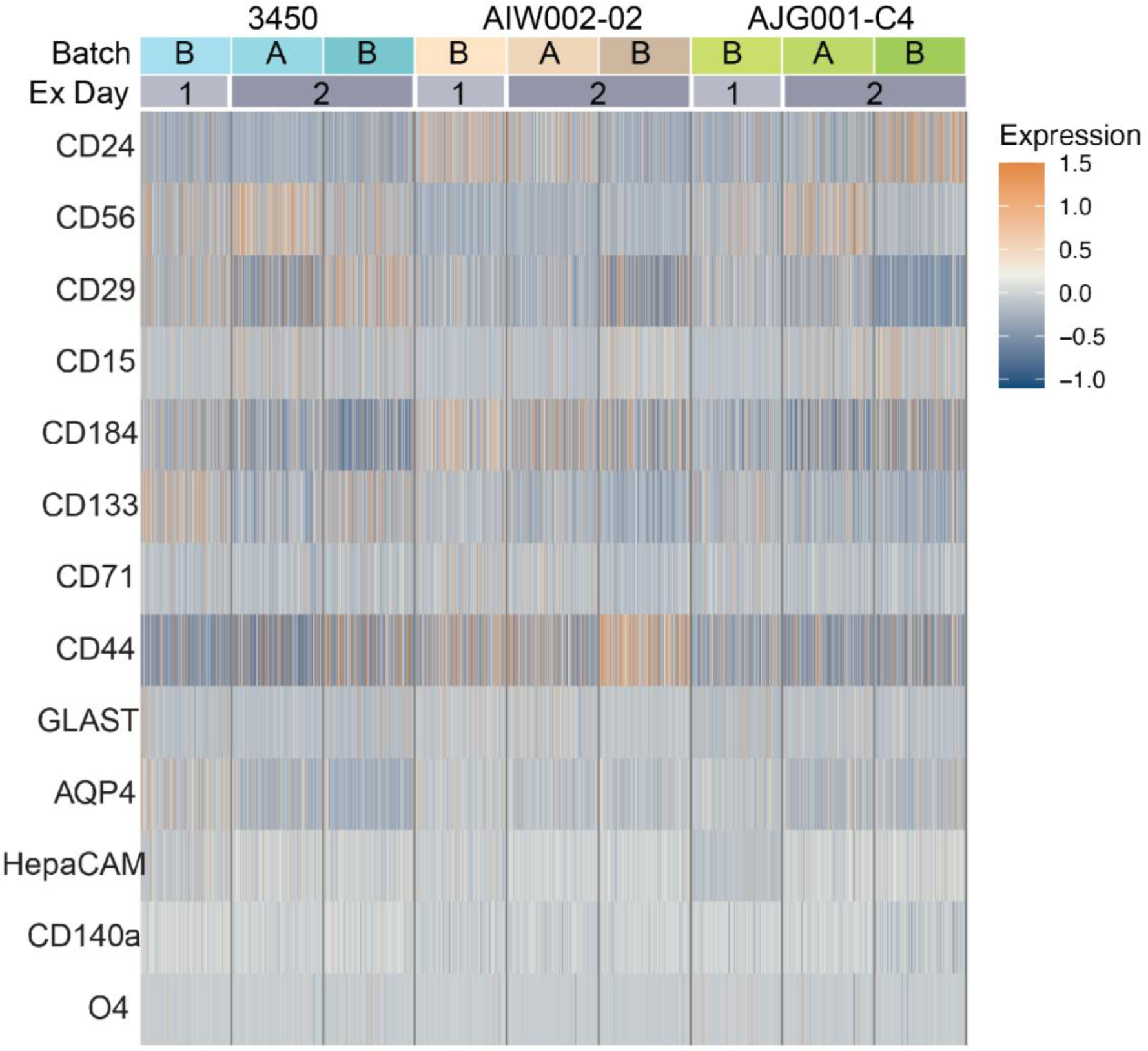
Expression of the 13 protein markers varies across cells in hMOs. Protein expression levels measured by FC in a subset of cells from 9 different hMO samples. The three iPSC lines (3450, AIW002-02-02, AJG001-C4, two batches (A = 20190620, B = 20190530) and two different experiment days (1 = 06/03/2020, 2 = 17/03/2020) are indicated at the top.

**Figure S3:**
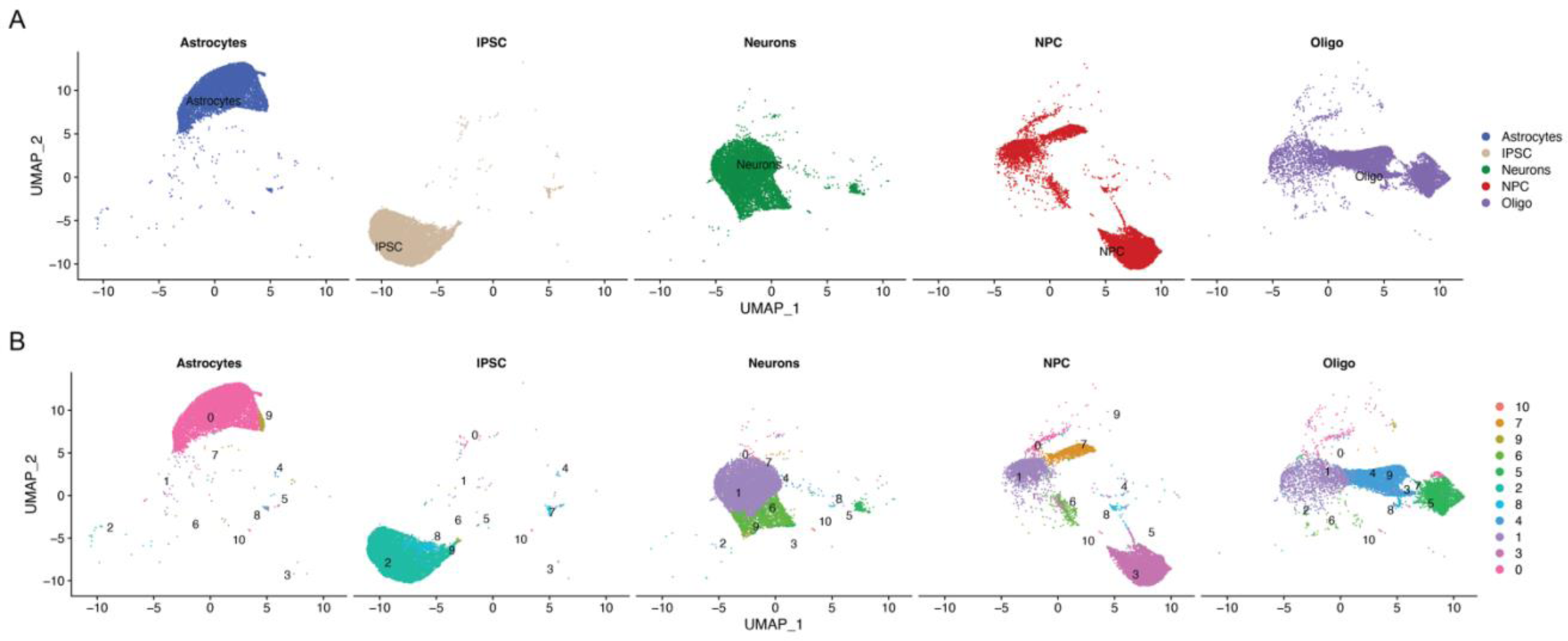
UMAPs of 2D cell cultures and clusters. A) UMAP showing cell split by the original culture type and coloured by the original culture type. B) UMAP of cell split by the original culture type and colour by the clusters identified by unsupervised Louvain network detection.

**Figure S4:**
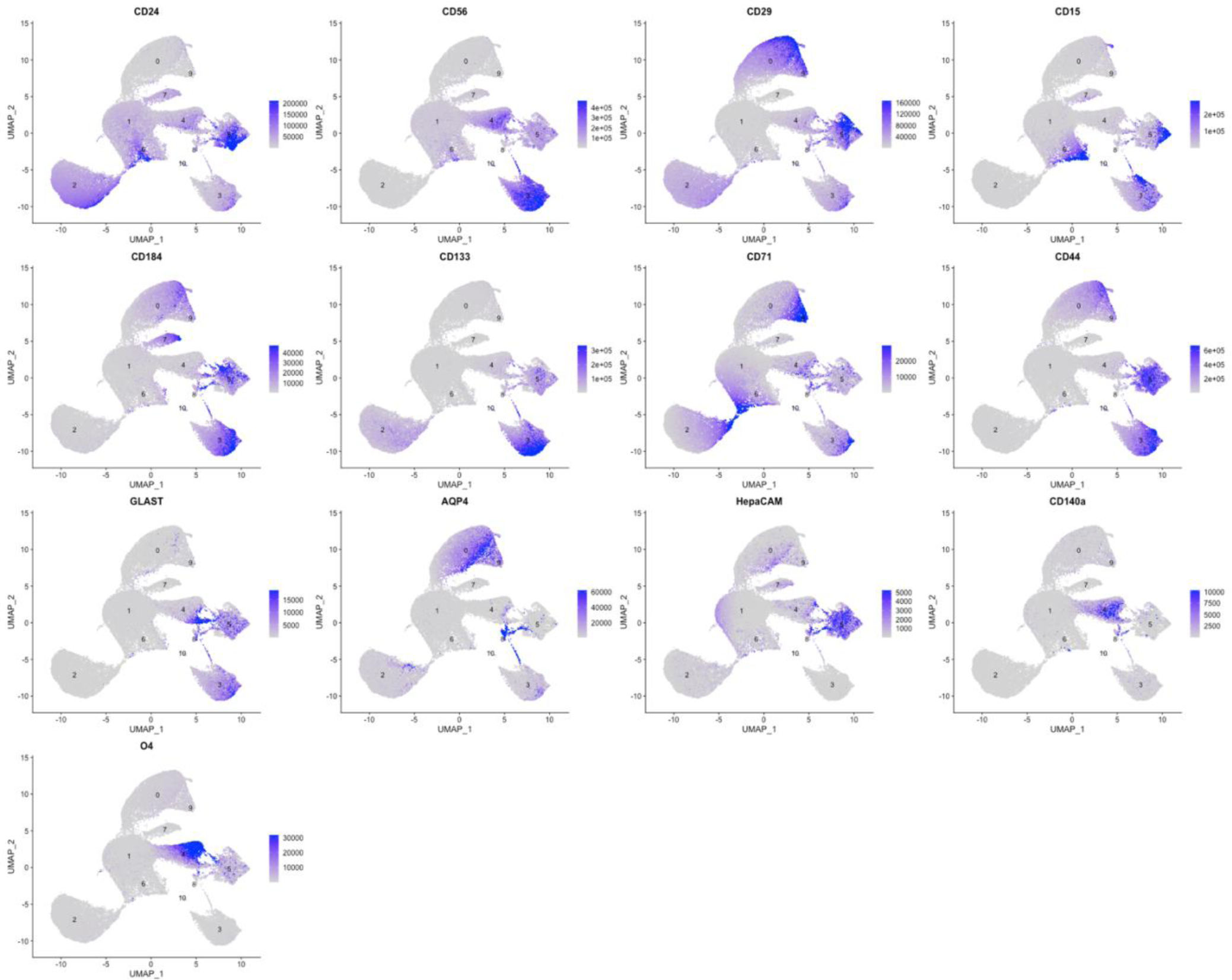
UMAPs of 2D cell cultures with clusters labelled. The marker expression measured is labelled above each plot.

**Figure S5:**
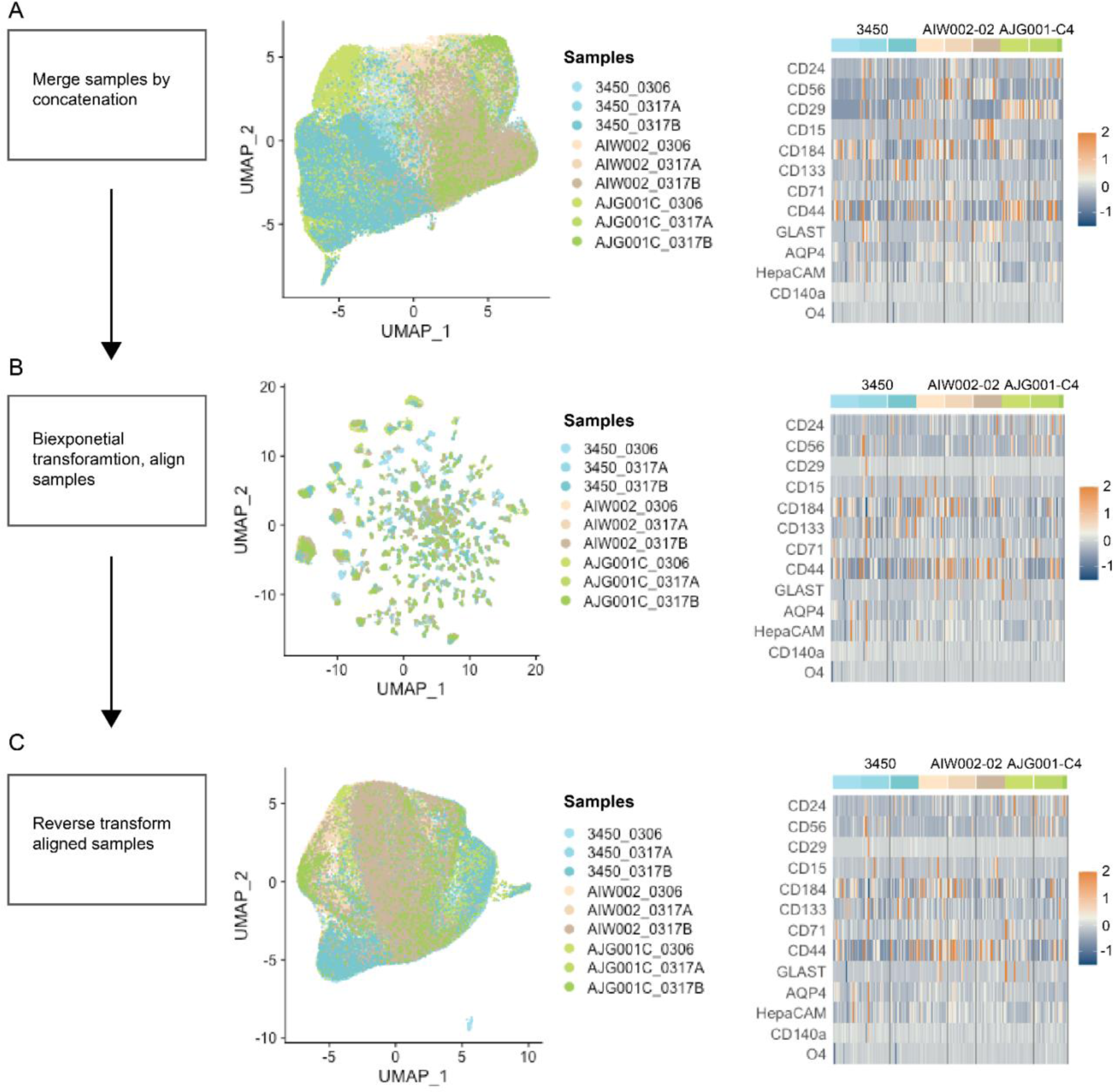
Preprocessing of fsc files exported from FlowJo and read into R. **A)** Samples are merged by concatenation and not transformed or aligned in preprocessing. Files can be saved at this level of processing and one can proceed with the rest of the CelltypeR workflow if desired. For individual 2D iPSC derived cell lines, processing was stopped at this step. **B)** The merged expression data is biexponentially transformed and aligned by shifting means to match peaks between samples. **C)** The merged, biexponentially transformed, and aligned data is reverse transformed removing the biexponential transformation. The full processing was applied for hMO samples to remove experimental variability. Left panel indicates the data processing performed. Center panel indicates UMAPS to visualize hMO cell samples indicated by colour in the sample legend. Seurat was used to scale before PCA and UMAP dimensional reductions. Right panel indicates heatmap of the marker expression levels across each sample. Scale bars indicate the relative expression and are matched in the heatmap and UMAP plots. Normalized expression is plotted.

**Figure S6:**
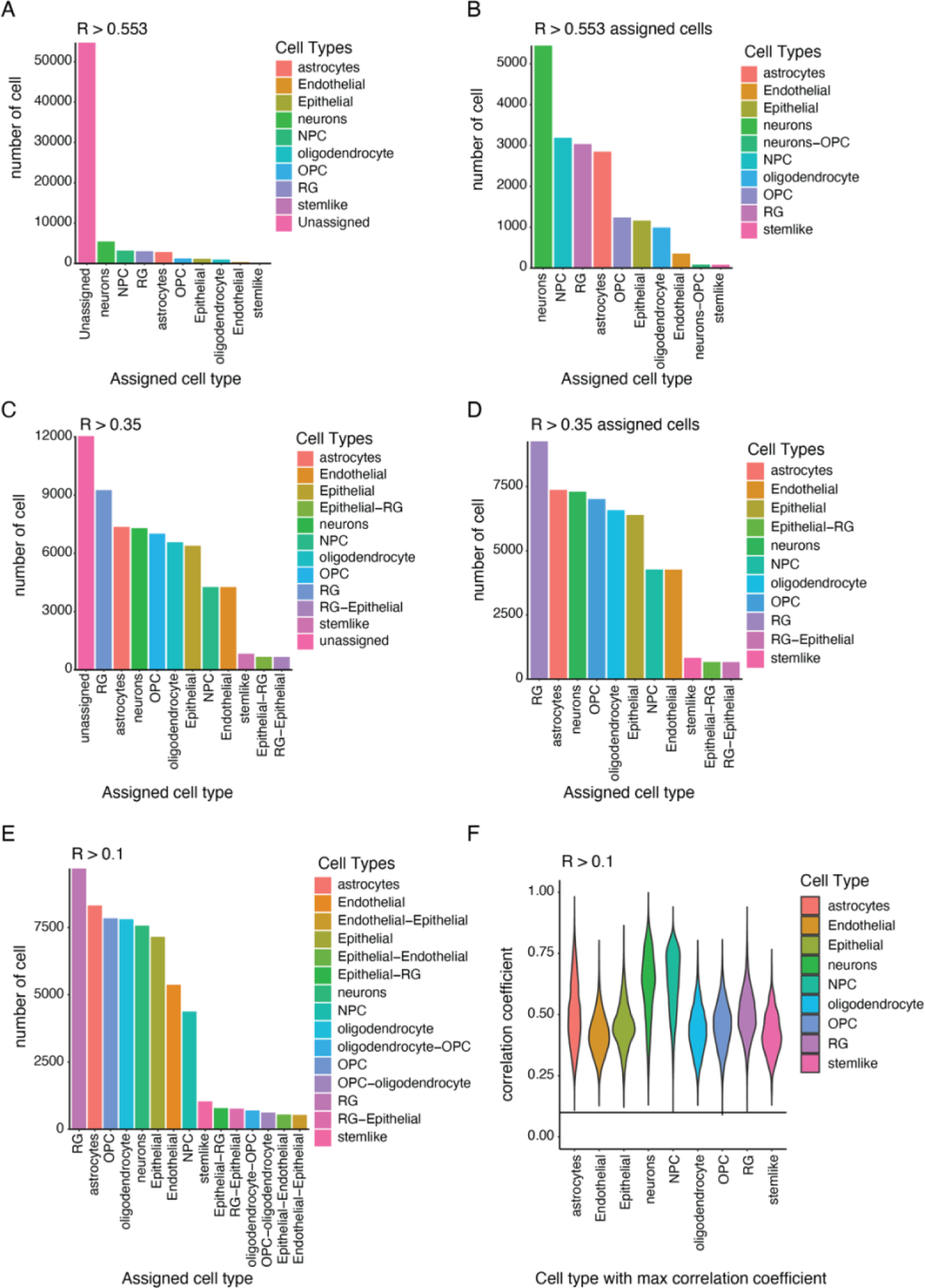
Comparison of different R thresholds for CAM to predict cell types. **A)** Bar chart showing cell counts for assigned cell types and cells left unassigned with the significant R threshold of 0.553. **B)** Bar chart for cell counts excluding the unassigned cells for the R threshold of 0.553. **C)** Bar chart showing cell counts for assigned cell types and cells left unassigned with R threshold of 0.35. **D)** Bar chart for cell counts excluding the unassigned cells for the R threshold of 0.35. **E)** Bar chart showing cell counts for assigned cell types with R threshold of 0.1. All cells pass the threshold with this correlation threshold. **F)** Violin plot showing the max correlation coefficients grouped by the cell type with the max correlation coefficient. The R threshold of 0.1 is indicated by the horizonal line.

**Figure S7:**
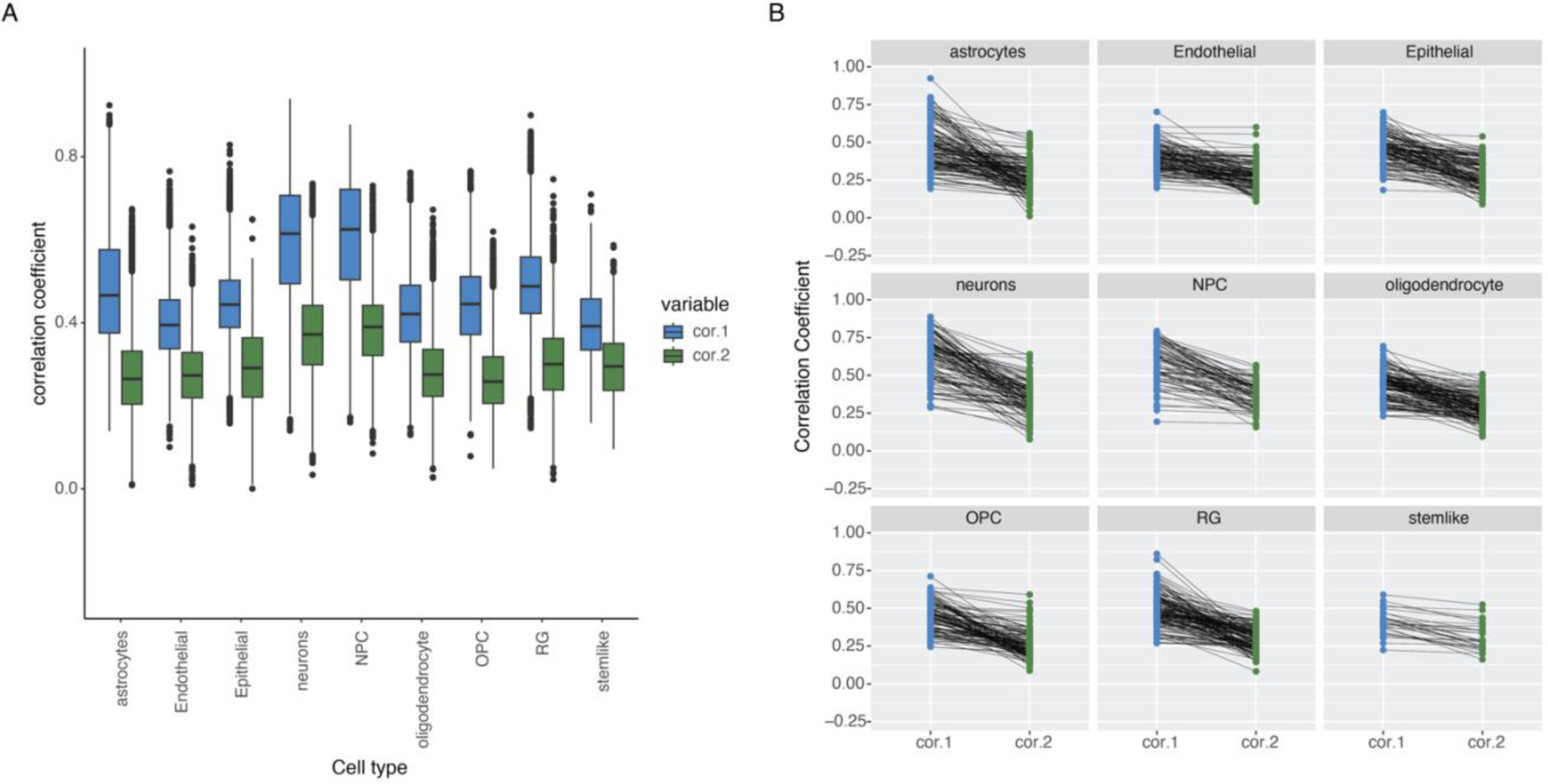
Some cells have high correlation with two cell types. **A)** Box plot showing the max correlation coefficient for each cell type (cor1) and the second max correlation coefficient (cor2) for each cell. The cor2 value is not for a specific cell type, it is the second highest correlation value regardless of the cell type. **B)** Connected point plots from a subsampling of cells in each of the predicted cell types with cor1 matched to cor2. The cor1 values are for the indicated cell type and the cor2 values are corresponding to the second max value for each hMO cell. Horizontal lines indicate that the first and second highest R values are close to equivalent.

**Figure S8:**
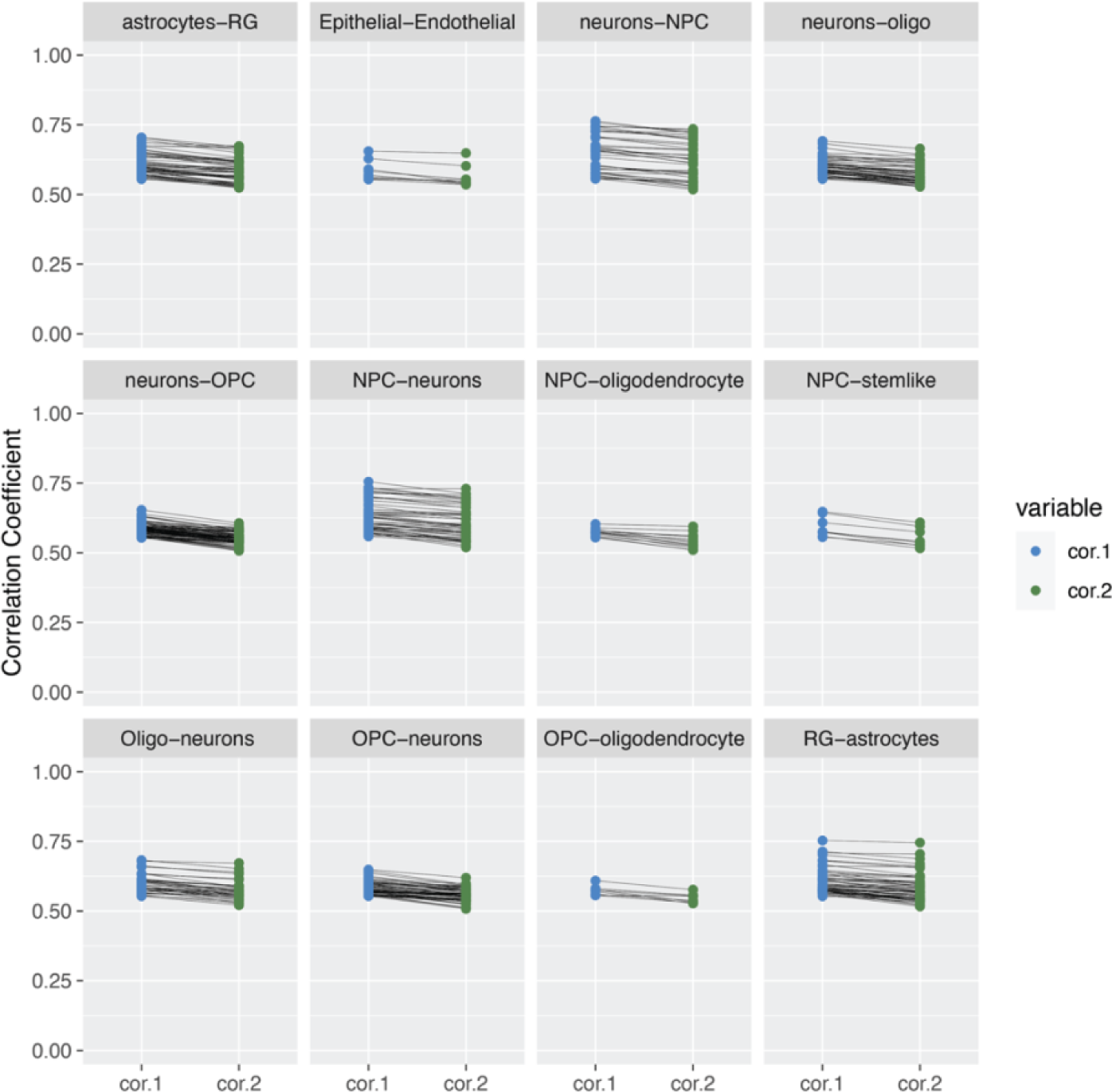
Pairs of cell types with close correlation coefficients could indicate an intermediate cell type. Connected point plots of pairs of predicted cell types with a difference between the first and second highest R values less than 0.05. Only correlation with R value greater than 0.553 were included. Only cell type pairs with more than 8 cells were included. The cell type with the max correlation coefficient is on the left (cor1, blue) and the cell type with the second max correlation coefficient (cor2, green) is on the right. The hMO pairs of correlations coefficients for a given hMO cell are joined by a black line. Cell type names are abbreviated as follows: oligodendrocytes (Oligo), oligodendrocyte precursor cells (OPC), neural precursor cells (NPC), radial glia (RG), neural stem cells (stemlike). Cell types that are a continuum of differentiation, such as neural stem cells and NPCs, or NPCs and neurons have close R values, possibly indicating these cells are starting to express markers of differentiated cell types or retaining some earlier marker expression. The neuron-oligodendrocyte pairs are not a match of a cell type continuum; however, the expression profiles of these cell types overlap.

**Figure S9:**
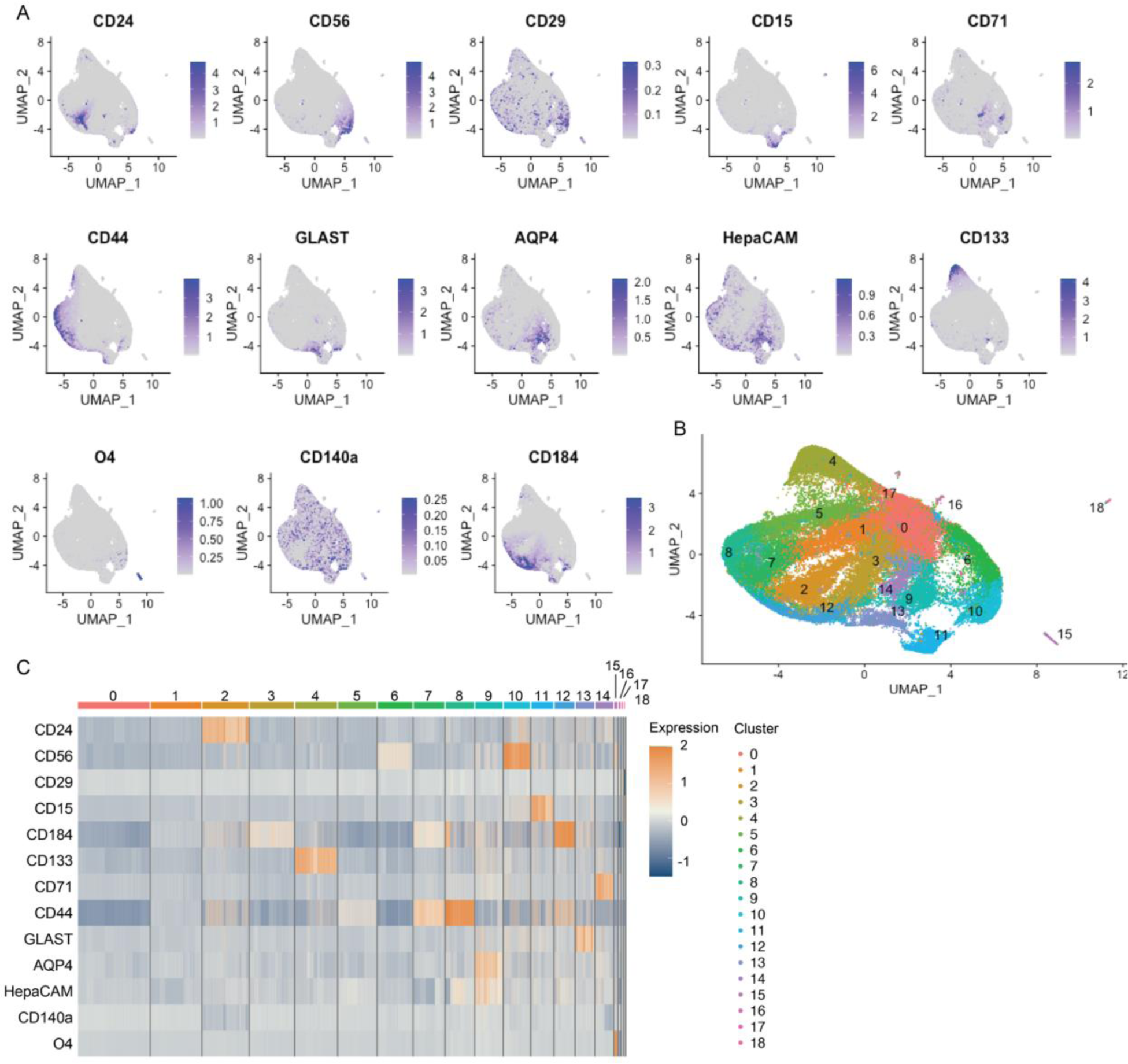
Visualization of antibody expression levels in the subset of hMO cells used for cell type annotation. **A)** Feature plots for each antibody. Normalized expression for the indicated antibodies shown as intensity on UMAPs. The scale for each antibody is indicated. **B)** UMAP pseudo coloured by cluster. Cluster numbers are indicated. **C)** Heatmap of relative expression for each antibody in the panel grouped by cluster.

**Figure S10:**
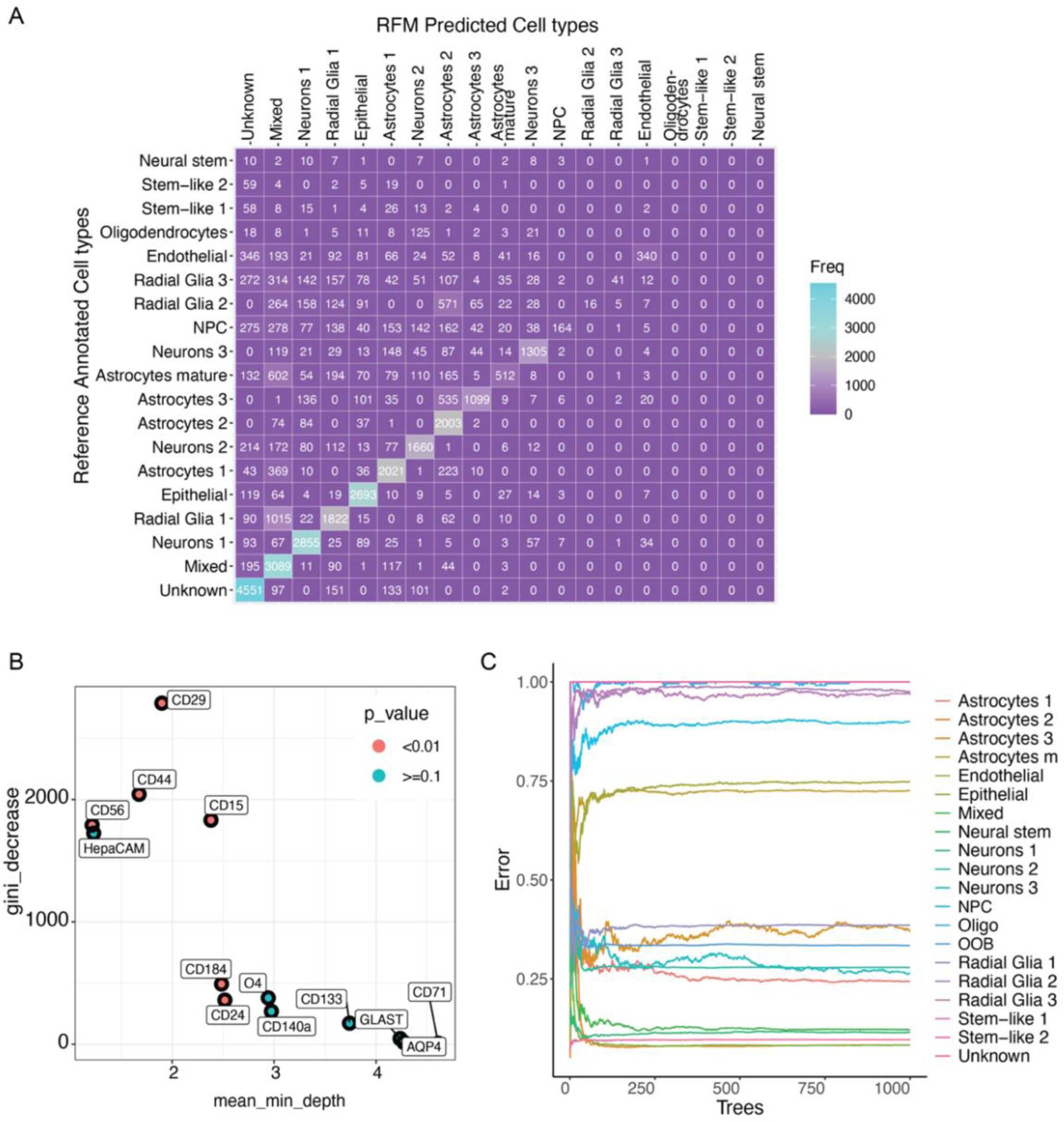
Training random forest model (RFM) with the annotated subset of hMO data. **A)** Confusion matrix showing the number of cells predicted to be in each cell type for a hidden group of cells from the annotated data. The y-axis shows the true label annotation in the subset of cells and the x-axis shows the predicted cell labels. When the x and y axis labels match, the cells are correctly predicted. The number of cells predicted are indicated in the squares. The scale also indicates the number of cells in each true label to predicted label match. **B)** MDS plot showing the contribution of antibodies to the prediction in the RFM. High Gini decrease (y-axis) and lower mean minimum depth (x-axis) indicate a greater importance to classification. **C)** Line graph showing the prediction error (y- axis) for different numbers of trees used in training the RFM (x-axis). Coloured lines correspond to each cell type. OOB is the overall error rate. A low error rate indicates a better prediction. Most cell types are predicted accurately, but radial glia2, radial glia3, and oligodendrocytes are not predicted well in the RFM.

**Figure S11:**
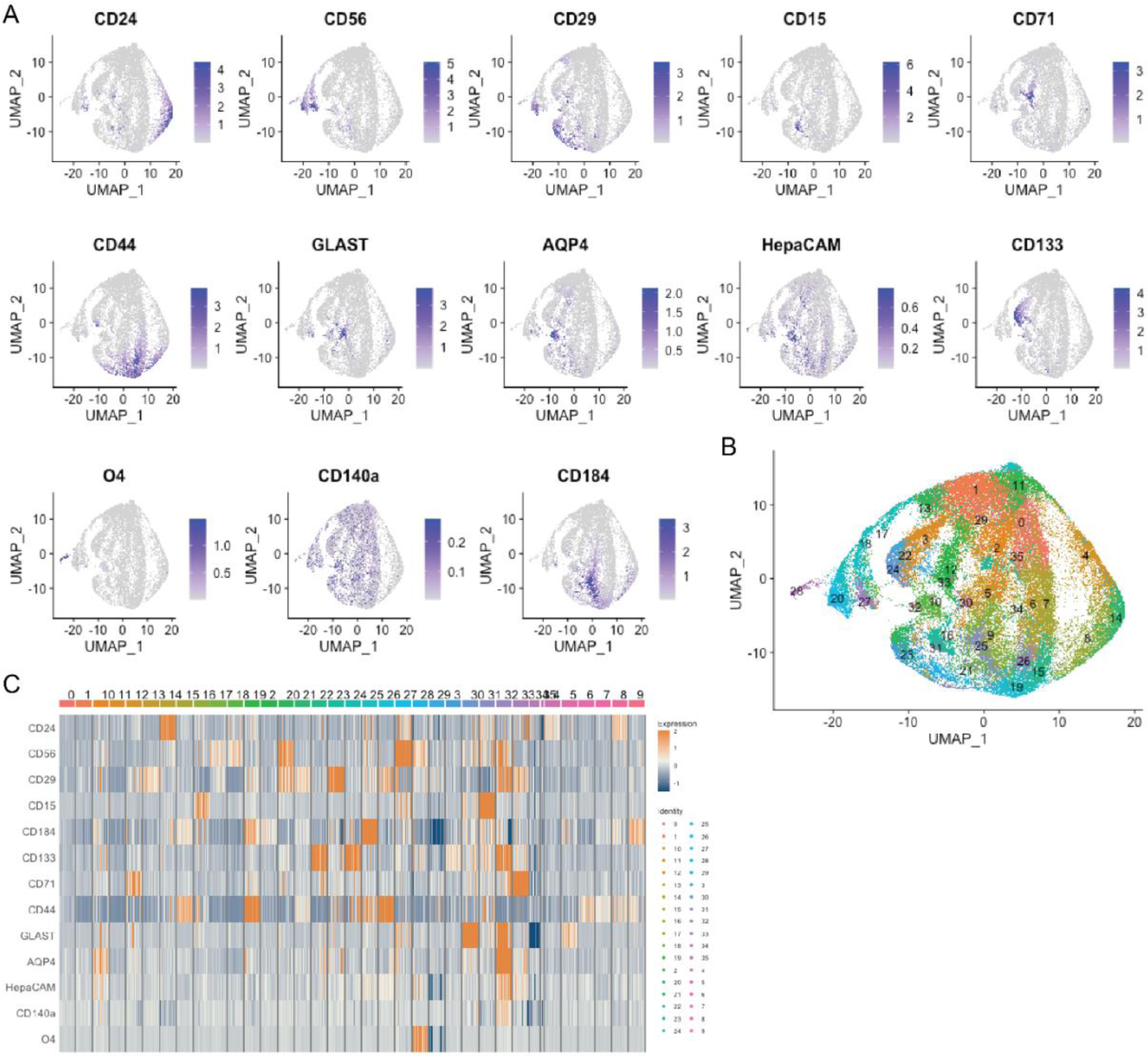
Visualization of protein expression levels in the full nine sample hMO dataset for cell type annotation. **A)** Feature plots for each antibody. **B)** UMAP visualization of Seurat clusters. **C)** Heatmap protein expression per cluster.

**Figure S12:**
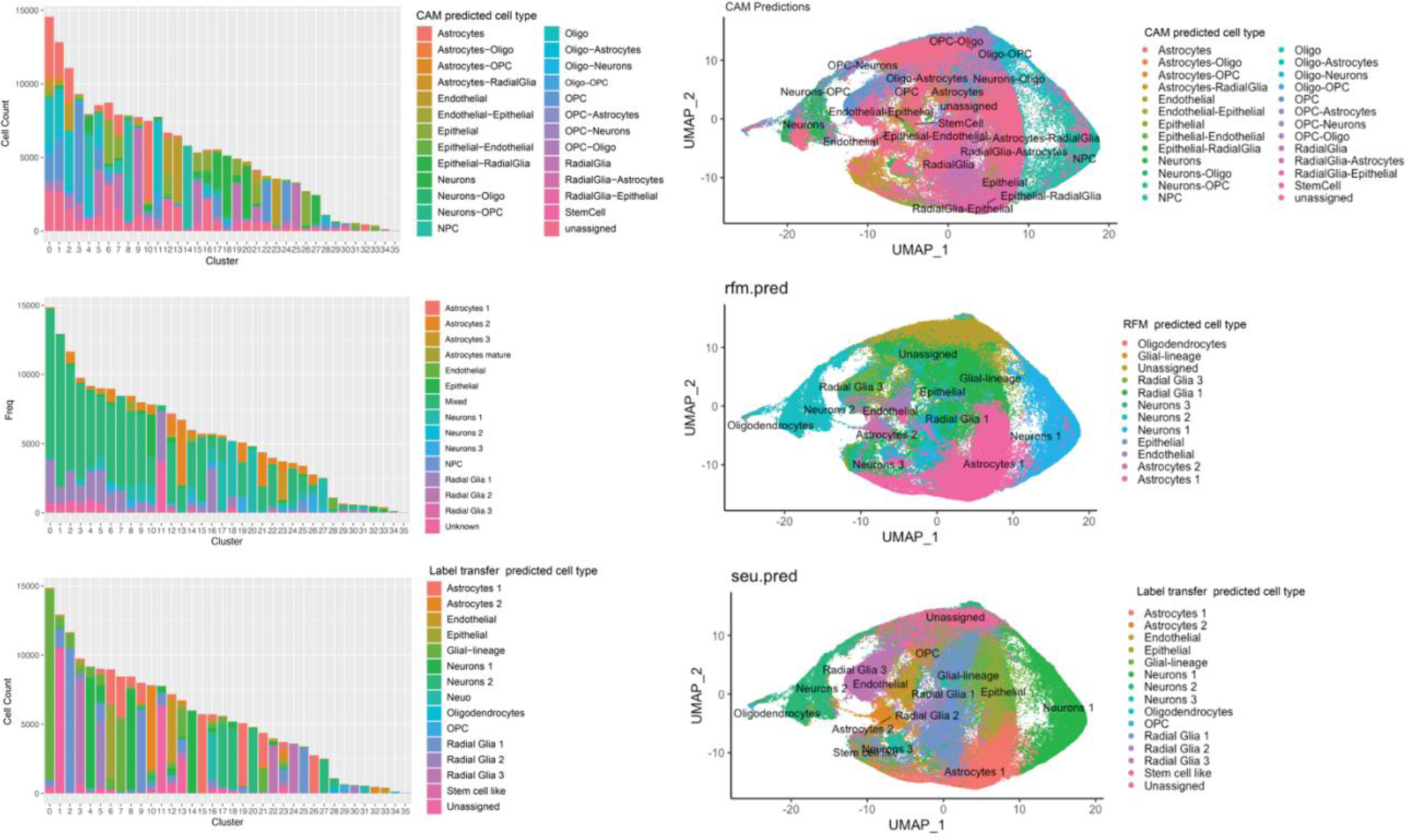
Cell type annotation predictions from CAM, RFM and Seurat label transfer. Left, bar charts with the counts of cells predicated as each cell type in each cluster. Right, UMAPs colours by predicted cell types. Top; CAM predictions with an R threshold for assignment of 0.35 and a double cell type threshold of max R-second max1 of less than 0.01. The results were then filtered to included only predicted cell types with over 200 cells. Middle, RFM predictions from the model trained on the subset of 9000 cells from each of 9 hMOs. Bottom, Seurat label transfer method using the Seurat object from the subset of cells as the reference data.

**Figure S13:**
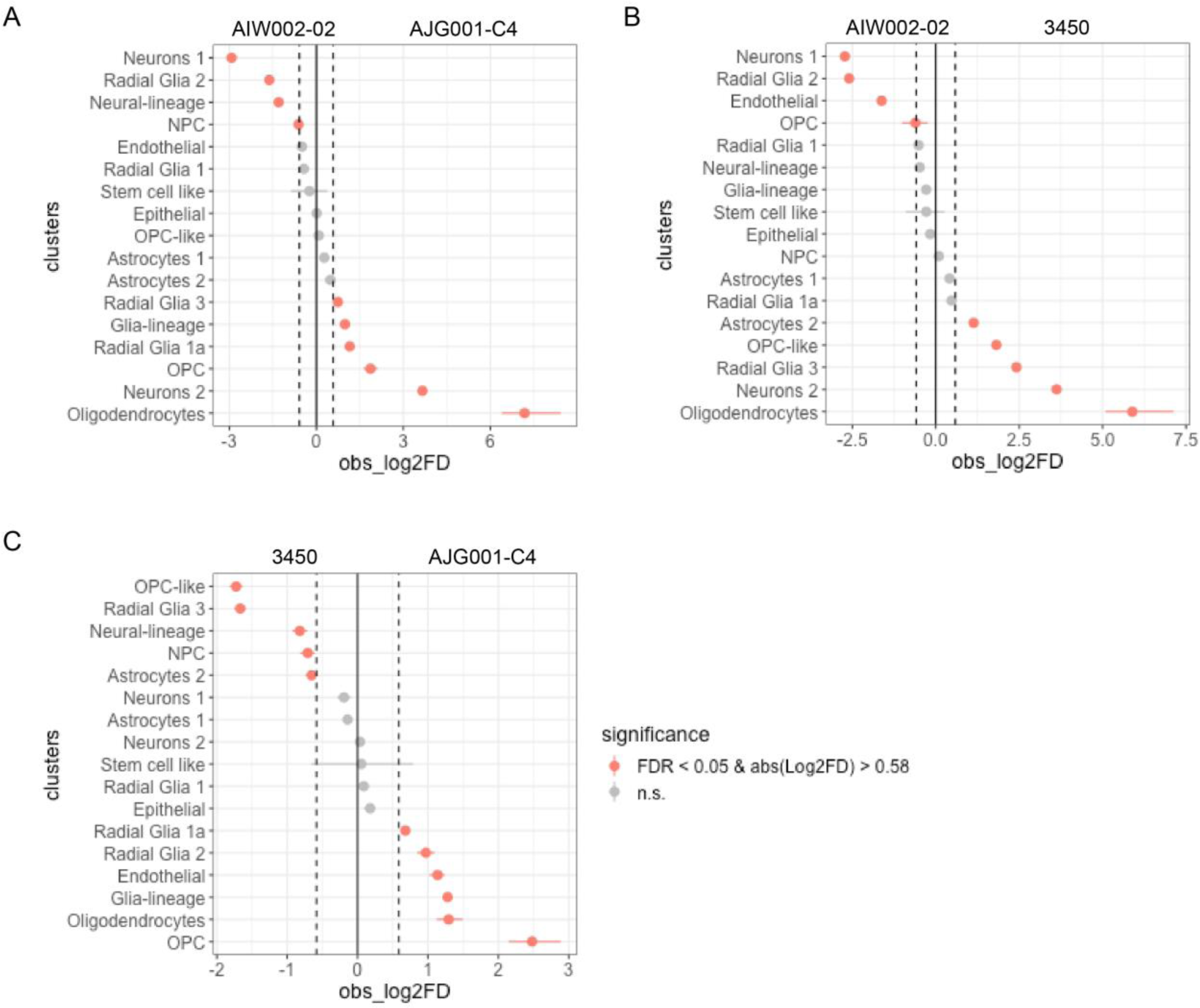
Proportionality test of cell types between each iPSC line. Point range plots showing significant differences in proportions of cell types between iPSC lines. The line contrasts are indicated above the plots. Pink dots indicate a significant difference in the proportion of the indicated cell type between the two iPSC lines. Each iPSC line has three samples. **A)** AIW002-02 compared to AJG001- C4. **B)** AIW002-02 compared to 3450. **C)** AJG001-C4 compared to 3450.

**Figure S14:**
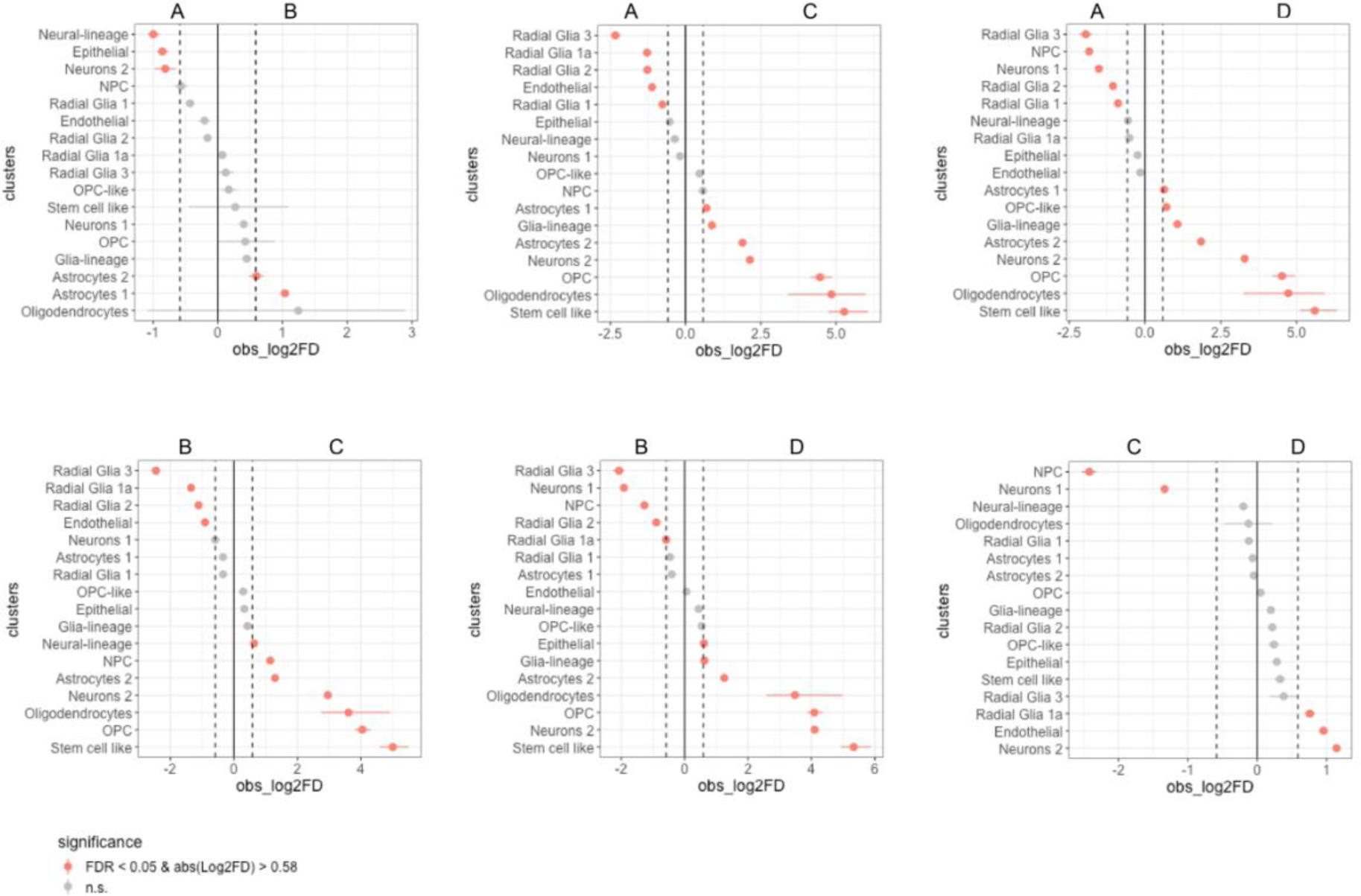
Proportionality test of cell types between AIW002-02 batches. Point range plots showing significant differences in proportions of cell types between hMO batches. The line contrasts are indicated above the plots. Pink dots indicate a significant difference in the proportion of the indicated cell type between the two batches.

**Figure S15:**
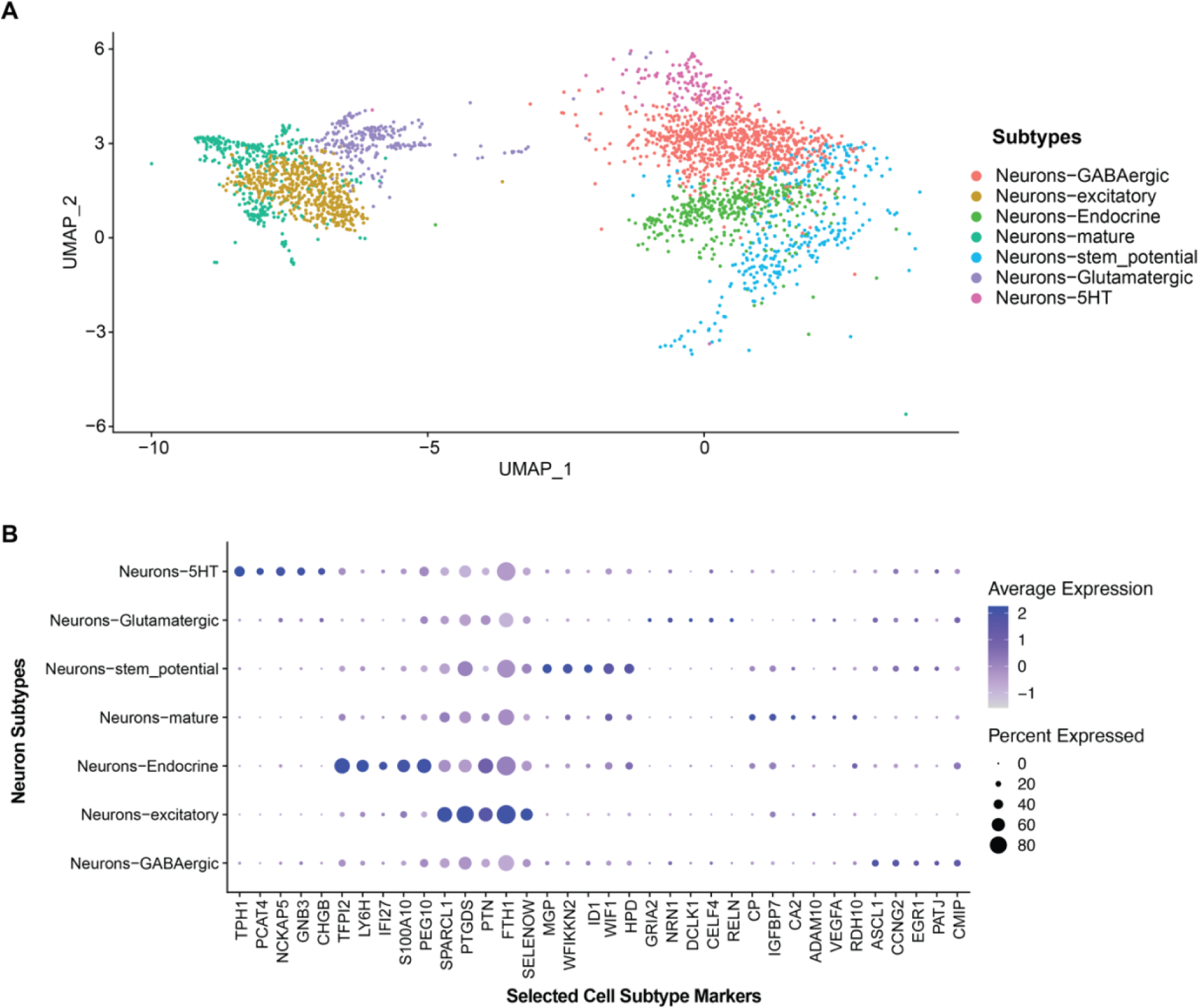
Neuronal subtypes and selected markers. **A)** Neurons were subset from the total population and plotted on a UMAP. Subtypes based are differentially expressed genes are indicated in the legend. **B)** Dot plot of 5 selected differentially expressed genes. The proportion of cells in each group expressing a marker is indicated by dot size and the expression level is shown by intensity.

**Figure S16:**
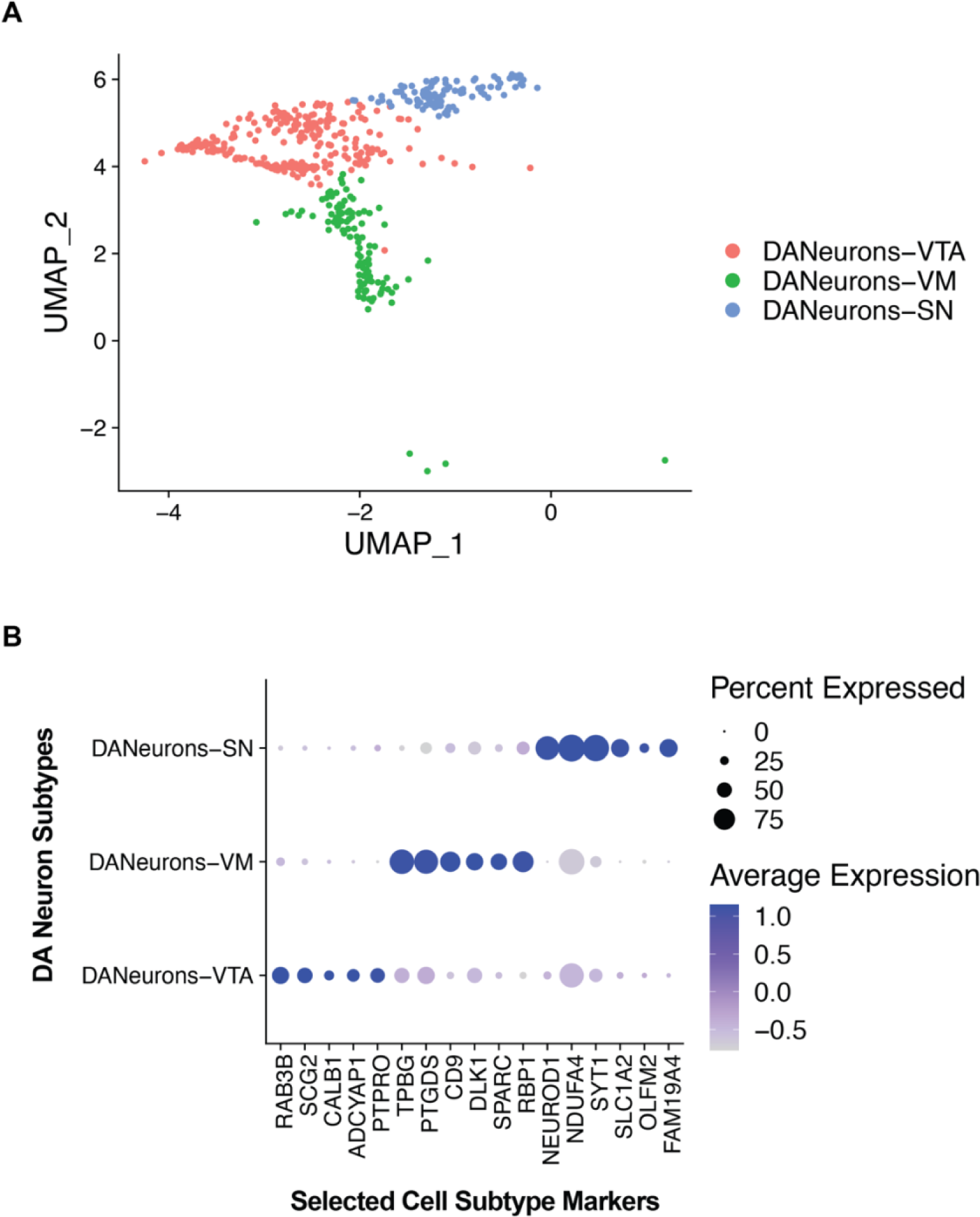
DA neuronal subtypes and selected markers. **A)** DA neurons were subset from the total population and plotted on a UMAP. Subtypes were annotated using the markers identified by differential gene expression and analysis of expression of known markers and subtypes DANeurons-VTA (ventral tegmental area), DANeurons-VM (ventral midbrain) and DANeurons-SN (substantia nigra). See Table S13 and S14. **B)** Dot plot of 5 selected differentially expressed genes. The proportion of cells in each group expressing a marker is indicated by dot size and the expression level is shown by intensity.

**Figure S17:**
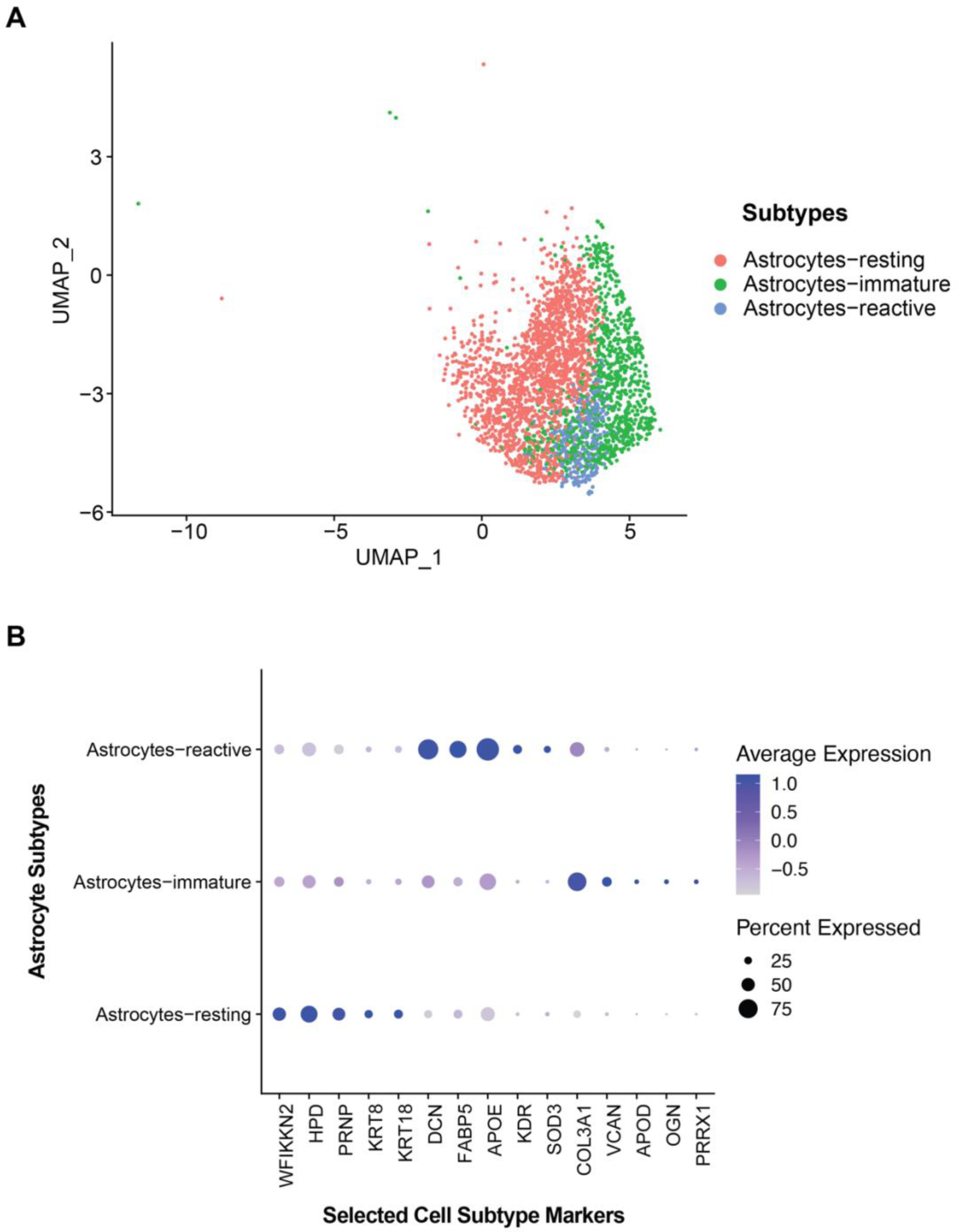
Astrocyte subtypes and selected markers. **A)** Astrocytes were subset from the total population and plotted on a UMAP. Subtypes based are differentially expressed genes are indicated in the legend. **B)** Dot plot of 5 selected differentially expressed genes. The proportion of cells in each group expressing a marker is indicated by dot size and the expression level is shown by intensity.

**Figure S18:**
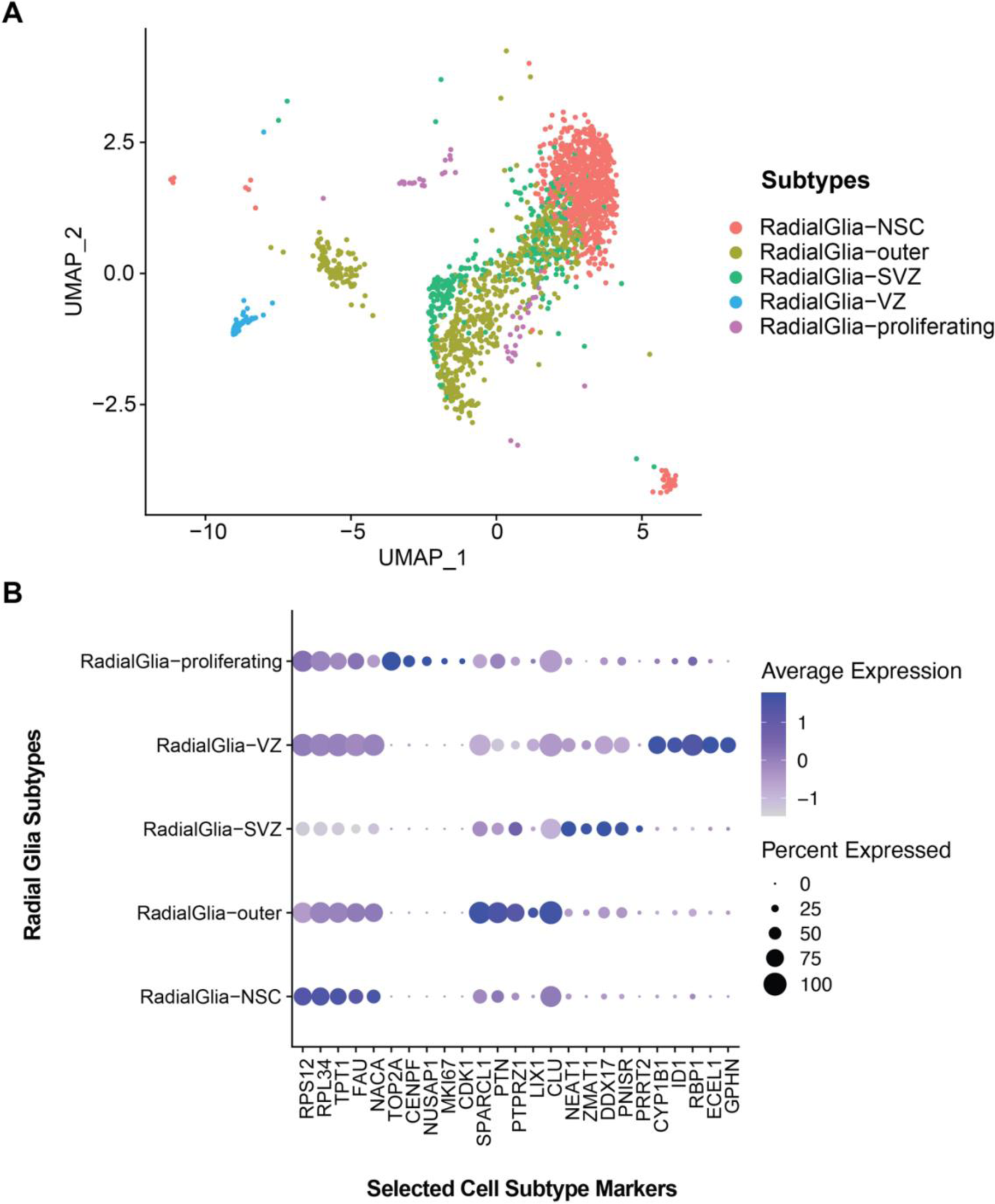
Radial glia subtypes and selected markers. **A)** Radial glia cells were subset from the total population and plotted on a UMAP. Subtypes based on differentially expressed genes are indicated in the legend**. B)** Dot plot of 5 selected differentially expressed genes. The proportion of cells in each group expressing a marker is indicated by dot size and the expression level is shown by intensity.

**Figure S19:**
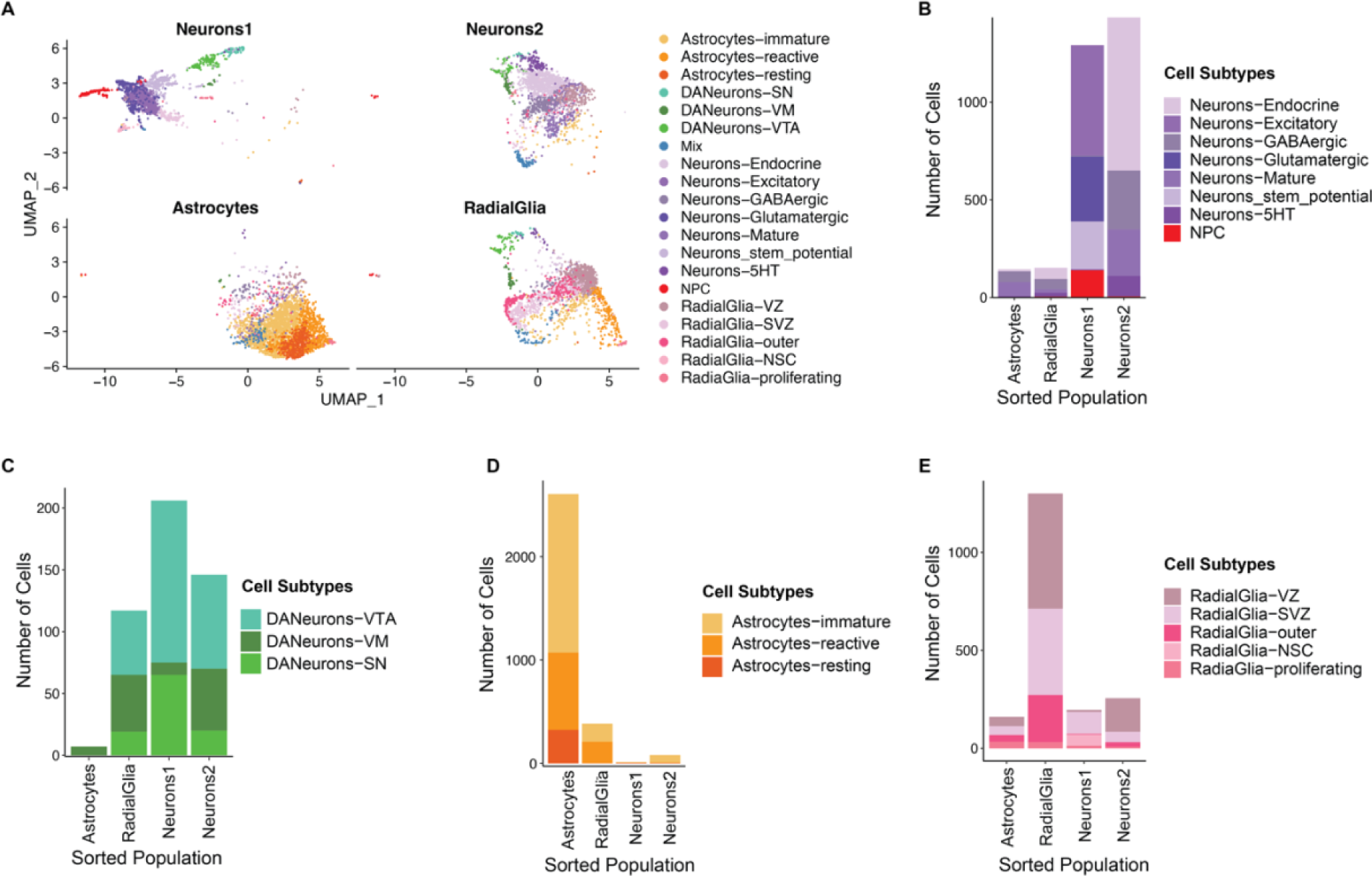
Visualization of the number of cellular subtypes identified by scRNAseq transcriptomes in each FC sorted population. **A)** UMAP split by sorted population and coloured by cell subtypes. **B)** Stacked bar chart showing the number of each non-DA neuron cell subtype in each sorted population. **C)** Stacked bar chart showing the number of each DA neuron cell subtype in each sorted population. **D)** Stacked bar chart showing the number of each astrocyte cell subtype in each sorted population. **E)** Stacked bar chart showing the number of each radial glia cell subtype in each sorted population.

**Figure S20:**
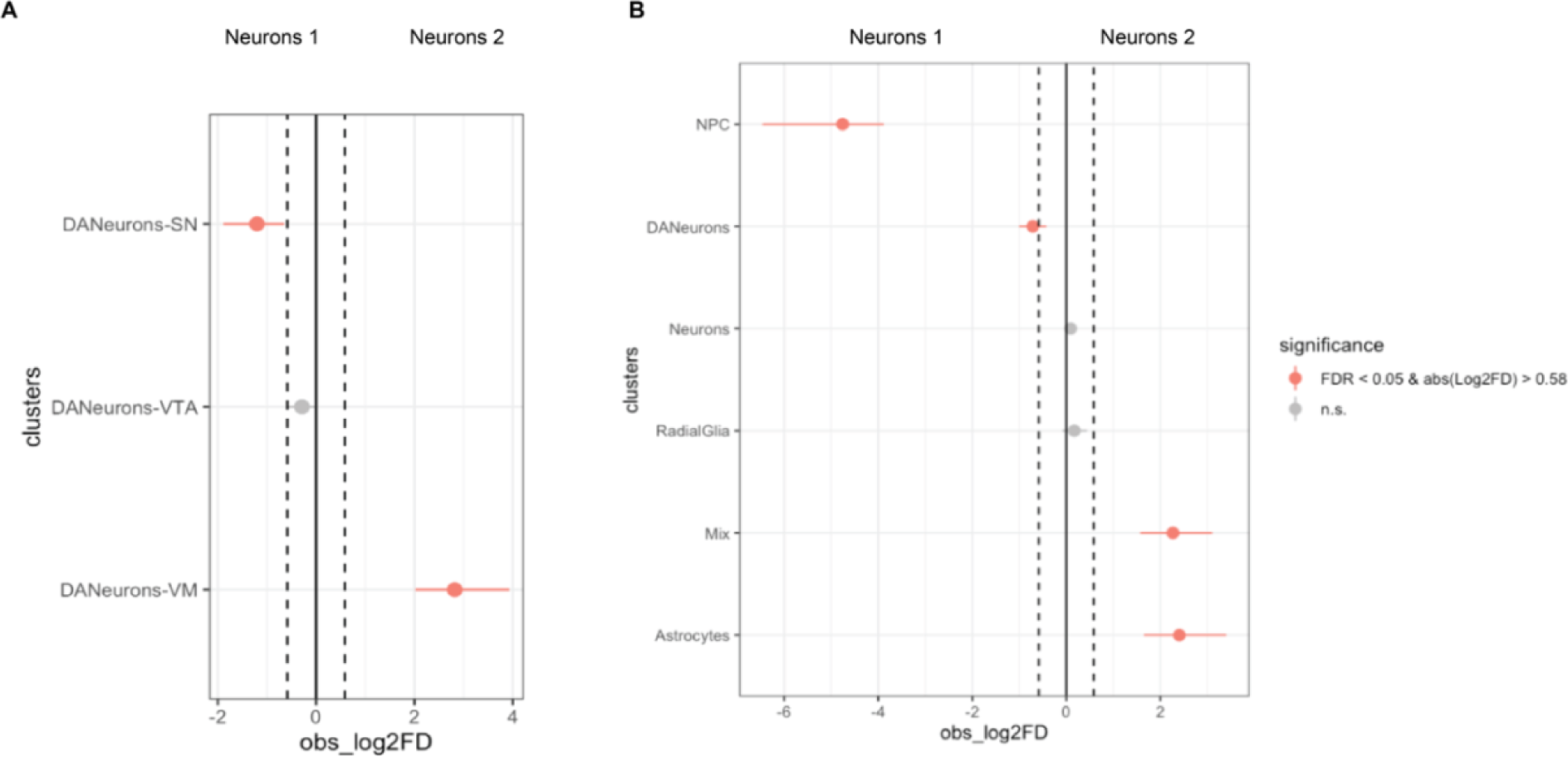
Proportion of cell types and DA neuron subtypes in FACS populations neurons 1 compared to neurons 2. **A)** Point range plot showing the differences in proportions of DA neuron subtype cells neurons1 and neurons 2. **B)** Point range plot showing the differences in proportions of cell types in neurons1 compared to neurons2. Negative log2FD values indicate a greater proportion of cells in neurons1 and positive log2FD values indicate a greater proportion of cells in neurons2. Pink dots indicate a significant difference.

**Figure S21:**
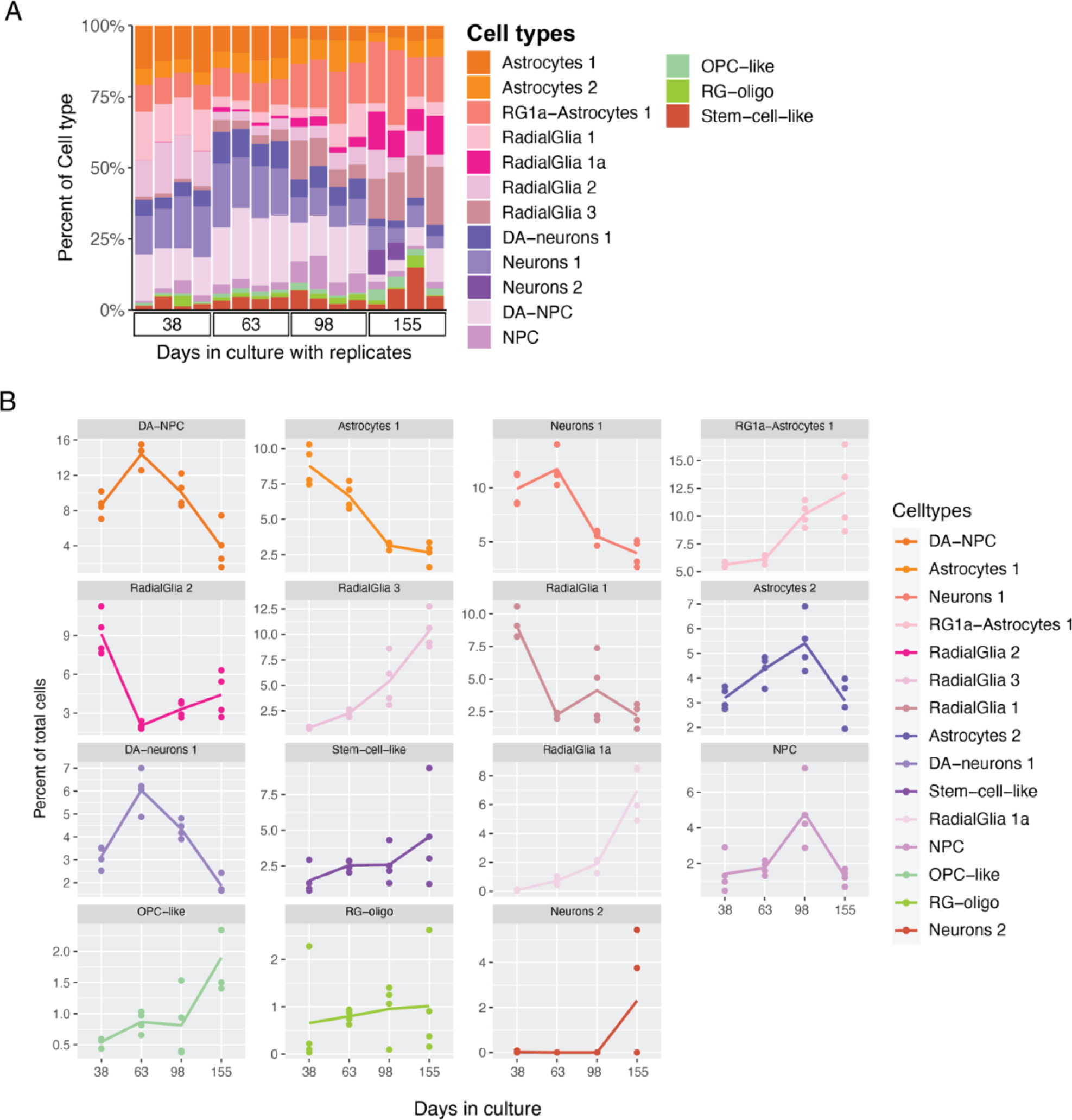
Proportions of cell types in AIW002 over time. **A)** Bar chart showing proportions of cell types in each sample. Cell types are coloured and shown in the legend. Sample are replicates grouped by the indicated type points. **B)** Line plots showing the proportion of each cell type over time. Dots are shown for each replicate, time points are shown on the x-axis. Each cell type is shown separately. Cell types are indicated in the legend and facet labels above each plot.

**Methods Figure 1:**
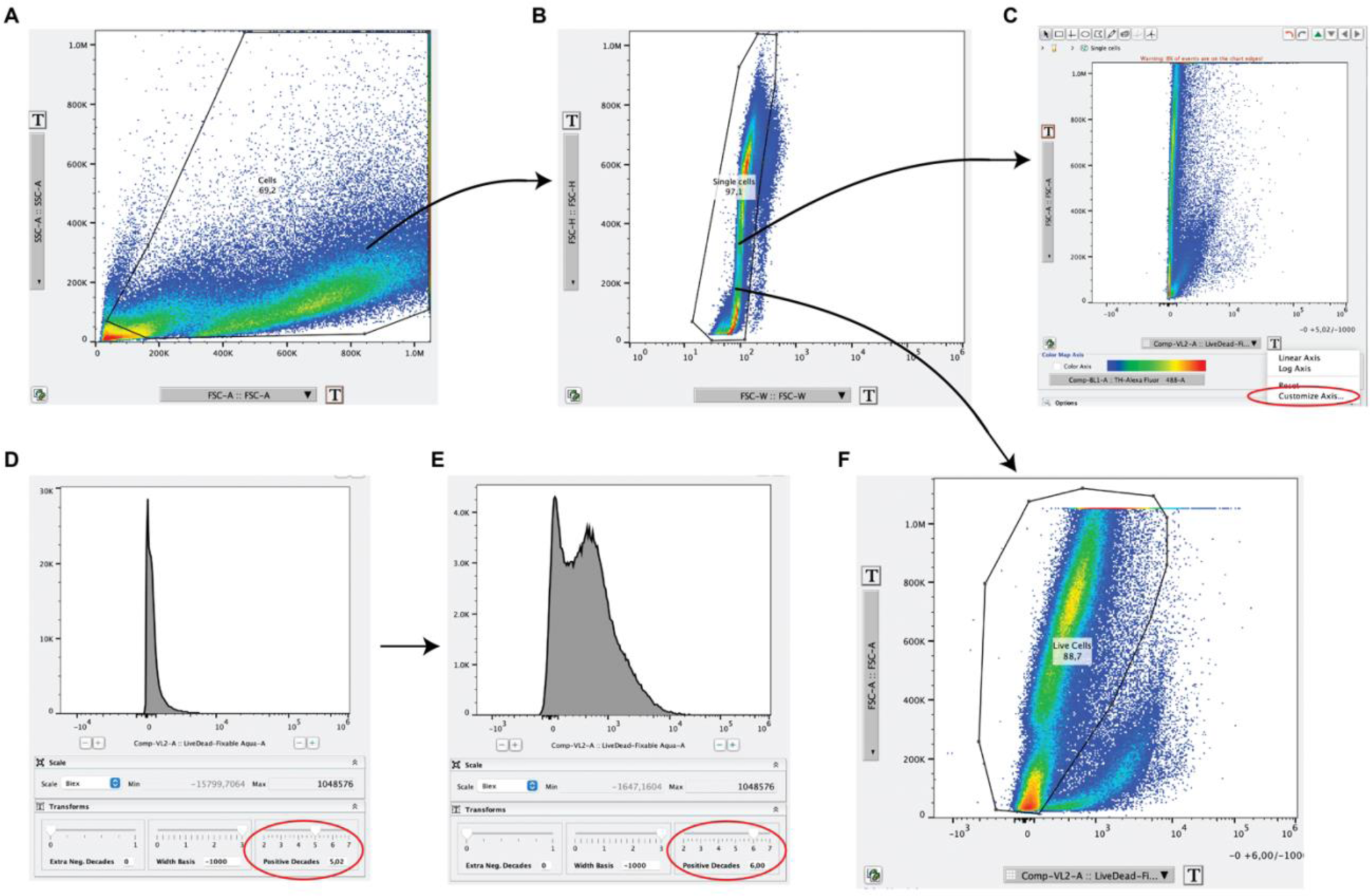
Live cell gating strategy. **A)** Using SCC-A by FSC-A axis the cells are selected, and debris is excluded. Double clicking the gated cells will give the selected population. **B)** Change the axis to FSC-H by FSC-W and adjust the x-axis (FSC-W) to log scale. (By clicking the ‘T’). Draw a gate on the cells on the left to select single cells and exclude doublets. **C)** Change the axis to FSC-A by the LiveDead stain. The dead cells take-up the dye. To easily distinguish the live and dead cell populations the x-axis scale needs to be adjusted. Click the ‘T’ and select ‘Customize Axis’ **D)** The custom axis window. The scale ‘Positive Decades’ (circled in red) needs to be adjusted. **E)** Adjust the scale until you can see two peaks in the histogram. Click apply. **F)** With the adjusted scale the live cell population is now gated from the dead cells by drawing a gate. These live single cells are the cells selected for further analysis.

**Table S1:** Main effect of iPSC line from a 2-way ANOVA and Tukey’s posthoc test of the significant differences. Significant differences with a p-value < 0.05 are highlighted in bold. P values are shown for the 2-way ANOVA main effects and the adjusted p Value from the Tukey’s HSD post hoc tests.

| Cell type | Variable | Contrast | Test | P value |
| --- | --- | --- | --- | --- |
| Neurons 1 | IPSC | Main effect | ANOVA 2-way | **0.00046** |
| Neurons 2 | IPSC | Main effect | ANOVA 2-way | **0.00174** |
| NPC | IPSC | Main effect | ANOVA 2-way | **0.04065** |
| Oligodendrocytes | IPSC | Main effect | ANOVA 2-way | **0.00127** |
| OPC-like | IPSC | Main effect | ANOVA 2-way | **0.00177** |
| Neurons 1 | IPSC | AJG001-AIW002 | Tukey | **0.00409** |
| Neurons 1 | IPSC | 3450-AIW002 | Tukey | 0.87746 |
| Neurons 1 | IPSC | 3450-AJG001 | Tukey | **0.00085** |
| NPC | IPSC | AJG001-AIW002 | Tukey | 0.96515 |
| NPC | IPSC | 3450-AIW002 | Tukey | **0.05442** |
| NPC | IPSC | 3450-AJG001 | Tukey | 0.09643 |
| Neurons 2 | IPSC | AJG001-AIW002 | Tukey | 0.76817 |
| Neurons 2 | IPSC | 3450-AIW002 | Tukey | **0.00215** |
| Neurons 2 | IPSC | 3450-AJG001 | Tukey | **0.01683** |
| Oligodendrocytes | IPSC | AJG001-AIW002 | Tukey | **0.00081** |
| Oligodendrocytes | IPSC | 3450-AIW002 | Tukey | 0.07856 |
| Oligodendrocytes | IPSC | 3450-AJG001 | Tukey | 0.10891 |
| OPC-like | IPSC | AJG001-AIW002 | Tukey | **0.01652** |
| OPC-like | IPSC | 3450-AIW002 | Tukey | **0.00222** |
| OPC-like | IPSC | 3450-AJG001 | Tukey | 0.77824 |

**Table S2:**
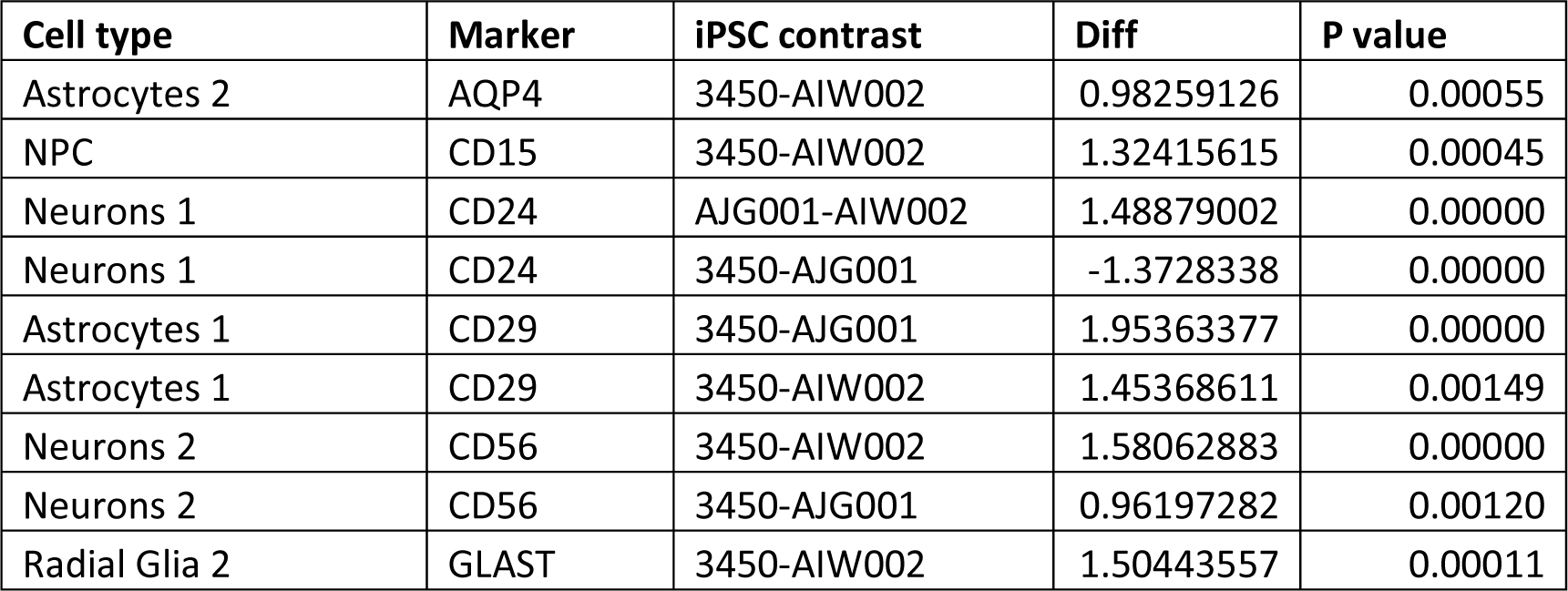
Tukey’s HSD test of the interaction effect between iPSC line and protein expression. Showing all significant differences between iPSC pairs for a given protein marker. The significance threshold was considered p value < 0.05. Mean values across cells were taken for each sample in each cell type (n=3). Diff, indicates the difference between the mean values of the 3 replicates.

**Table S3:** *Hypergate* prediction of cell types. Astrocytes 1 and 2 are combined and radial glia 1 and 2 are combined. Neurons1, neurons2, NPCs, and oligodendrocytes are left as separate populations. The rest of the cell types were combined into one group labelled ‘other’.

| Cell type | Accuracy | Gating strategy |
| --- | --- | --- |
| Astrocytes | 95.30% | "CD44 >= 69134.93, CD184 >= 4790.36, CD24 <= 14941.93, CD15 <= 21652.39, CD133 <= 16111.91, CD71 <= 24201.84, CD56 <= 31661.17, CD184 <= 41979.12, GLAST >= -1829.09, CD29 >= -221.8, GLAST <= 19910.19, O4 <= 28771.54, CD15 >= -5153.37, O4 >= -13039.53, CD133 >= -5203.24" |
| Endothelial | 97.78% | "CD71 >= 10690.89, CD133 <= 14612.87, CD56 <= 23612.89, CD44 <= 154039.7, CD15 <= 33664.2, CD184 >= -8415.42, O4 <= 6648.85, HepaCAM >= -2266.38, CD184 <= 26695.98, GLAST >= -3953.12, AQP4 <= 179930, GLAST <= 41721.09, CD24 <= 204827.5, CD29 <= 93582.52, CD133 >= -6085.38" |
| Epithelial | 96.14% | "CD44 >= 57643.44, CD184 <= 3973.42, CD24 <= 13819.15, CD133 <= 10956.84, CD15 <= 18368.35, CD44 <= 217613.6, CD184 >= -7614.27, CD29 <= 41852.39, CD71 <= 16803.39, CD56 <= 27459.87, HepaCAM >= -2503.27, GLAST >= -21787.42, O4 <= 13506.06, GLAST <= 7895.25, O4 >= -15995.77, CD71 >= -7398.14, AQP4 >= -27593.11, AQP4 <= 58460.66, CD140a <= 2380.11" |
| Glial lineage | 98.19% | "CD44 >= 18981.62, CD44 <= 67947.27, CD24 <= 8532.04, CD133 <= 6503.12, CD184 >= 671.64, CD184 <= 6027.05, CD56 <= 10803.37, CD15 <= 9272.85, CD71 <= 9899.44, CD29 <= 18512.49, GLAST <= 2972.65, GLAST >= -4303.57, O4 >= -8600.3, AQP4 <= 25934.81, O4 <= 3831.41, AQP4 >= -30283.57, HepaCAM <= 2853.07, CD15 >= -8752.85" |
| Neural lineage | 99.12% | "CD29 <= 5164.46, CD44 <= 16345.29, CD56 <= 8228.51, CD15 <= 10097.88, CD24 <= 14430.59, CD133 <= 4690.34, AQP4 <= 10768.68, CD71 <= 9058.63, CD184 >= -4118.05, CD184 <= 4616.04, GLAST <= 3205.48, GLAST >= -5846.58, HepaCAM >= -2172.31, AQP4 >= -32414.35, O4 <= 7012.81, O4 >= -10615.22, CD15 >= -9709.93" |
| Neurons 1 | 97.84% | "CD24 >= 11411.88, CD71 <= 13046.05, CD15 <= 26938.4, CD133 <= 9407.79, CD56 <= 24563.79, HepaCAM >= -2301.21, GLAST <= 7464.28, GLAST >= -10157.93, CD29 >= 388.64, CD29 <= 32358.1, CD184 >= -8356.64, CD184 <= 34967.37, AQP4 <= 54412.99, CD44 <= 330276.01, CD44 >= 370.32, CD71 >= -6552.91, O4 <= 5850.52, AQP4 >= -36288.88, CD140a <= 1853.66" |
| Neurons 2 | 97.35% | "CD56 >= 11306.47, O4 <= 6512.56, CD184 <= 5993, CD15 <= 35454.67, CD133 <= 14378.52, CD44 <= 69578.63, CD184 >= -4544.7, HepaCAM <= 2516.97, AQP4 <= 56704.2, O4 >= -3686.94, CD71 <= 37546.04, GLAST >= -8086.99, CD24 <= 89382.14, CD133 >= -6581.01, AQP4 >= -35872.41, CD29 >= -1750.44, CD29 <= 127463.18, CD44 >= -1745.39, GLAST <= 46130.06" |
| NPC | 98.19% | "CD15 >= 16883.48, CD56 <= 68152.13, CD29 <= 32492.33, O4 <= 6603.71, CD133 <= 34264.57, CD71 <= 33362.52, CD44 <= 579529, GLAST >= -8585.92, CD184 >= -8395.97, AQP4 <= 143544.3, CD24 <= 126973.72, CD184 <= 64418.66, HepaCAM <= 8196.44, CD24 >= -10750.26, CD56 >= -863.58, GLAST <= 100720.56, CD140a <= 4207.56, O4 >= -20194.79, CD133 >= -4657.86" |
| Oligodendrocytes | 99.85% | "O4 >= 6954.58, CD71 <= 259264.61, AQP4 <= 251950.1, CD24 <= 79343.15, GLAST >= -12629.13" |
| OPC | 99.14% | "CD184 <= -3881.47, O4 <= 5819.9, HepaCAM <= 2576.93, CD15 <= 20078.87, CD133 <= 24914.78, GLAST <= 25128.87, CD44 <= 169438.5, CD71 <= 47633.15, CD56 <= 70209.7" |
| OPC-like | 98.69% | "CD133 >= 17973.84, HepaCAM <= 7386.68, CD15 <= 93841.76, AQP4 <= 93903.5, O4 <= 19514.81, CD71 <= 171467.93, CD24 <= 186318.34, CD29 <= 75236.89, CD184 >= -18571.9, CD56 <= 102289.4, CD184 <= 66643.74, GLAST <= 54848.83, HepaCAM >= -8102.46" |
| Radial Glia | 91.93% | "CD184 >= 4670.84, CD24 <= 13151.85, CD15 <= 16116.35, CD44 <= 108487.72, AQP4 <= 27956.54, CD133 <= 19864.36, CD71 <= 10261.43, CD29 <= 39977.27, O4 <= 2775.15, GLAST >= -2949.46, CD56 <= 43016.34, O4 >= -6620.26, HepaCAM >= -1840.22, CD56 >= -1630.13, CD71 >= -8436.27, CD44 >= -164.1, AQP4 >= -27462.2, HepaCAM <= 5943.6" |
| Stem-like | 99.95% | "GLAST <= -5784.06, CD133 <= 17757.4, CD184 >= -2183.33, O4 <= 43826.44, CD44 <= 132336.8" |

**Table S4:** Correlation coefficient between protein intensity levels measured by FC and RNA expression measured by scRNAseq. The four FACS sorted populations are indicated. The 13 proteins targeted in the FC antibody panel and the corresponding genes are used as the input.

|  |  | scRNAseq |  |  |  |
| --- | --- | --- | --- | --- | --- |
|  |  | Astrocytes | RadialGlia | Neurons1 | Neurons2 |
| FC | Astrocytes | 0.55533284 | 0.15228784 | -0.4517431 | -0.1806433 |
|  | RadialGlia | 0.13058154 | 0.44510732 | -0.3314321 | 0.05559489 |
|  | Neurons1 | -0.4098105 | -0.3016957 | 0.45349104 | 0.05175022 |
|  | Neurons2 | -0.4858683 | -0.1624758 | 0.37703856 | 0.23752196 |

**Table S5:**
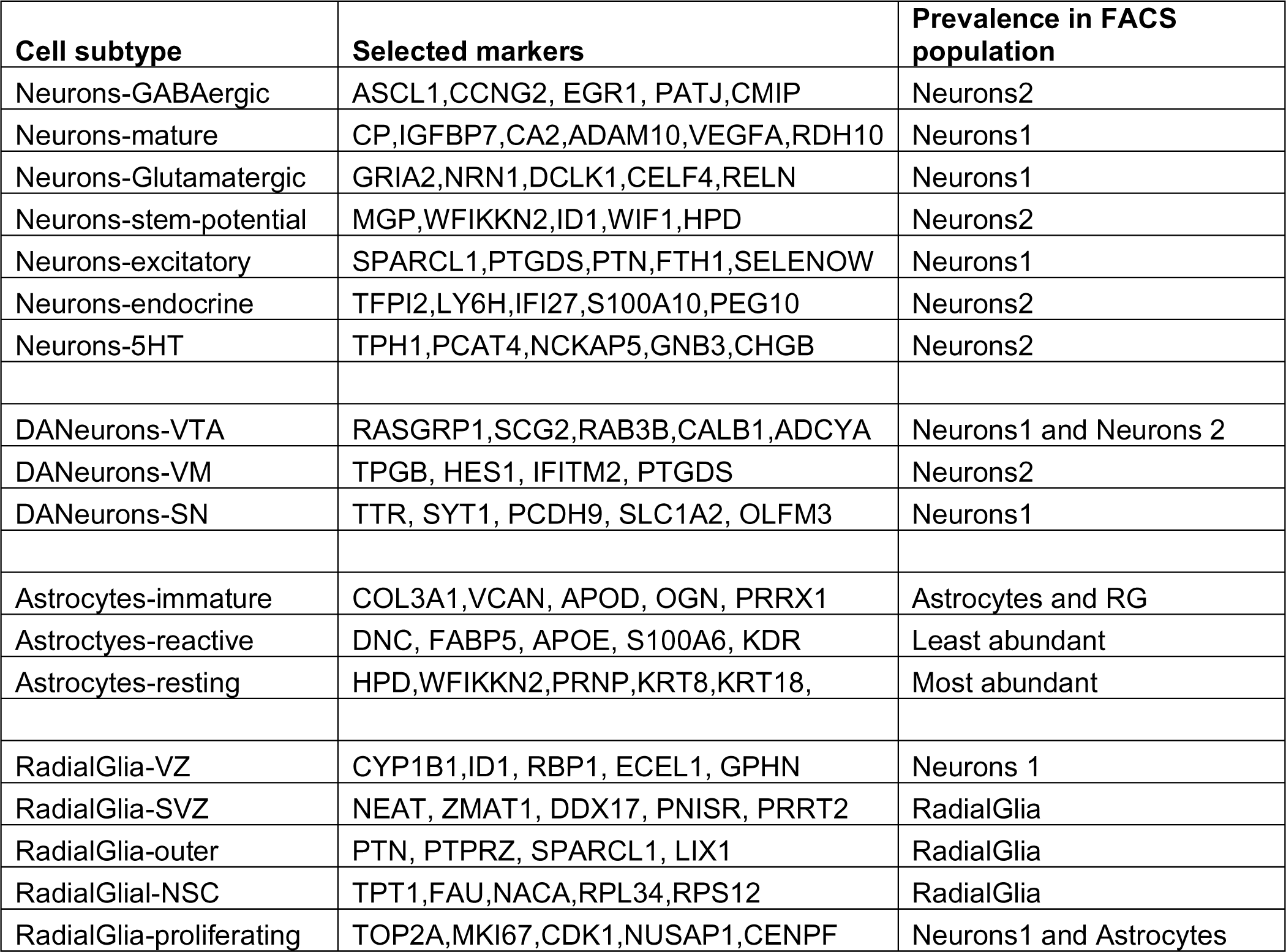
Cell type name with selected cell type markers and which FACS sorted population contains each cell subtype.

**Table S6:** References of DA neuron subtypes for selected cluster markers.

| Subgroup Cluster | Marker gene | Region or subtype indicated | Reference |
| --- | --- | --- | --- |
| DA1<br>DANeurons-VTA | RAB3B | VM DA maturation | Monzon-Sandoval 2020<sup>57</sup> |
|  | SCG2 | VTA | Wen 2021<sup>58</sup>, Greene 2015<sup>59</sup> |
|  | CALB1 | VTA | Chung 2005<sup>60</sup>, Greene 2005<sup>59</sup> |
|  | RASGRP1 | VM | Eshraghi 2020<sup>61</sup> |
|  | STMN2 | VM DA development | Yin 2009<sup>62</sup>, Fernandes 2020<sup>63</sup> |
|  | ADCYAP1 | VTA | Chung 2005<sup>60</sup>, Greene 2005<sup>59</sup> |
|  | PTPRO | VM DA maturation | Xu 2022<sup>64</sup> |
| DA2<br>DANeurons-VM | TPBG | VM DA maturation | Yoo 2021<sup>42</sup> |
|  | PTGDS | VM | Zeisel 2018<sup>65</sup> |
|  | CD9 | VTA | Li 2014<sup>66</sup> |
|  | DLK1 | DA neurons | Birtele 2022<sup>67</sup> |
|  | SPARC | SN | Monzon-Sandoval 2020<sup>57</sup> |
|  | RBP1 | VM | Veenvliet 2013<sup>68</sup> |
|  | HES1 | VM | Kameda 2011<sup>69</sup>, Hegarty 2013<sup>70</sup> |
| DA3<br>DANeurons-SN | TTR | DA neurons | Kim 2021<sup>71</sup> |
|  | NEUROD1 | DA neurons | Earley 2021<sup>72</sup>, Termine 2022<sup>73</sup> |
|  | NDUFA4 | DA neurons | Fernandes 2020<sup>63</sup> |
|  | SYT1 | SN | Poulin 2014<sup>74</sup> |
|  | SLC1A2 | SN | Zhang 2020<sup>75</sup> |
|  | OLFM2 | SN | Bosser 2009<sup>76</sup>, Yang 2022<sup>77</sup> |
|  | FAM19A4 | SN | Li 2016<sup>78</sup>, Kamath 2022<sup>50</sup> |

**Table S7:** Annotation of DA subtypes from different references.

| Source | DA1 - VTA | DA2 - VM | DA3 - SN |
| --- | --- | --- | --- |
| Kamath 2022<sup>50</sup> DA subtypes of SN (Human) | SOX6_DDT | CALB1_RBP4 | CALB1_CALCR (not a good fit)<br>Should be FAM19A4 |
| Poulin 2014<sup>74</sup> DA subtypes (Mouse) | DA2B (VTA) | DA1A (SN) | Maybe DA2C<br>OTX2 and SLC17A6 (VGLUT2). |
| Poulin 2014<sup>74</sup> VTA vs SN list | Both – more VTA | Both | Both – RAB3C SN and OTX2 VTA |
| Tiklova 2019<sup>79</sup> Gene expression in shiny app (Mouse) | VT-Dat-high (doral VTA/PAG) | VT-Dat-high (doral VTA/PAG) | AT-Dat-high (SN), N-Dat-low (not matched to a region) |
| Tiklova 2019<sup>79</sup> From marker list (Mouse) | AT-Dat (SN), VT-Dat (dorsal VTA/PAG), T-Dat (VTA some SN) | AT-Dat (SN) (possible strongest match), VT-Dat (dorsal VTA/PAG), T-Dat (VTA some SN) | AT-Dat, VT-Dat, T-Dat also G-dat-low, GT-Dat-low (VTA) |
| Aguila 2022<sup>80</sup> SN vs VTA (Human) | VTA | VM | SN |
| Cluster markers (Table 13) | Ventral Midbrain possible VTA | Ventral midbrain both SN and VTA | Possible SN |
| Overall | VTA | VM | SN |

**Table S8:** Proportions of cell types and cell subtypes in four FACS sorted populations from scRNAseq transcriptomics.

| **Cell types** | **Neurons 1** | **Neurons 2** | **Astrocytes** | **Radial Glia** |
| --- | --- | --- | --- | --- |
| Astrocytes | 1% | 4% | 87% | 19% |
| DANeurons | 12% | 7% | 0% | 6% |
| Neurons | 67% | 71% | 5% | 7% |
| NPC | 8% | 0% | 0% | 0% |
| All Neuronal Cells | 87% | 79% | 5% | 13% |
| RadialGlia | 11% | 13% | 5% | 65% |
| Mix | 1% | 4% | 3% | 2% |
| **Cell subtype** | **Neurons 1** | **Neurons 2** | **Astrocytes** | **Radial Glia** |
| Astrocytes-immature | 0% | 4% | 51% | 9% |
| Astrocytes-reactive | 0% | 0% | 25% | 10% |
| Astrocytes-resting | 0% | 0% | 11% | 0% |
| DANeurons-SN | 4% | 1% | 0% | 1% |
| DANeurons-VM | 1% | 3% | 0% | 2% |
| DANeurons-VTA | 8% | 4% | 0% | 3% |
| Mix | 1% | 4% | 3% | 2% |
| Neurons_stem_potential | 14% | 0% | 0% | 0% |
| Neurons-5HT | 0% | 5% | 0% | 1% |
| Neurons-Endocrine | 0% | 39% | 0% | 3% |
| Neurons-Excitatory | 33% | 0% | 0% | 0% |
| Neurons-GABAergic | 0% | 15% | 2% | 3% |
| Neurons-Glutamatergic | 19% | 0% | 0% | 0% |
| Neurons-Mature | 0% | 12% | 2% | 1% |
| NPC | 8% | 0% | 0% | 0% |
| RadialGlia-proliferating | 1% | 0% | 1% | 2% |
| RadialGlia-NSC | 3% | 0% | 0% | 0% |
| RadialGlia-outer | 0% | 1% | 1% | 12% |
| RadialGlia-SVZ | 7% | 3% | 2% | 22% |
| RadialGlia-VZ | 1% | 9% | 2% | 29% |

**Table S9:** Antibody panel used for time course analysis with cell types previously reported to be identified by each marker.

| **Antibody/**<br>**Marker** | **Protein/**<br>**Gene** | **Reported Cell type**<br>**marker** | **References** |
| --- | --- | --- | --- |
| CD24 | CD24 | Neurons and neural stem cells<br>Cancer stem cells | Uchida 2000,<sup>25</sup> Pruszk 2007,<sup>16</sup><br>Pruszk 2009,<sup>26</sup> Sundberg 2009,<sup>27</sup><br>Yuan 2011,<sup>19</sup> Wang 2013<sup>28</sup> |
| CD56 | NCAM1 | Neurons and neural stem cells<br>Cancer cells | Pruszk 2007,<sup>16</sup> Pruszk 2009,<sup>26</sup><br>Sundberg 2009<sup>27</sup> |
| CD29 | ITGB1 | Stem cell | Pruszk<sup>16</sup>, Yuan 2011<sup>19</sup>, |
| CD15 | FUT4 | Neural precursor | Pruszk 2007,<sup>16</sup> Pruszk 2009,<sup>26</sup><br>Yuan 2011,<sup>19</sup> Sandor 2017<sup>29</sup> |
| CD184 | CXCR4 | Neural stem cell | Yuan 2011,<sup>19</sup> Sandor 2017<sup>29</sup> |
| CD133 | PROM1 | Stem cell | Uchida 2000,<sup>25</sup> Pruszk 2007,<sup>16</sup><br>Barraud 2007<sup>30</sup>, Pruszk 2009,<sup>26</sup> |
| CD44 | CD44 | Glia | Liu 2004,<sup>31</sup> Yuan 2011,<sup>19</sup> |
| CD140a | PDGFRA | OPC | Liu 2004,<sup>31</sup> Wang 2013<sup>28</sup> |
| TH | TH | Dopaminergic neurons and lineage | Wolf 1989,<sup>81</sup> Kan 2007<sup>82</sup> |
| SSEA4 | Carbohydrate epitope | Neural stem cell, stem cell | Henderson, 2002,<sup>40</sup> Barraud 2007,<sup>30</sup><br>Pruszk 2007,<sup>16</sup> Abujarour 2013<sup>83</sup> |
| CD49f | ITA6/ITGA6 | Activated astrocytes | Barbar 2020<sup>41</sup> |

**Table S10:**
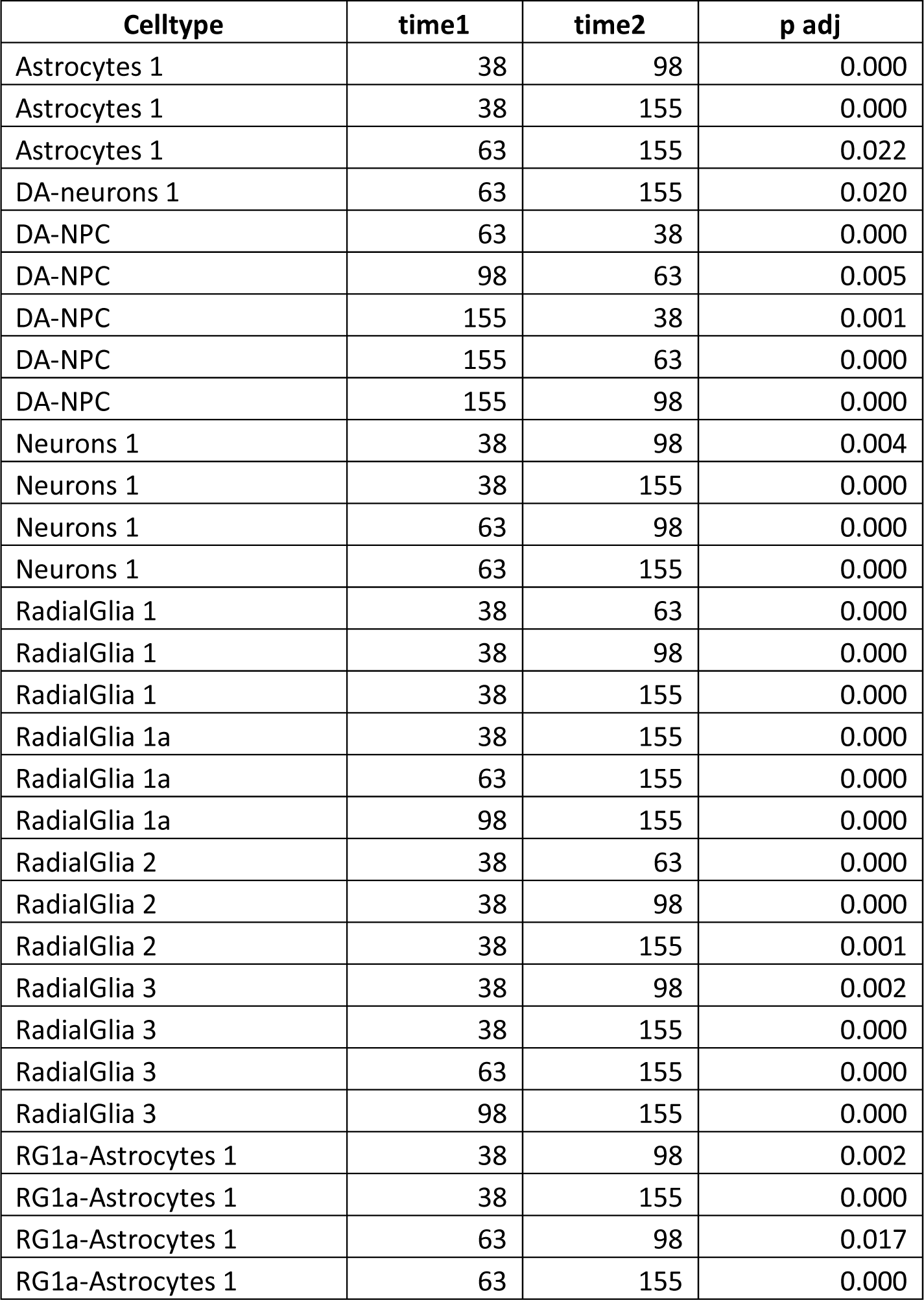
Tukey’s test results comparing each time point for each cell types. Significant differences are shown, all not listed contrasts are not significant.

## References

1. Mohamed, N.-V. et al. Midbrain organoids with an *SNCA* gene triplication model key features of synucleinopathy. Brain Commun. 3, fcab223 (2021).

2. Di Lullo, E. & Kriegstein, A. R. The use of brain organoids to investigate neural development and disease. Nat. Rev. Neurosci. 18, 573 (2017).

3. Wray, S. Modelling neurodegenerative disease using brain organoids. in Seminars in Cell & Developmental Biology vol. 111 60–66 (Elsevier, 2021).

4. Mohamed, N.-V. et al. Generation of human midbrain organoids from induced pluripotent stem cells. MNI Open Res. 3, 1 (2021).

5. Fiorenzano, A. et al. Single-cell transcriptomics captures features of human midbrain development and dopamine neuron diversity in brain organoids. Nat. Commun. 12, 1–19 (2021).

6. Andrews, T. S. & Hemberg, M. Identifying cell populations with scRNASeq. Mol. Aspects Med. 59, 114–122 (2018).

7. Chen, G., Ning, B. & Shi, T. Single-cell RNA-seq technologies and related computational data analysis. Front. Genet. 10, 317 (2019).

8. Jiang, R., Sun, T., Song, D. & Li, J. J. Statistics or biology: the zero-inflation controversy about scRNA-seq data. Genome Biol. 23, 1–24 (2022).

9. Fernández-Zapata, C., Leman, J. K., Priller, J. & Böttcher, C. The use and limitations of single-cell mass cytometry for studying human microglia function. Brain Pathol. 30, 1178–1191 (2020).

10. Nguyen, Q. H., Pervolarakis, N., Nee, K. & Kessenbrock, K. Experimental considerations for single-cell RNA sequencing approaches. Front. Cell Dev. Biol. 6, 108 (2018).

11. Bremond Martin, C., Simon Chane, C., Clouchoux, C. & Histace, A. Recent Trends and Perspectives in Cerebral Organoids Imaging and Analysis. Front. Neurosci. 15, 717 (2021).

12. Albanese, A. et al. Multiscale 3D phenotyping of human cerebral organoids. Sci. Rep. 10, 1–17 (2020).

13. Brown, M. & Wittwer, C. Flow cytometry: principles and clinical applications in hematology. Clin. Chem. 46, 1221–1229 (2000).

14. Drouet, M. & Lees, O. Clinical applications of flow cytometry in hematology and immunology. Biol. Cell 78, 73–78 (1993).

15. Woo, J., Baumann, A. & Arguello, V. Recent advancements of flow cytometry: new applications in hematology and oncology. Expert Rev. Mol. Diagn. 14, 67–81 (2014).

16. Pruszak, J., Sonntag, K.-C., Aung, M. H., Sanchez-Pernaute, R. & Isacson, O. Markers and methods for cell sorting of human embryonic stem cell-derived neural cell populations. Stem Cells 25, 2257–2268 (2007).

17. Turaç, G. et al. Combined flow cytometric analysis of surface and intracellular antigens reveals surface molecule markers of human neuropoiesis. PLoS One 8, e68519 (2013).

18. Woodard, C. M. et al. iPSC-derived dopamine neurons reveal differences between monozygotic twins discordant for Parkinson’s disease. Cell Rep. 9, 1173–1182 (2014).

19. Yuan, S. H. et al. Cell-surface marker signatures for the isolation of neural stem cells, glia and neurons derived from human pluripotent stem cells. PloS One 6, e17540 (2011).

20. Cheung, M. et al. Current trends in flow cytometry automated data analysis software. Cytometry A 99, 1007–1021 (2021).

21. Saeys, Y., Van Gassen, S. & Lambrecht, B. N. Computational flow cytometry: helping to make sense of high-dimensional immunology data. Nat. Rev. Immunol. 16, 449–462 (2016).

22. Chen, C. X.-Q. et al. A Multistep Workflow to Evaluate Newly Generated iPSCs and Their Ability to Generate Different Cell Types. Methods Protoc. 4, 50 (2021).

23. Monzel, A. S. et al. Derivation of human midbrain-specific organoids from neuroepithelial stem cells. Stem Cell Rep. 8, 1144–1154 (2017).

24. Atamian, A., Cordón-Barris, L. & Quadrato, G. Taming human brain organoids one cell at a time. in Seminars in Cell & Developmental Biology vol. 111 23–31 (Elsevier, 2021).

25. Uchida, N. et al. Direct isolation of human central nervous system stem cells. Proc. Natl. Acad. Sci. 97, 14720–14725 (2000).

26. Pruszak, J., Ludwig, W., Blak, A., Alavian, K. & Isacson, O. CD15, CD24, and CD29 define a surface biomarker code for neural lineage differentiation of stem cells. Stem Cells 27, 2928–2940 (2009).

27. Sundberg, M. et al. CD marker expression profiles of human embryonic stem cells and their neural derivatives, determined using flow-cytometric analysis, reveal a novel CD marker for exclusion of pluripotent stem cells. Stem Cell Res. 2, 113–124 (2009).

28. Wang, J., O’Bara, M. A., Pol, S. U. & Sim, F. J. CD133/CD140a-based isolation of distinct human multipotent neural progenitor cells and oligodendrocyte progenitor cells. Stem Cells Dev. 22, 2121– 2131 (2013).

29. Sandor, C. et al. Transcriptomic profiling of purified patient-derived dopamine neurons identifies convergent perturbations and therapeutics for Parkinson’s disease. Hum. Mol. Genet. 26, 552–566 (2017).

30. Barraud, P., Stott, S., Møllgård, K., Parmar, M. & Björklund, A. In vitro characterization of a human neural progenitor cell coexpressing SSEA4 and CD133. J. Neurosci. Res. 85, 250–259 (2007).

31. Liu, Y. et al. CD44 expression identifies astrocyte-restricted precursor cells. Dev. Biol. 276, 31–46 (2004).

32. Jurga, A. M., Paleczna, M., Kadluczka, J. & Kuter, K. Z. Beyond the GFAP-astrocyte protein markers in the brain. Biomolecules 11, 1361 (2021).

33. Henrik Heiland, D., et al. Tumor-associated reactive astrocytes aid the evolution of immunosuppressive environment in glioblastoma. Nat. Commun. 10, 1–12 (2019).

34. McPhie, D. L. et al. Oligodendrocyte differentiation of induced pluripotent stem cells derived from subjects with schizophrenias implicate abnormalities in development. Transl. Psychiatry 8, 1–10 (2018).

35. Chen, C. X.-Q. et al. Generation of homozygous PRKN, PINK1 and double PINK1/PRKN knockout cell lines from healthy induced pluripotent stem cells using CRISPR/Cas9 editing. Stem Cell Res. 62, 102806 (2022).

36. Soubannier, V. et al. Rapid Generation of Ventral Spinal Cord-like Astrocytes from Human iPSCs for Modeling Non-Cell Autonomous Mechanisms of Lower Motor Neuron Disease. Cells 11, 399 (2022).

37. Kwak, T. H. et al. Generation of homogeneous midbrain organoids with in vivo-like cellular composition facilitates neurotoxin-based Parkinson’s disease modeling. Stem Cells 38, 727–740 (2020).

38. Nowakowski, T. J. et al. Spatiotemporal gene expression trajectories reveal developmental hierarchies of the human cortex. Science 358, 1318–1323 (2017).

39. Becht, E. et al. Reverse-engineering flow-cytometry gating strategies for phenotypic labelling and high-performance cell sorting. Bioinformatics 35, 301–308 (2019).

40. Henderson, J. K. et al. Preimplantation human embryos and embryonic stem cells show comparable expression of stage-specific embryonic antigens. Stem Cells 20, 329–337 (2002).

41. Barbar, L. et al. CD49f is a novel marker of functional and reactive human iPSC-derived astrocytes. Neuron 107, 436–453 (2020).

42. Yoo, J.-E. et al. Trophoblast glycoprotein is a marker for efficient sorting of ventral mesencephalic dopaminergic precursors derived from human pluripotent stem cells. Npj Park. Dis. 7, 1–11 (2021).

43. Mohamed, N.-V. et al. Microfabricated disk technology: Rapid scale up in midbrain organoid generation. Methods 203, 465–477 (2022).

44. Van Gassen, S. et al. FlowSOM: Using self-organizing maps for visualization and interpretation of cytometry data. Cytometry A 87, 636–645 (2015).

45. Levine, J. H. et al. Data-driven phenotypic dissection of AML reveals progenitor-like cells that correlate with prognosis. Cell 162, 184–197 (2015).

46. Stuart, T. et al. Comprehensive integration of single-cell data. Cell 177, 1888–1902 (2019).

47. Elbaz, B. & Popko, B. Molecular control of oligodendrocyte development. Trends Neurosci. 42, 263–277 (2019).

48. Zhang, Y. et al. Purification and characterization of progenitor and mature human astrocytes reveals transcriptional and functional differences with mouse. Neuron 89, 37–53 (2016).

49. La Manno, G. et al. Molecular diversity of midbrain development in mouse, human, and stem cells. Cell 167, 566–580 (2016).

50. Kamath, T. et al. Single-cell genomic profiling of human dopamine neurons identifies a population that selectively degenerates in Parkinson’s disease. Nat. Neurosci. 25, 588–595 (2022).

51. Bhaduri, A. et al. Cell stress in cortical organoids impairs molecular subtype specification. Nature 578, 142–148 (2020).

52. van Bruggen, D. et al. Developmental landscape of human forebrain at a single-cell level identifies early waves of oligodendrogenesis. Dev. Cell 57, 1421–1436 (2022).

53. Bhaduri, A. et al. An atlas of cortical arealization identifies dynamic molecular signatures. Nature 598, 200–204 (2021).

54. Tanaka, Y., Cakir, B., Xiang, Y., Sullivan, G. J. & Park, I.-H. Synthetic analyses of single-cell transcriptomes from multiple brain organoids and fetal brain. Cell Rep. 30, 1682–1689 (2020).

55. McGinnis, C. S., Murrow, L. M. & Gartner, Z. J. DoubletFinder: doublet detection in single-cell RNA sequencing data using artificial nearest neighbors. Cell Syst. 8, 329–337 (2019).

56. Speir, M. L. et al. UCSC Cell Browser: visualize your single-cell data. Bioinformatics 37, 4578– 4580 (2021).

57. Monzón-Sandoval, J. et al. Human-Specific Transcriptome of Ventral and Dorsal Midbrain Dopamine Neurons. Ann. Neurol. 87, 853–868 (2020).

58. Wen, G. et al. Proteomic characterization of secretory granules in dopaminergic neurons indicates chromogranin/secretogranin-mediated protein processing impairment in Parkinson’s disease. Aging 13, 20335 (2021).

59. Greene, J. G., Dingledine, R. & Greenamyre, J. T. Gene expression profiling of rat midbrain dopamine neurons: implications for selective vulnerability in parkinsonism. Neurobiol. Dis. 18, 19– 31 (2005).

60. Chung, C. Y. et al. Cell type-specific gene expression of midbrain dopaminergic neurons reveals molecules involved in their vulnerability and protection. Hum. Mol. Genet. 14, 1709–1725 (2005).

61. Eshraghi, M. et al. RasGRP1 is a causal factor in the development of l-DOPA–induced dyskinesia in Parkinson’s disease. Sci. Adv. 6, eaaz7001 (2020).

62. Yin, M. et al. Ventral mesencephalon-enriched genes that regulate the development of dopaminergic neurons in vivo. J. Neurosci. 29, 5170–5182 (2009).

63. Fernandes, H. J. R. et al. Single-Cell Transcriptomics of Parkinson’s Disease Human In Vitro Models Reveals Dopamine Neuron-Specific Stress Responses. Cell Rep. 33, 108263 (2020).

64. Xu, P. et al. Human midbrain dopaminergic neuronal differentiation markers predict cell therapy outcomes in a Parkinson’s disease model. J. Clin. Invest. 132, (2022).

65. Zeisel, A. et al. Molecular architecture of the mouse nervous system. Cell 174, 999–1014 (2018).

66. Li, M. D., Burns, T. C., Morgan, A. A. & Khatri, P. Integrated multi-cohort transcriptional meta-analysis of neurodegenerative diseases. Acta Neuropathol. Commun. 2, 1–23 (2014).

67. Birtele, M. et al. Single-cell transcriptional and functional analysis of dopaminergic neurons in organoid-like cultures derived from human fetal midbrain. Development 149, dev200504 (2022).

68. Veenvliet, J. V. et al. Specification of dopaminergic subsets involves interplay of En1 and Pitx3. Development 140, 3373–3384 (2013).

69. Kameda, Y., Saitoh, T. & Fujimura, T. Hes1 regulates the number and anterior–posterior patterning of mesencephalic dopaminergic neurons at the mid/hindbrain boundary (isthmus). Dev. Biol. 358, 91–101 (2011).

70. Hegarty, S. V., Sullivan, A. M. & O’keeffe, G. W. Midbrain dopaminergic neurons: a review of the molecular circuitry that regulates their development. Dev. Biol. 379, 123–138 (2013).

71. Kim, T. W. et al. Biphasic activation of WNT signaling facilitates the derivation of midbrain dopamine neurons from hESCs for translational use. Cell Stem Cell 28, 343–355 (2021).

72. Earley, A. M., Burbulla, L. F., Krainc, D. & Awatramani, R. Identification of ASCL1 as a determinant for human iPSC-derived dopaminergic neurons. Sci. Rep. 11, 1–13 (2021).

73. Termine, A. et al. A Hybrid Machine Learning and Network Analysis Approach Reveals Two Parkinson’s Disease Subtypes from 115 RNA-Seq Post-Mortem Brain Samples. Int. J. Mol. Sci. 23, 2557 (2022).

74. Poulin, J.-F. et al. Defining midbrain dopaminergic neuron diversity by single-cell gene expression profiling. Cell Rep. 9, 930–943 (2014).

75. Zhang, Y. et al. Generation of a novel mouse model of Parkinson’s disease via targeted knockdown of glutamate transporter GLT-1 in the substantia nigra. ACS Chem. Neurosci. 11, 406–417 (2020).

76. Bossers, K. et al. Analysis of gene expression in Parkinson’s disease: possible involvement of neurotrophic support and axon guidance in dopaminergic cell death. Brain Pathol. 19, 91–107 (2009).

77. Yang, Y., Huang, X., Wang, C. & Wang, Y. Identification of hub genes of Parkinson’s disease through bioinformatics analysis. Front. Neurosci. 1709 (2022).

78. Li, C.-L. et al. Somatosensory neuron types identified by high-coverage single-cell RNA- sequencing and functional heterogeneity. Cell Res. 26, 83–102 (2016).

79. Tiklová, K. et al. Single-cell RNA sequencing reveals midbrain dopamine neuron diversity emerging during mouse brain development. Nat. Commun. 10, 1–12 (2019).

80. Aguila, J. et al. Spatial RNA sequencing identifies robust markers of vulnerable and resistant human midbrain dopamine neurons and their expression in Parkinson’s disease. Front. Mol. Neurosci. 14, 699562 (2021).

81. Wolf, M. E., Zigmond, M. J. & Kapatos, G. Tyrosine hydroxylase content of residual striatal dopamine nerve terminals following 6-hydroxydopamine administration: a flow cytometric study. J. Neurochem. 53, 879–885 (1989).

82. Kan, I. et al. Dopaminergic differentiation of human mesenchymal stem cells—utilization of bioassay for tyrosine hydroxylase expression. Neurosci. Lett. 419, 28–33 (2007).

83. Abujarour, R. et al. Optimized surface markers for the prospective isolation of high-quality hiPSCs using flow cytometry selection. Sci. Rep. 3, 1179 (2013).

